# Perturbations in fumarate levels in *Plasmodium berghei* leads to cysteine succination and impairs ookinete formation

**DOI:** 10.64898/2026.03.16.711975

**Authors:** Anusha Chandrashekarmath, Arpitha Suryavanshi, Chitralekha Sen Roy, Hemalatha Balaram

## Abstract

The TCA cycle intermediates comprise 8 carboxylic acids of which only fumarate has unsaturated carbons rendering it capable of electrophilic addition to cysteine thiols. Accumulation of fumarate due to loss of function of fumarate hydratase converts it into a powerful driver of cancer and therefore is classified as an oncometabolite. We have examined the consequences of perturbing the metabolism of fumarate and its product, malate, on *Plasmodium berghei* development across erythrocytic and early insect stages. Our studies on *P. berghei* lines lacking the genes *fh* and *mqo*, coding for the enzymes fumarate hydratase (FH) and malate quinone oxidoreductase (MQO), respectively as well as *dtc* and *ogc* coding for the transporters dicarboxylate‐tricarboxylate carrier (DTC) and citrate‐oxoglutarate carrier (OGC), show dramatic impairment in ookinete formation, while gametocytes and erythrocytic asexual stages remain largely unaffected. Comparative metabolomic analysis of gametocytes revealed that elevated fumarate levels in the knockouts led to succination of glutathione, possibly resulting in oxidative stress. The increased levels of the M+5 isotopologue of inosine monophosphate in the knockout gametocytes, observed in isotope tracer experiments, suggest enhanced ribose‐5‐phosphate and NADPH production through the pentose phosphate pathway, with the latter potentially mitigating elevated oxidative stress. The isotope tracer studies also informed that the *P. berghei* DTC is a transporter of malate and fumarate and OGC, a transporter of fumarate. The impaired ookinete formation arising from cysteine succination underscores the potential for developing transmission‐blocking agents through selective inhibition of class I parasite FH which is distinct from its class II human counterpart.

**Significance statement:** Metabolic requirements of *Plasmodium* that lives across two hosts, traversing developmental stages with different rates of growth, are highly varied. Previous studies have established the non‐essentiality of the TCA cycle genes for the erythrocytic asexual stages. Our studies show that knockout of genes involved in fumarate and malate metabolism, and transporters involved in anaplerosis, impairs development of the mosquito‐stage, ookinete, with the gametocytes of the knockout lines exhibiting elevated levels of fumarate. The accumulation of succinated glutathione, formed via Michael addition of fumarate to the cysteinyl thiol group and leading to oxidative stress, is probably the leading cause of impairment in ookinete development.

## Introduction

Malaria, caused by *Plasmodium spp*., remains a burden in 83 endemic countries with 263 million estimated cases and 597,000 deaths reported in 2023 (1). Given the parasite’s remarkable ability to develop drug resistance (2–6), new intervention strategies are continuously warranted. During the asexual blood stages, despite heavy reliance on glycolysis for ATP generation (7), the expression of the tricarboxylic acid (TCA) cycle genes at the transcript level (8) and the functional operation of the cycle have been established (9, 10). Two enzymes of the TCA cycle, namely fumarate hydratase (FH) which harbors a 4Fe‐4S cluster (11) and malate quinone oxidoreductase (MQO) that is of bacterial origin (12) are distinct from their human counterparts. The flux through the TCA cycle and the parasite’s dependency on the pathway vary across the different developmental stages as the parasite differentiates from asexual blood stages to gametocytes (9) and subsequently, to gametes and ookinetes (13).

The first study to examine the essentiality of the TCA cycle enzymes in *P. falciparum (Pf)*, showed that, except for FH and MQO, the other six enzymes are not essential during the asexual stages (14). FH catalyzes the seventh step in the TCA cycle, converting fumarate to malate. The first successful deletion of the *fh* gene in *P. berghei (Pb)* was reported by Niikura and coworkers (15), followed by studies that showed mouse strain‐specific essentiality of the gene (11). Seven years after the initial report of the essentiality of *fh* in *Pf*, Rajaram and coworkers, after numerous attempts, generated viable *Pf* asexual stages lacking the *fh* gene, suggesting cell‐to‐cell metabolic variability governing the essentiality of the gene (16).

*Plasmodium* MQO catalyzes the oxidation of malate to oxaloacetate (OAA) using ubiquinone as a cofactor, which in turn transfers electrons to Complex III of the electron transport chain (ETC), the *in vivo* activity of which has been validated through inhibition by atovaquone (17). *Plasmodium* also harbours a cytosolic malate dehydrogenase (MDH), that catalyzes the reduction of OAA to malate with the concomitant generation of NAD^+^ (18–21), with the reverse reaction being highly unfavourable (18). The presence of mitochondrial MQO and cytosolic MDH, along with metabolite transporters, suggests the possible operation of a malate‐OAA shuttle regulating cytosolic NAD^+^ and mitochondrial ubiquinol levels. Knockout of *Pbmdh* was not lethal to the asexual erythrocytic stages (22) and MQO was also found to be non‐essential in both *Pf* and *Pb* (15, 16).

Apart from pyruvate entry, flux through the TCA cycle is mediated by anaplerotic pathways. The asexual blood stages of both *Pf* and *Pb* show a large flux in the anaplerotic entry of glutamine‐derived α‐ketoglutarate (α‐KG) into the TCA cycle (9, 13). In *Plasmodium*, fumarate is also generated in the cytosol as a byproduct of the adenylosuccinate lyase (ASL) reaction, and studies in *Pf* have established its anaplerotic entry into the TCA cycle (17). Additional support for the anaplerosis of fumarate and also malate comes from the dependence on these two metabolites for the knockout of phospho*enol*pyruvate carboxylase (*pepc*) in *Pf* (23). PEPC catalyzes the conversion of phospho*enol*pyruvate (PEP) to OAA in the cytosol. The knockout of *pepc* being conditional to external supplementation with fumarate and malate indicate entry of these two metabolites into the mitochondrion and anaplerotically feeding into the TCA cycle, and also, that OAA is exported from the mitochondrion into the cytosol (23). Therefore, the anaplerotic entry of fumarate, malate, and α‐KG must be facilitated by transporters on the mitochondrial inner membrane, with the two likely candidates being dicarboxylate‐tricarboxylate carrier (DTC; PBANKA_0706700) and citrate‐oxoglutarate carrier (OGC, PBANKA_1438700, a homolog of YHM2 (24)). Proteoliposomal studies on *Pf*DTC have shown that it mediates oxoglutarate/oxoglutarate homoexchange as well as oxaloacetate/oxoglutarate and malate/oxoglutarate counter exchanges (25) with the transporter being localized to the mitochondrion (26). Knockout of *Pfdtc* and *Pfogc* did not show growth defects in the asexual blood stages (27) with ^13^C‐glutamine tracer studies suggesting their involvement in the transport of only α‐KG into the mitochondrion (27), and not fumarate and malate.

The study presented here, reports the complete impairment of gametocyte to ookinete conversion upon knockout of *fh, mqo, dtc, ogc* and in the double knockouts *mqo/mdh* (*mm*) and *dtc/ogc* (*od*) in *Pb*. A unique observation from comparative metabolomics is the elevated oxidative stress in the knockout lines, arising from a substantial increase in the levels of succinic‐glutathione (succ‐GSH), a consequence of elevated fumarate levels. Isotope tracer studies suggest cellular attempts at compensation of oxidative stress through enhanced levels of NADPH generated via the pentose phosphate pathway (PPP), enabled by increased IMP production. Consistent with elevated fumarate, enhanced levels of succinated thiols were seen in seven proteins in the *fh* knockout. Our studies suggest the possibility of exploiting the unique features of parasite FH for development of transmission‐blocking drugs. Contrary to observations in *Pf* (27), our studies demonstrate that DTC is a transporter of malate and, to a lesser extent fumarate, whereas OGC is a transporter of fumarate.

## Results

Figure 1A shows a schematic of the metabolic pathways relevant to this study. *Pb Δfh, Δmdh, Δmqo, Δmm, Δdtc, Δogc*, and *Δod* lines were generated and phenotypically examined across erythrocytic asexual and gametocyte stages, as well as the mosquito ookinete stage. To understand the metabolic consequences of these gene knockouts, metabolomic analyses were performed on *Δfh, Δmm, Δdtc, Δogc*, and *Δod* gametocytes. As endogenous tagging of the *ogc* gene was unsuccessful, episomal expression of OGC‐GFP in *Pb* showed mitochondrial localization (Fig. S1), consistent with the localization seen in *Pf* (27).

**Figure 1.**
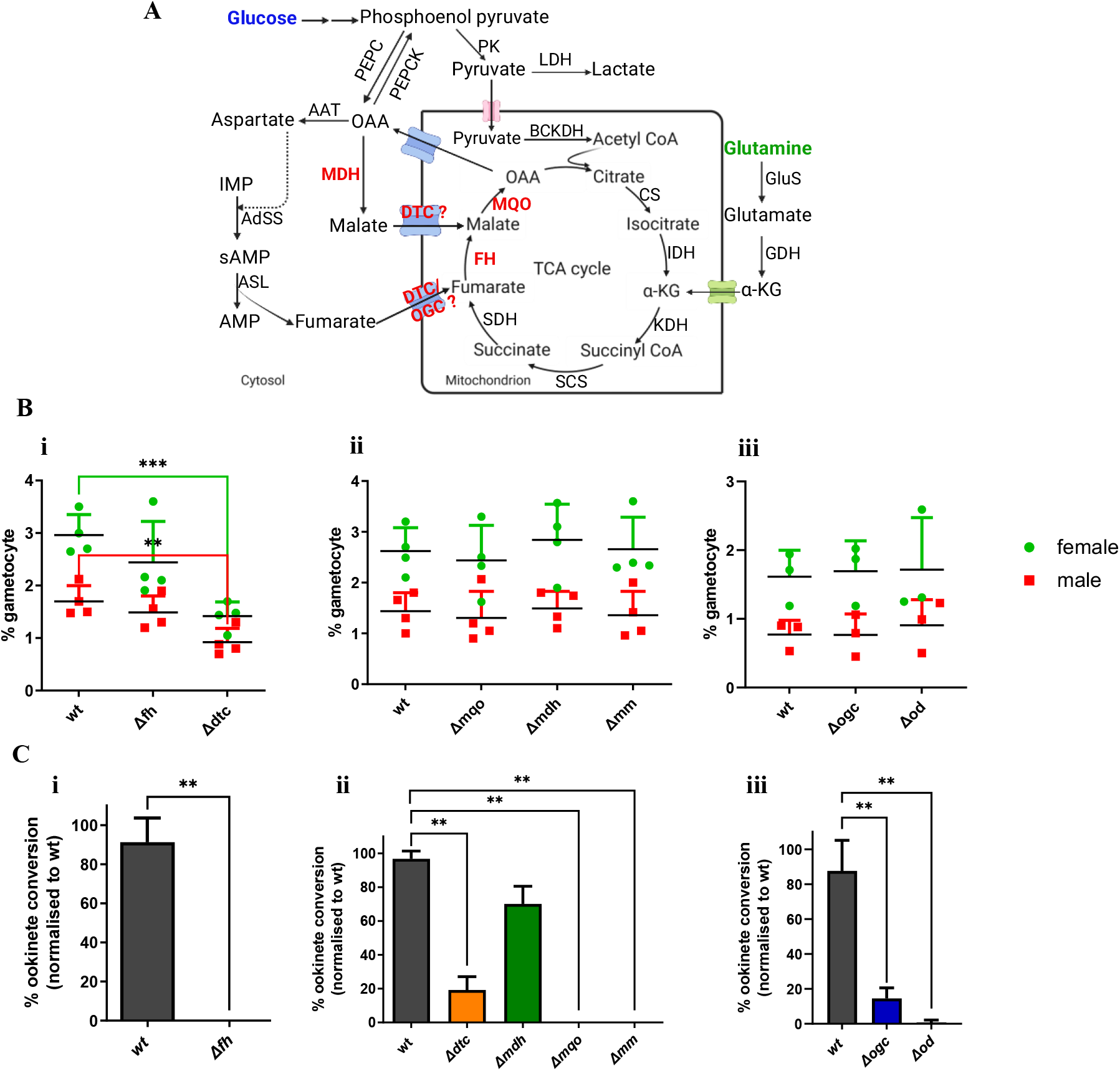
Schematic of metabolic pathways involved in the study and growth phenotype of wild‐type (WT) and knockout lines across different developmental stages. (A) The schematic shows the canonical entry of glucose‐derived pyruvate entry into the TCA cycle and various routes of anaplerosis. The knockouts under study, FH, MDH, MQO, DTC, and OGC are labelled in red. FH, fumarate hydratase; MDH, malate dehydrogenase; MQO, malate‐quinone oxidoreductase; DTC, dicarboxylate-tricarboxylate carrier; OGC, citrate-oxoglutarate carrier; PK, pyruvate kinase; LDH, lactate dehydrogenase; BCKDH, branched chain ketoacid dehydrogenase; CS, citrate synthase; IDH, isocitrate dehydrogenase; KDH, ketoglutarate dehydrogenase; SCS, succinyl‐CoA synthetase; SDH, succinate dehydrogenase; AAT, aspartate aminotransferase; AdSS, adenylosuccinate synthetase; ASL, adenylosuccinate lyase; GluS, glutamate synthase; GDH, glutamate dehydrogenase. (B) Comparison of percent female and male gametocyte between WT and the seven knockouts. Female and male gametocytemia of (i) *Δfh* and *Δdtc*, (ii) *Δmdh, Δmqo*, and *Δmm*, and (iii) *Δogc* and *Δod* plotted with their respective WT controls. The mean of percent gametocytes is plotted with error bars representing SD. Statistical analysis using Student’s unpaired t‐test did not show any significant difference in the gametocytemia between WT and the knockouts except for *Δdtc*. N = 3‐4 for WT and knockouts. (C) Comparison of *in vitro* ookinete formation between WT and the knockout lines. The percent ookinete conversion for (i) *Δfh*, (ii) *Δdtc, Δmdh, Δmqo*, and *Δmm*, and (iii) *Δogc* and *Δod* is plotted with their respective WT controls. The mean of percent ookinete formation is plotted with error bars representing SD. N=2 for wildtype, N=2‐3 for knockouts. Statistical analysis was carried out using Student’s unpaired t‐test; p<0.05 (*), p<0.005 (**), p<0.001(***). All the statistical analysis and plotting of the graphs were carried out using GraphPad Prism v9.

### Absence of *fh, mqo, dtc*, and *ogc* impairs ookinete formation

Generation of *Δfh* parasites has been described previously (11). Genetic ablation of *mqo, mdh, mqo/mdh, dtc, ogc*, and *dtc/ogc* genes in *Pb* was carried out in C57BL/6 mice. Genotyping of drug‐resistant clonal lines by PCR confirmed right integration of the selection markers along with the absence of the native gene locus (Fig. S2‐S6). The various gene knockouts and wild‐type (WT) parasites were assessed for asexual erythrocytic growth rate, mouse mortality, gametocyte formation, and *in vitro* ookinete conversion.

Comparison of the intra‐erythrocytic replication rate across *Pb* WT and *Δfh, Δmqo, Δmdh, Δmm, Δdtc, Δogc*, and *Δod* parasites in mice, as well as the mortality rate of infected mice, showed no significant differences (Fig. S7). Among the seven knockouts, only *Δdtc* showed a slight drop in the number of gametocytes formed, with the male to female ratio remaining unperturbed (Fig. 1B, i‐iii). In *in vitro* ookinete conversion assays, except for *Δmdh*, the rest of the gene knockout parasites were severely impaired in ookinete formation (Fig. 1C). Although *Δdtc* and *Δogc* parasites showed a small percentage converting to mature ookinetes, *Δod* parasites were almost completely defective in the development of gametocytes to mature ookinetes (Fig. 1C, ii‐iii). Notably, *Δfh, Δmqo*, and *Δmm* parasites exhibited a complete absence of mature ookinetes (Fig. 1C, i‐ii).

### Enhanced fumarate levels lead to succination of cysteine and cysteinyl residues

Gametocytes of *Pb* were chosen for metabolomics studies as the knockouts under investigation showed defects only in the development of the mosquito ookinete stages. The knockouts examined were *Δfh, Δmm, Δdtc, Δogc*, and *Δod* and the heat map representing log_2_ fold‐change (knockout/wt) for 23 metabolites is shown in Figure 2A. In all the five knockouts, an increase in the levels of fumarate was observed, leading to increased levels of 2‐(succino)cysteine (S‐cysteinosuccinic acid, 2SC) and succ‐GSH (Fig. 2A) arising from the Michael addition of fumarate with thiols, resulting in covalent adduct formation (28). The MS/MS spectra of 2SC and succ‐GSH validated their identification (Fig. S8A and B). Significant accumulation of malate was observed in *Δmm, Δdtc*, and *Δod*, and to a much lower extent in *Δfh*. A higher increase in malate levels in *Δdtc*, along with only a minor increase in levels of 2SC and succ‐GSH, suggests that the primary metabolite being transported by DTC is malate, with a lower affinity for fumarate. In contrast, the absence of a difference in malate levels, together with 2SC and succ‐GSH showing a large increase in *Δogc*, suggests that OGC functions as a fumarate transporter. Surprisingly, all the five knockouts displayed lower levels of IMP compared to WT. As a control, metabolite extract from uninfected RBC (uRBC) was subject to identical analysis and was found not to contribute significantly to the levels of the metabolites examined (Table S1).

**Figure 2.**
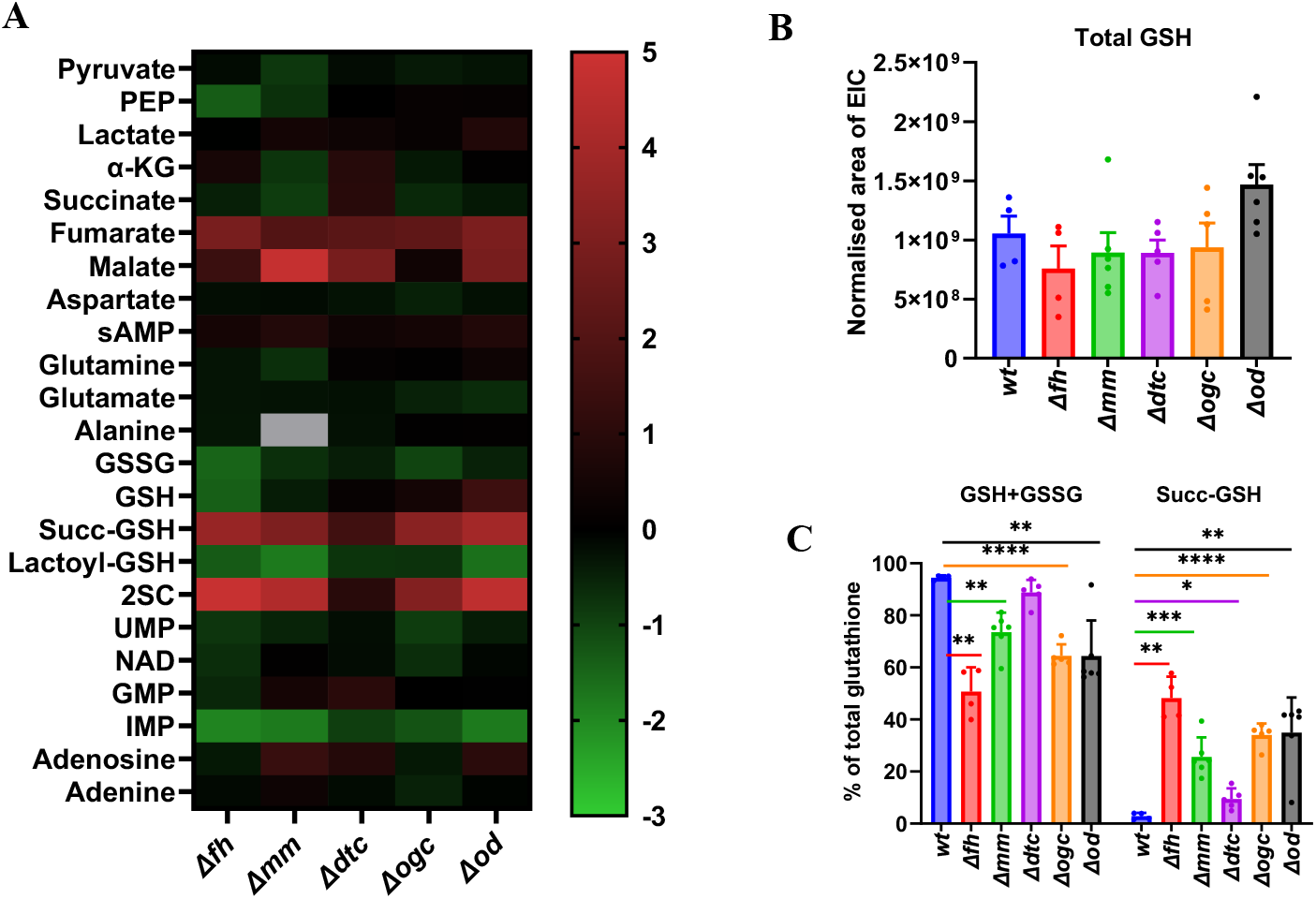
Semi‐targeted metabolomic analysis between WT and five knockouts. (A) A heat map showing log_2_ fold change (FC) in levels of metabolites between *P. berghei Δfh, Δmm, Δdtc, Δogc, Δod* and WT gametocytes. FC is the change in abundance of a metabolite computed as the ratio of knockout/WT. The area of the extracted ion chromatogram (EIC) normalised to cell numbers was used as an indicator of the amount of a metabolite, and this was computed for all 23 metabolites across WT and knockouts. The log_2_FC value is represented as a colour gradient as shown in the bar on the right, with the colours indicating the magnitude of FC. As EIC of alanine was not consistent across runs in *Δmm*, it was not considered for relative quantification and hence, shown in grey colour. (B) Total levels of glutathione estimated as a sum of the EIC areas of GSH, GSSG, lactoyl‐GSH, and succinic‐GSH did not differ across WT and knockout gametocytes. Mean values with error bars representing SEM are plotted. (C) Percent of unmodified GSH (reduced and oxidized forms) and succinic‐GSH (succ‐GSH) in WT and knockout gametocytes. Mean values with error bars representing SD were plotted. N=2 with two or three technical replicates each for WT and knockouts. Unpaired two‐tailed t‐tests were carried out using GraphPad Prism v9; p<0.05 (*), p<0.005 (**), p<0.001(***), p<0.0001 (****).

The *Δfh, Δmm*, and *Δod* gametocytes showed lowered levels of lactoyl‐GSH, with *Δfh* also displaying a reduction in the levels of reduced glutathione (GSH) and oxidized GSH (GSSG, Fig. S8C) (Fig. 2A). However, the levels of total glutathione estimated as the sum of reduced and oxidised forms, along with lactoyl‐GSH and succ‐GSH remained the same across WT and all the five knockout parasites (Fig. 2B). The percent succ‐GSH was highest in *Δfh* (48.2%), followed by *Δogc* and *Δod*, showing similar levels of 34‐35%, and *Δmm* and *Δdtc* showing 25.6% and 9.5%, respectively (Fig. 2C).

As we observed increase in levels of succinated cysteine and glutathione in *Δfh* gametocytes, an increase in the succination of cysteinyl residues in specific proteins was expected. Western blot analysis of gametocyte lysates from WT and *Δfh*, probed using anti‐2SC antibody, revealed the presence of a ∼45 KDa band harboring the 2SC modification that was present only in the knockout and absent in the WT lysate (Fig. S9). MS/MS analysis of in‐gel tryptic peptides extracted from *Δfh* gametocytes identified seven proteins containing a single 2SC modification that was absent in the WT (Table 1, Figure S10‐S16). Among these, RNA helicase (45.3 KDa), tRNA import protein (46.1 KDa), and chromatin assembly factor 1 protein WD40 domain (50.5 KDa) fall within the 45±5 KDa range seen in the Western blot and have percent 2SC modification levels of 38%, 49%, and 61%, respectively. These results show that fumarate accumulation also leads to succination of cysteinyl residues in proteins.

**Table 1.**
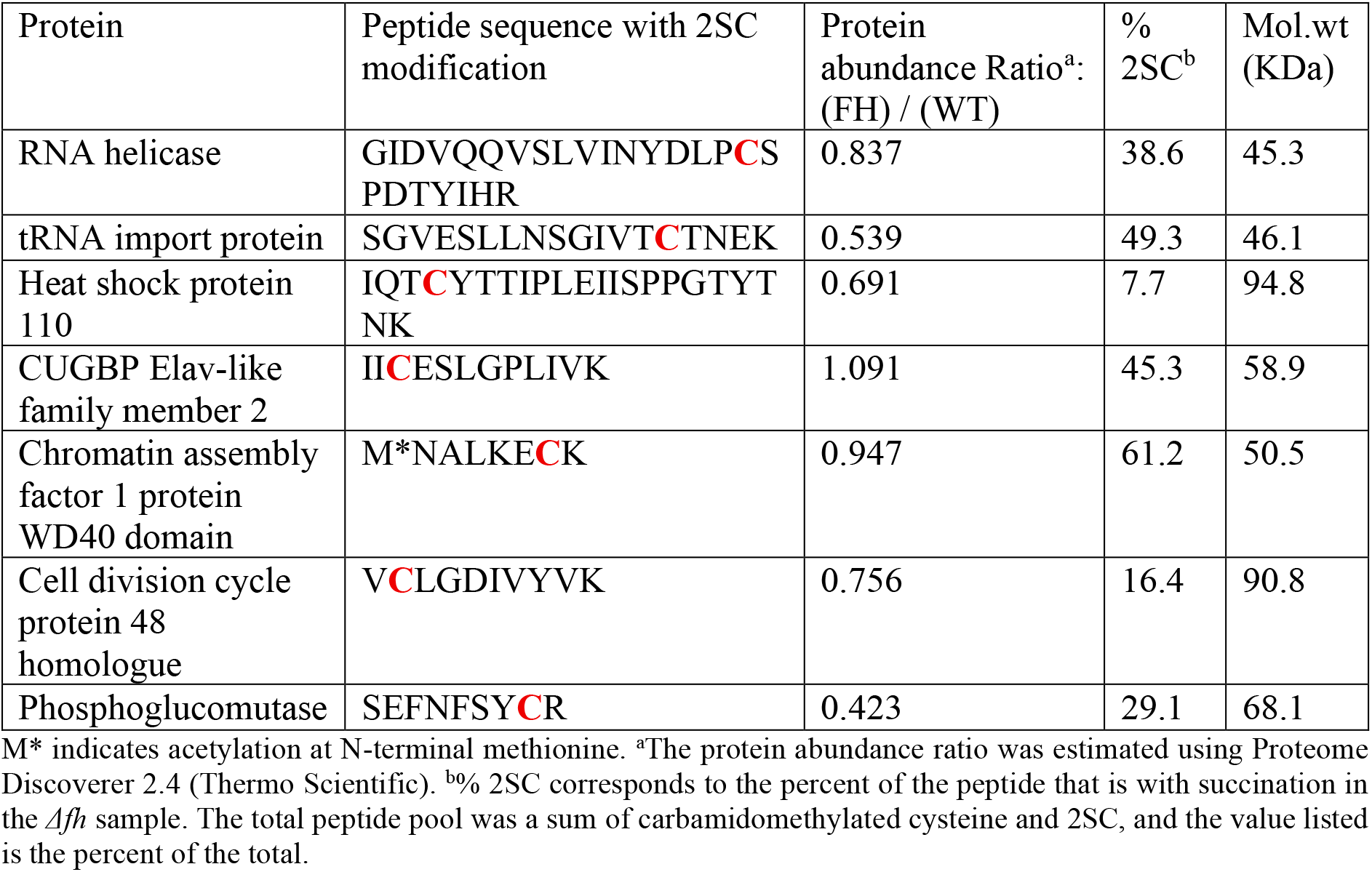
Proteins with 2SC modification. The seven proteins and their respective peptides harboring the modified cysteine are listed along with their abundance, percent modification, and molecular weight (Mol. wt). The cysteinyl residue with 2SC modification is indicated in red.

### Knockouts display an alteration in glucose‐derived TCA cycle

The stable isotope tracers, ^13^C_6_‐glucose and ^13^C_5_^15^N_2_‐glutamine were used to follow alterations in isotopologues levels across WT and *Δfh, Δmm, Δdtc, Δogc*, and *Δod* gametocytes. Analysis of the MS data after natural abundance correction was done in two ways, *viz*., percent isotopologue distribution and extracted ion chromatogram (EIC) areas, as both these representations provide useful insights (29). The expected labelling patterns in TCA cycle intermediates arising from ^13^C_6_‐glucose and ^13^C_5_^15^N_2_‐glutamine (30) are shown in the Figure S17.

The glycolytic intermediates PEP and lactate, along with the TCA cycle intermediates α‐KG, succinate, fumarate and malate, were examined for ^13^C_6_‐glucose**‐**derived carbon incorporation. A significant proportion of PEP and lactate was M+3 labelled in the knockouts, indicating an active glycolytic pathway (Fig. S18A and B). Further, an increase in EIC area of M+3 lactate was observed in *Δmm* and *Δod* (Fig. S18C), suggesting an increased level of NAD^+^ production through the LDH reaction. The observed labelling in α‐KG, succinate, fumarate and malate suggest that glucose‐derived pyruvate entry into the TCA cycle decreases significantly in *Δfh, Δmm*, and *Δod* gametocytes (Fig. 3A‐B and S18F‐G).

**Figure 3.**
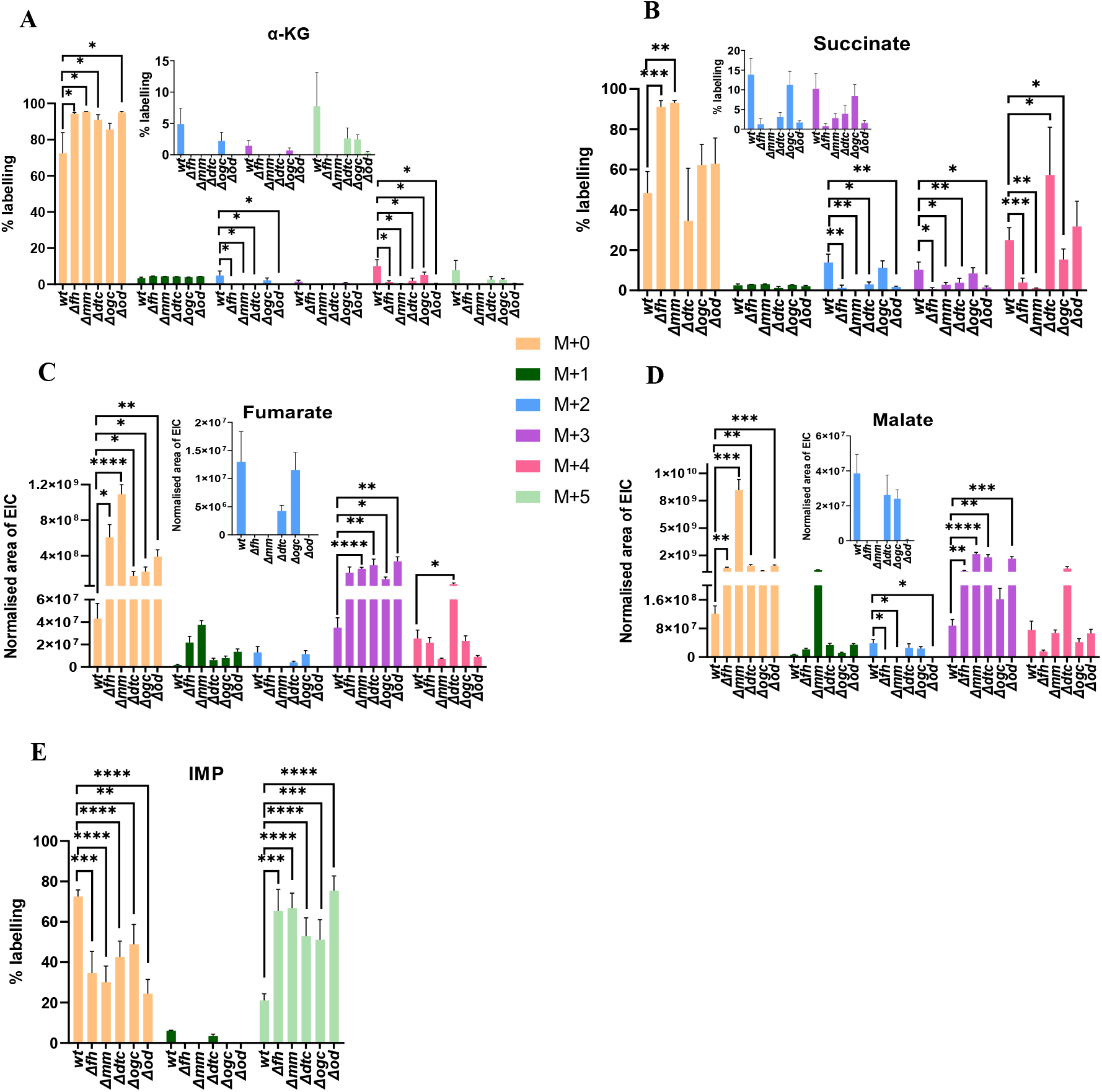
Incorporation of ^13^C_6_‐glucose‐derived carbon into the TCA cycle intermediates and IMP. The bar graphs represent the percent enrichment in the isotopologues of (A) α‐KG, (B) succinate, and (E) IMP from WT and knockout gametocytes. Mean values with error bars representing SD were plotted. The EIC area of the various isotopologues of (C) fumarate and (D) malate from WT and knockout gametocytes. Mean values with error bars representing SEM were plotted. The insets in the panels (A) ‐ (D) show the zoomed‐in histograms for some of the isotopologues. N=2 for WT, *Δfh, Δogc*, and *Δod* and N=3 for *Δmm* and *Δdtc* each with two or three technical replicates. Unpaired two‐tailed t‐tests were carried out using GraphPad Prism v9; p<0.05 (*), p<0.005 (**), p<0.001 (***), p<0.0001(****).

The overall labelling of α‐KG from ^13^C_6_‐glucose in WT was low due to its simultaneous generation from unlabeled glutamine (Fig. 3A). While the combined percent isotopologues (+2 to +5 combined) of α‐KG was 24.2±12.0% in the WT, this dropped to 1.3±0.7%, 0.2±0.1%, 4.7±2.8%, 0.6±0.4% in *Δfh, Δmm, Δdtc*, and *Δod*, respectively. This was also reflected in the total percent labelling of succinate (+2 to +4 combined), which decreased from 49±11% in WT to 6.0±3.2% and 3.8±1.2% in *Δfh* and *Δmm*, respectively (Fig. 3B). The difference in total percent labelling across α‐KG and succinate in WT is likely a consequence of a significant fraction of the former metabolite being retained in the cytosol. Unexpectedly, the percent label and EIC area of M+4 succinate increased in *Δdtc* and remained largely unchanged in *Δod* (Fig. 3B and S18E), despite the lowered levels of M+4 and M+5 α‐KG (Fig. S18D). In contrast, the percent label of M+2 succinate decreased in both *Δdtc* and *Δod*. The *Δogc* gametocytes showed only a mild reduction in TCA cycle‐derived isotopologues of α‐KG, succinate, and malate (Fig. 3A‐B and S18G).

M+2 labelled fumarate and malate arise exclusively from the entry of M+3 pyruvate into the TCA cycle, whereas M+4 can be contributed by both the TCA cycle and the anaplerotic entry of M+3 malate, fumarate, or OAA generated by MDH, ASL, and PEPC reactions in the cytosol. M+3 fumarate and malate can also be produced via the TCA cycle (Fig. S17B). In *Δfh, Δmm*, and *Δod* gametocytes, there was no active synthesis of fumarate through the TCA cycle, as M+2 levels were below the detection limit (Fig. 3C and S18F). The EIC area of M+3 fumarate was 6‐fold higher in *Δfh* and 7.2‐fold higher in *Δmm* (Fig. 3C), indicating the accumulation of fumarate generated by the ASL reaction that is not further metabolised either due to the absence of FH or MQO in the parasite. In *Δdtc, Δogc*, and *Δod* gametocytes, an increase in EIC area of M+3 fumarate by 8.3, 3.9, and 9.6‐fold (Fig. 3C), respectively indicates that both transporters contribute to fumarate uptake into the mitochondrion, with the impairment in uptake enhanced in the double knockout.

M+2 malate was below the limits of detection in *Δfh, Δmm*, and *Δod* gametocytes (Fig. 3D and S18G). In *Δfh* gametocytes, the 3.4‐fold increase in EIC area of M+3 malate (Fig. 3D), despite the presence of MQO indicates that due to reduced TCA cycle flux, the MDH generated malate accumulates. In *Δmm* gametocytes, the EIC area of M+3 malate was 24.4‐fold higher (Fig. 3D). Due to the absence of MDH in *Δmm* gametocytes, M+3 malate is generated exclusively via the FH reaction. The high levels of fumarate arising from the ASL reaction in *Δmm* gametocytes, contribute to M+3 malate accumulation through FH‐mediated conversion of anaplerotic fumarate. In *Δdtc* gametocytes, M+3 malate showed 20.4‐fold higher EIC area (Fig. 3D), which was also evident in enhanced labelling in the M+3 population (55.1±8.2%) (Fig. S18G), indicating that the cytosolic malate generated by MDH accumulates due to compromised mitochondrial import. This clearly demonstrates that apart from fumarate, DTC transports malate also. Although mild changes in percent labelling of malate were observed in *Δogc*, the EIC areas of the isotopologues were similar to WT. An increase in M+3 percent labelling of malate to 62.1±3.6% (Fig. S18G) and EIC area of this isotopologue by 18.8‐fold in *Δod* (Fig. 3D) was similar to that observed in *Δdtc*. These data indicate that the increase in malate levels arises primarily from disruption of *dtc* rather than *ogc*, confirming that DTC, and not OGC, mediates transport of malate.

In addition, the presence of M+2 and M+4 aspartate in WT gametocytes, which arise from TCA cycle‐derived OAA, confirmed the export of mitochondrial OAA into the cytosol (Fig. S19A). A decrease in EIC area of M+2 and M+4 aspartate by 14.1 and 4.2‐fold, respectively, in *Δdtc* (Fig. S19B), and M+4 aspartate by 21‐fold in *Δod*, indicates that OAA export from the mitochondrion is impaired in these two knockouts. These results suggest that DTC could also have substrate specificity for OAA, as observed in *in vitro* proteoliposomal studies (25). However, despite the decrease in EIC area of M+4 aspartate, we are unable to provide a mechanistic explanation for the observed large increase in M+4 succinate, fumarate, and malate in *Δdtc* (Fig. S18E, 3C and D). The labelling in aspartate from glucose tracer has also been reported previously in *Pf* gametocytes (9).

Analysis of ^13^C_6_‐glucose‐fed uRBC showed that α‐KG, succinate, fumarate, malate, and aspartate remained largely unlabelled (M+0 > 90%) (Fig. S20A‐E) indicating that the observed labelling in *P. berghei* gametocytes arises predominantly from parasite metabolism.

### Increased IMP synthesis upregulates NADPH production via PPP in the knockouts

The isotopologues of the nucleotides IMP, GMP, succinyl‐AMP (sAMP), NAD, and UMP were also examined. In WT gametocytes, IMP was not highly labelled, with M+0 population at 72.5±3.2% and the M+5 population at a lower level of 21.0±3.3% (Fig. 3E). The M+5 population arises from the +5 isotopologue of phosphoribosyl pyrophosphate (PRPP), which phosphoribosylates hypoxanthine to form M+5 IMP. PRPP is synthesized from ribose‐5‐phosphate, an intermediate of the PPP. In all the five knockout parasites, the percent labelling of M+5 IMP increased drastically, ranging from 51.1 to 75.4% (Fig. 3E), indicating a higher level of IMP synthesis compared to WT. In addition, the ratio of M+5 to M+0 EIC areas was also higher in the knockouts, further supporting increased level of IMP synthesis in the mutants (Fig. 21B). As IMP is the precursor for sAMP synthesis, the knockouts also showed moderately increased labelling of (M+5)+(M+8) sAMP (Fig. 21C). This apart, we observe an active utilization of M+5 IMP in the generation of M+5 GMP and M+5 NAD (Fig. 21D and E). In addition, the absence of an increase in EIC area of M+5 IMP, along with the large drop in the EIC area of the M+0 isotopologue (Fig. 21A) and its reduced levels in comparative metabolomics (Fig. 2A), supports its active consumption for the synthesis of downstream metabolites. Taken together, the increase in IMP synthesis in the knockouts, requiring increased supply of PRPP, could have triggered an increase in the levels of NADPH through the PPP. It should be noted that PRPP is generated from ribose‐5‐phosphate, an intermediate in PPP. In contrast, UMP, a pyrimidine nucleotide remained largely unlabelled in both WT and the five knockout gametocytes (Fig. S21F).

### Substrate specificity of DTC and OGC confirmed by glutamine tracer studies

Upon using ^13^C_5_ ^15^N_2_ ‐glutamine as the tracer, glutamate in all the five knockout lines showed high levels of M+5, N+1 isotopologue, indicating active synthesis from glutamine (Fig. S22A). Labelled glutamate is converted by either glutamate dehydrogenase (GDH) or aspartate aminotransferase (AAT) to M+5 α‐KG that feeds anaplerotically into the TCA cycle. The labelling patterns of TCA cycle intermediates arising from ^13^C_5_ ^15^N_2_ ‐glutamine are shown in Figure S17D and the percent isotopologue distribution is shown in Figure S22B‐E. In *Δfh* and *Δmm*, the reduction in EIC area of M+3 α‐KG and M+2 succinate (Fig. 4A and B), arising from M+5 α‐KG completing the first round of TCA cycle, indicates a blockage of the TCA cycle due to absence of FH and MQO. As a consequence, a decrease in EIC area of M+2 fumarate and malate (Fig. 4C and D) was observed. In contrast, there was an increase in EIC areas of M+4 fumarate (8.2‐fold in *Δfh* and 4.6‐fold in *Δmm*) and malate (10‐fold in *Δmm*) (Fig. 4C and D). Taken together, our glucose and glutamine tracer studies indicate that fumarate accumulation in *Δfh* arises from both glutamine‐derived carbon and ASL generated fumarate, whereas malate accumulation in *Δmm* primarily results from fumarate anaplerosis, with a lesser contribution from glutamine‐derived carbon. This apart, elevated malate in *Δmm*, could drive the reverse FH reaction, further contributing to fumarate accumulation.

**Figure 4.**
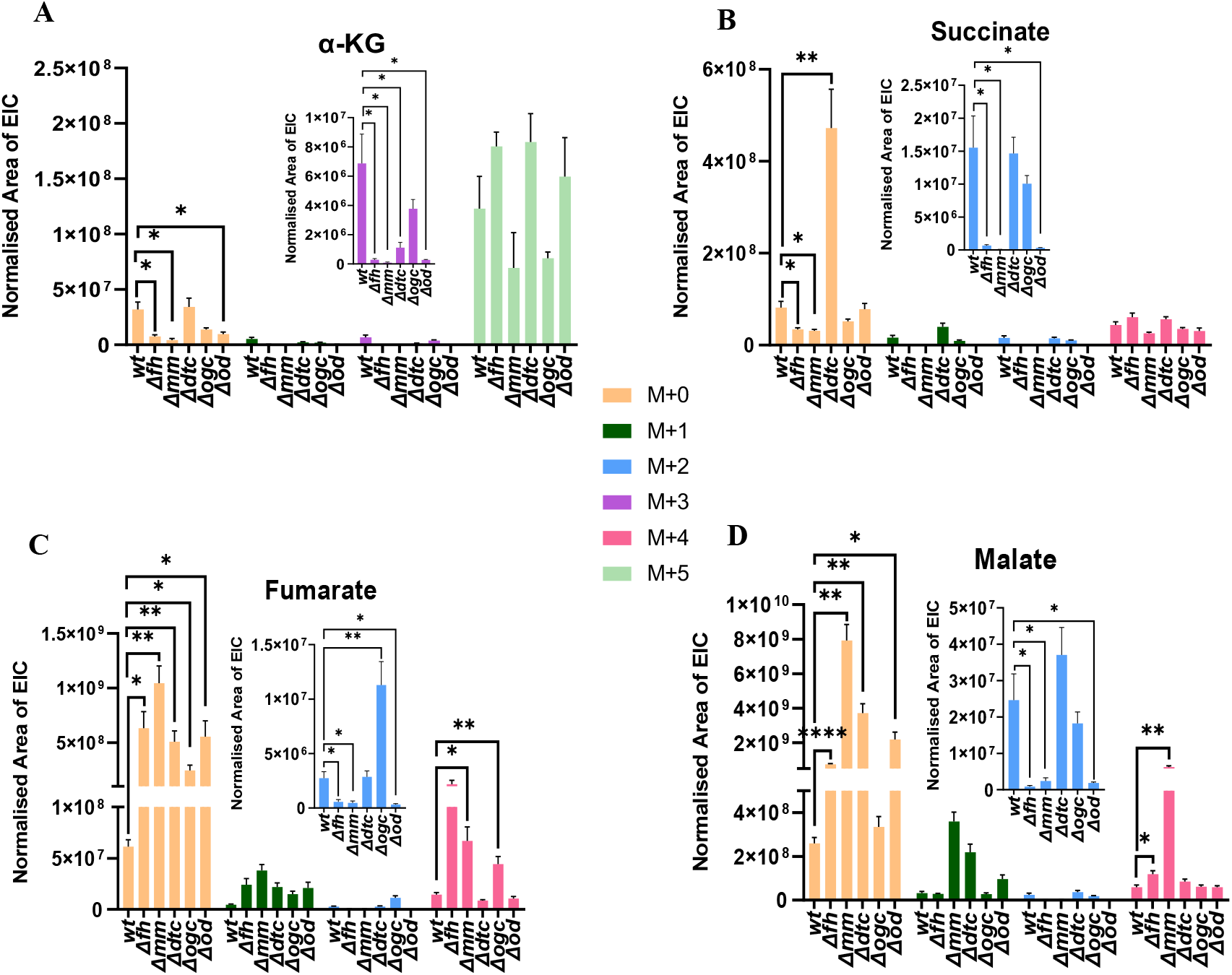
Incorporation of ^13^C_5_ ^15^N_2_‐glutamine‐derived carbon or nitrogen into the TCA cycle intermediates. The EIC areas of various isotopologues of (A) α‐KG, (B) succinate, (C) fumarate, and (D) malate in the WT and knockout gametocytes. Mean values with error bars representing SEM were plotted. The inset shows the zoomed‐in plots for some of the isotopologues. N=2 for WT and knockouts with two to three technical replicates each. Unpaired two‐tailed t‐tests were carried out using GraphPad Prism v9 (p<0.05 (*), p<0.005 (**), p<0.001 (***), p<0.0001 (****)).

With regard to *Δdtc* and *Δogc*, a block in the transport of cytosolic α‐KG (27) into the mitochondrion should either lead to an increase in the levels of M+5 α‐KG or a decrease in M+4 succinate, fumarate, and malate. However, the EIC area of these isotopologues remained unchanged in both the single knockouts and in *Δod* (Fig. 4A‐D), except for an increase in the M+4 fumarate in *Δogc* (Fig. 4C). Taken together, these results do not provide strong evidence that DTC and OGC are transporters of α‐KG. An increase in the EIC area of M+0 malate and fumarate in *Δdtc* and *Δod*, and M+0 fumarate in *Δogc* (Fig. 4C and D) indicates that MDH/ASL reactions contribute to this change. This suggests that DTC is a transporter of both dicarboxylic acids, whereas OGC transports fumarate, consistent with the glucose tracer results. Further, support for OGC functioning as a transporter of fumarate comes from the observed accumulation of M+2 (4.1‐fold) and M+4 (3.1‐fold) fumarate in *Δogc* (Fig. 4C). These isotopologues likely arise from the efflux of labelled OAA from the mitochondrion, followed by its conversion to fumarate through the sequential action of AAT, adenylosuccinate synthetase (ADSS), and ASL.

As expected, IMP remained largely unlabelled in all the five knockouts, as glutamine‐derived carbons are not incorporated into this metabolite (Fig. S23A). During XMP to GMP conversion, the nitrogen derived from glutamine is incorporated into GMP, producing a significant population M+0, N+1 GMP in both WT and knockouts (Fig. S23B). This confirms that IMP dehydrogenase (IMPDH) and GMP synthetase (GMPS) are active, forming the key pathway for GMP synthesis in the parasites. Due to active synthesis of N+1 Asp (Fig. S23D), we also observe about 69.9±3.2% labelling (combined N+1 of M+0, M+2, and M+4) of sAMP in WT, with knockouts showing a moderate reduction (Fig. S23C, Dataset 1).

All the atoms of aspartate feed into pyrimidine synthesis with the additional nitrogen coming from the side‐chain amide of glutamine. Although aspartate levels in the WT parasite were 57.4±9.4% M+0, N+1 labelled and remained largely similar in the knockouts (Fig. S23D), labelling in UMP was minimal in both WT and knockouts (Fig. S23E). Both tracer studies indicate active synthesis of sAMP and GMP in the knockouts and WT gametocytes, while pyrimidine nucleotide synthesis, as reflected by labelled UMP levels, is very low.

In ^13^C_5_^15^N_2_‐glutamine fed uRBCs, succinate, fumarate, and malate remained largely unlabelled (M+0 in the range of 90 – 92%) (Fig. S24B‐D). This clearly indicated that the labelling observed for these metabolites in *P. berghei* gametocytes arises from metabolism in the parasite alone. The metabolites, α‐KG and aspartate displayed labelling in uRBC, however the extent of labelling of M+5 α‐KG and M+0/N+1, M+4/N+0 aspartate in uRBC differed from that in *Pbwt* gametocytes (Fig. S24A and E).

## Discussion

Figure 5 shows a schematic overview of the metabolic consequences arising from the gene knockouts studied here. Our study has examined the essentiality of two mitochondrially localised TCA cycle genes (*fh* and *mqo*), two mitochondrial transporters (*dtc* and *ogc*), and the cytoplasmic *mdh* across erythrocytic and ookinete stages in *Pb*. All the seven knockouts‐*Δfh, Δmqo, Δmdh, Δmm, Δdtc, Δogc*, and *Δod*‐showed no growth impairment during the asexual blood stages of *Pb*. The viability of erythrocytic asexual stages of *Δfh* and *Δmqo* in both *Pb* (15) and *Pf* (16) has been previously reported, with the *Pb* mutants showing a delayed growth phenotype. Similarly, deletion of aconitase (*aco*) gene in *Pb* (13) also shows lower growth rate, whereas knockout in *Pf* (14) shows no change. Earlier studies on knockouts of other TCA cycle genes (14, 31–33) and *mdh* (22) also reported no impairment of asexual blood stage growth. The development of gametocytes was also not substantially impaired in any of the seven knockouts, with observations in *Δfh, Δmqo*, and *Δmdh* consistent with earlier findings in *Pb* (15, 22). Despite the increased flux through the TCA cycle and OXPHOS during the gametocyte stage (9, 13, 34), many genes associated with these two pathways were found to be non‐essential for the development of this stage (13, 14, 31, 35). Our results further reinforce the view on the non‐essentiality of the TCA cycle, anaplerosis and malate/OAA shuttle for gametocyte development. An exception is *Δaco* in *Pf*, which shows defects in gametocyte formation (14), unlike the phenotype observed in *Pb* (13).

**Figure 5.**
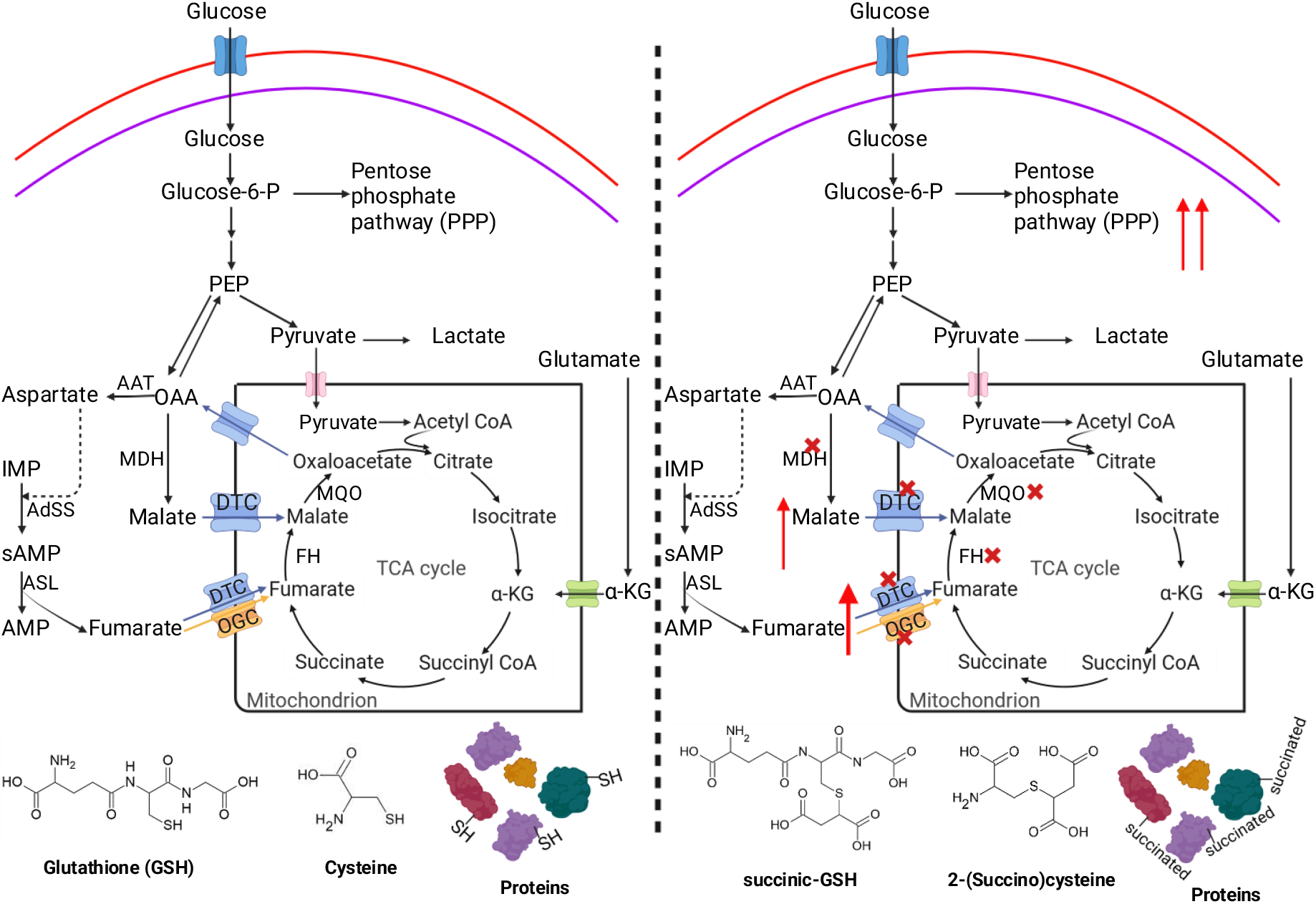
Metabolic consequences arising from abrogation of malate and fumarate anapleoris. Left panel, metabolic state of WT parasites with thiols of cysteinyl residues in GSH and proteins, and free cysteine largely lacking the modification, succination. DTC is a transporter of malate and fumarate and OGC a transporter of fumarate as evident from our studies. Right panel, knockout of *fh, mqo/mdh* (*mm), dtc, ogc* and *dtc/ogc* (*od*) (indicated with a red cross) leads to fumarate accumulation that in turn leads to succination of thiols resulting in oxidative stress, which is mitigated by enhanced NADPH levels via upregulation of PPP. Upward red arrows indicate increase in flux through PPP or increase in levels of fumarate and malate.

In contrast, the inability to form ookinetes by *P. berghei Δfh, Δmqo, Δmm, Δdtc, Δogc*, and *Δod* resembles the impaired mosquito stage development observed upon knockout of genes coding for ACO, flavoprotein subunit of succinate dehydrogenase, NADH dehydrogenase, E1 subunit of branched‐chain ketoacid dehydrogenase, and β‐subunit of ATP synthase in *Pb* (13, 31, 35–37), as well as ACO, the E1 subunit of α‐ketoglutarate dehydrogenase, and isocitrate dehydrogenase (IDH), in *Pf* (14, 33). The only knockout, *Δmdh* that in our study retained the ability to form ookinetes in *Pb*, is consistent with a previous report (22), wherein the lack of a pronounced phenotype was attributed to the continued synthesis of malate/NADH by the TCA cycle or replenishment of NAD^+^ by other dehydrogenases. However, as seen from our results, perturbations in the TCA cycle, the malate‐OAA shuttle, and the ETC in *PbΔmm* lead to a complete block in ookinete formation. The significant reduction in gametocyte to ookinete conversion in *Δdtc* and *Δogc* indicates the absence of an alternate mitochondrial carrier protein that could compensate for these transporters. Srivastava and coworkers have reported that maturing ookinetes are metabolically active, catabolizing both glucose and glutamine via the TCA cycle, albeit with enhanced dependence on glutaminolysis (13). Although we observed defects in ookinete formation in *Pb* lacking *fh, mqo, mm, dtc, ogc*, and *od*, it is possible that the earlier processes of male or female gamete formation are compromised.

Despite FH and MQO being enzymes and DTC and OGC being transporters, metabolomic analysis of the knockout gametocytes revealed a common metabolic phenotype: cysteine and GSH succination. The accumulation of succinated GSH may lead to oxidative stress. An increase in succination of a few proteins was also observed in *PbΔfh* gametocytes. The Michael addition of fumarate to free cysteine and cysteinyl residues is a rather slow non‐enzymatic reaction (28). Nevertheless, in the cellular context, fumarate has been classified as an oncometabolite (38) due to its ability to succinate cysteinyl residues in proteins and GSH (28, 39–44). *Plasmodium* possesses γ‐glutamylcysteine synthetase (γ‐GCS) and glutathione synthetase (GS) for the biosynthesis of GSH, and both glutathione reductase (GR) and the thioredoxin system for reduction of GSSG to GSH (45, 46). Mammalian GR shows activity only on GSSG and not on succ‐GSH (44), indicating that succ‐GSH cannot be converted back to GSH. In *P. berghei* lacking *fh*, the large increase in succ‐GSH accompanied by reduction in levels of GSH and GSSG, would likely result in enhanced oxidative stress as observed in mammalian cells (40, 44). As the knockout parasites examined in this study show increase in succ‐GSH at the gametocyte stage, the oxidative stress may persist during subsequent extracellular developmental stages in the mosquito, leading to, as observed, the almost complete absence of ookinete formation. Consistent with this interpretation, ^13^C_5_^15^N_2_‐glutamine tracer experiments showed similar percent labelling in GSSG across WT and knockout (Fig. S23F) gametocytes, indicating the absence of a compensatory upregulation in the production of this reducing agent.

GSH homeostasis is maintained through the reduction of GSSG by GR and synthesis of GSH via γ‐GCS and GS. Previous studies have shown that single knockouts of *gr* (47) or *γ‐gcs* (48) in *Pb* do not impair the erythrocytic stages but disrupt development of mosquito stages (47–49), a phenotype that is similar to the six knockouts analyzed here. Although knockout of *γ‐gcs* in *Pb* results in reduced levels of parasite‐generated GSH, development of the erythrocytic stages was unaffected, suggesting compensation by host‐derived GSH. However, as the mosquito stages are extracellular, their development is impaired probably due to reduced GSH levels and the resulting oxidative stress. A double knockout of *gr* and *γ‐gcs* could not be obtained, indicating that simultaneous elimination of both GSH synthesis and regeneration pathways is lethal to the parasite (47).

NADPH, an essential cofactor in the GSH and thioredoxin redox systems (46), is generated in the cytosol through the PPP and GDHa, with the PPP being the major contributor to the cellular levels of this cofactor (50–52). In our metabolomic analysis of the knockout lines, enhanced labelling of M+5 IMP following ^13^C_6_‐glucose tracing indicates an upregulation of ribose‐5‐phosphate production through the PPP pathway. This, in turn, would lead to increased cellular NADPH levels required for the activity of GR and thioredoxin, thereby enabling the reduction of GSSG to GSH. Such a mechanism could help maintain GSH homeostasis in gametocytes. Similar metabolic rewiring via increase in PPP flux, has been observed in FH‐deficient cancer cells (53, 54). Moreover, it should be noted that apart from oxidative stress arising from the reduction in GSH levels in the cytosol, the disruption of TCA cycle as a consequence of *fh, mqo*, and *od* knockouts would lead to lowering of IDH‐generated NADPH levels in the mitochondrion too.

Metabolomic analysis on *PfΔfhΔmqo* schizonts showed no change in the levels of PEP, decreased lactate levels, and increased lactoyl‐GSH, suggesting an increase in glycolytic flux (16). In contrast, we observed decreased lactoyl‐GSH levels in both *PbΔfh* and *PbΔmm* and lowered PEP levels in *PbΔfh* gametocytes. Glucose tracer experiments showed increased lactate production in *PbΔmm* gametocytes. The study on *PfΔfhΔmqo* was restricted to asexual stages with information on the development and viability of gametocyte or mosquito stages absent (16). Moreover, the metabolomic analyses were performed on intraerythrocytic schizonts that have reduced TCA cycle flux (9). *PfΔfhΔmqo* schizonts did not show elevated levels of fumarate or malate, with no report on the presence of succ‐GSH or 2SC in the published study (16). However, these parasites were less tolerant to high concentrations of exogenously added fumarate. The authors attributed the detoxification of the accumulated fumarate to host argininosuccinate lyase (ARSL) activity and reduced parasite ADSS activity leading to the observed increase in IMP and reduction in sAMP levels. ARSL catalyses the interconversion of argininosuccinate to arginine and fumarate, whereas ADSS catalyses the formation of sAMP through the condensation of aspartate with IMP. Despite the presence of ARSL in mouse erythrocytes (55, 56), we observed elevated levels of fumarate in *PbΔfh* and *PbΔmm* gametocytes. In contrast to the study on *Pf*, a decrease in the levels of IMP was observed. Our findings suggest the presence of fumarate toxicity in all the five knockouts examined. TCA cycle flux being different across schizonts and gametocytes (9, 13) could be the main contributing factor to the observed differences. This apart, species‐specific differences between *Pf* and *Pb* or differences in their host erythrocyte environment, may also influence fumarate detoxification. However, studies on *Pf* gametocytes lacking *fh* and *mqo* are needed to clarify the differences between the two species.

The proteoliposomal studies on PfDTC showed that it counter‐transports α‐KG with malate and OAA, and to a lesser extent with fumarate and succinate (25). However, it must be noted that these measurements were conducted using only a single labelled substrate, α‐KG. From the metabolomics study carried out by Rajaram and coworkers on mitochondrial transporter knockouts *mpc1*/*mpc2* (*mpc*, mitochondrial pyruvate carrier), *mpc1*/*mpc2*/*dtc* and *mpc1/mpc2/dtc/ogc*(*yhm2*) in *Pf* schizonts, the authors conclude that DTC and OGC facilitate transport of α‐KG (27). However, as this study lacks information on metabolomic analysis of individual *dtc* and *ogc* knockouts, a direct comparison with our results becomes difficult. Contrary to this, our metabolomic data from *PbΔdtc, PbΔogc*, and *PbΔod* gametocytes show distinct metabolic differences. The entry of α‐KG into the mitochondrion is unaffected in *PbΔdtc, PbΔogc*, and *PbΔod*, suggesting that other transporter/s expressed in the gametocyte stages may mediate α‐KG import. The significantly elevated level of malate, along with minor increases in the levels of fumarate, 2SC, and succ‐GSH in the *PbΔdtc* gametocytes, suggest that malate is the primary substrate of PbDTC, along with some affinity for fumarate. Conversely, the increase in levels of fumarate, 2SC, and succ‐GSH in *PbΔogc*, despite the presence of FH, indicate that OGC transports fumarate into the mitochondrion. The isotope tracer experiments further support the accumulation of cytosolic malate and fumarate in the absence of DTC and OGC, respectively. Glucose tracer studies also suggest that DTC may transport OAA. Surprisingly, the sequences of *Pf* and *Pb* DTC with 85% identity (Fig. S25A) render alterations in substrate specificity across species unlikely. Regarding OGC, although *Pf* and *Pb* OGC sequences have 74% identity, the *Pb* sequence has a N‐terminal extension of 12 residues while the *Pf* sequence has an insertion of 10 amino acids between residues 195‐204 (Fig. S25B). The *in silico* modelled structure of PfOGC identified residues K67, R151, Q152, and R262 as potentially involved in substrate transport (57). These residues are conserved across *Pb* and *Pf* OGC suggesting conservation of substrate specificity across the two species (Fig. S25B). The differences observed between our results and those reported by Rajaram and coworkers (27) could be due to differences in the TCA cycle flux between asexual intra‐erythrocytic stages and gametocytes (9, 13). Another observation from our studies is the high level of conversion of glutamine to α‐KG in WT gametocytes, with only a small fraction entering the TCA cycle. This is in contrast to the observations on *Pf* asexual stages, where majority of the glutamine‐derived α‐KG feeds into the TCA cycle (27).

## Conclusion

Due to the toxicity across multiple cell types arising from FH deficiency (41, 43, 58, 59), fumarate has been classified as an oncometabolite (38). Its extreme toxicity has also been observed in fumarase‐deficient *Mycobacterium tuberculosis* (60). From our studies on *Δfh Pb* gametocytes, the finding of succinated cysteinyl residues in GSH and in proteins, and in free cysteine, indicates the presence of fumarate toxicity in the knockout. We propose that fumarate reactivity, leading to oxidative stress, contributes significantly to the abolishment of ookinete formation. This inability to form ookinetes presents FH as a potential candidate for malaria transmission blocking. A chemical strategy that specifically leads to fumarate accumulation in the parasite without affecting the host can be achieved through selective inhibition of parasite FH. All *Plasmodium spp*. possess a Class I, oxygen sensitive 4Fe‐4S cluster‐containing FH (11), whereas human FH belongs to the Class II family (61). Previous studies have shown that D/L‐thiomalate inhibits PfFH with *K*_i_ values of 321 ± 26 nM and 548 ± 46 nM with malate and fumarate as substrates, respectively, while no inhibition of Class II FH was seen even at 10 mM inhibitor concentration (11). Interestingly, aurothiomalate (gold complexed with thiomalate) has been used for the treatment of rheumatoid arthritis (62–65). If thiomalate induces a metabolic phenotype in circulating gametocytes similar to that of the *fh* knockout, this compound or its analogs could serve as effective transmission‐blocking agents. In addition, as *Δmm, Δdtc, Δogc*, and *Δod Pb* gametocytes also show fumarate accumulation, MQO, DTC and OGC also become potential candidates for transmission blocking.

## Methods

Detailed methodology is provided in the *SI Appendix*.

### Generation of *P. berghei* knockouts

Generation of *fh* knockout has been described earlier (11). The knockout constructs for *mqo, mdh*, and *ogc* genes and library construct of *dtc* were procured from the PlasmoGEM repository, Wellcome Trust Sanger Institute, UK (66, 67). *Pbdtc* knockout construct was generated from the library construct using earlier methods (68, 69). Details of procedure followed for gene knockout, genotyping, and phenotypic analyses are provided in the *SI Appendix*. The primers used in the study are listed in Table S2.

### Metabolite extraction, relative quantification, and isotope labelling

Isolation of gametocytes free of asexual stages and host erythrocytes, isotope tracing, and metabolite extraction are provided in the *SI Appendix*.

### LC‐MS methods for semi‐targeted metabolomics and isotope tracing

The LC‐MS conditions, and procedures for metabolite identification, relative metabolite quantification using El‐MAVEN (70), natural abundance correction, and correction for in‐source malate dehydration are provided in the *SI Appendix*.

LC‐MS processed data is provided in Dataset S1.

## Acknowledgements

The pBCEN5‐GFP plasmid was a kind gift from Prof. Shiroh Iwanaga, Mie University, Japan. We acknowledge Prof. Ravi Manjithaya, Jawaharlal Nehru Centre for Advanced Scientific Research (JNCASR) for the use of the microscopy facility. A.C. and A.S. thank Department of Biotechnology, India, and University Grants Commission, India, respectively, for the JRF and SRF fellowship. The authors thank Aparna Dongre and Vinay Bulusu for discussion on metabolomics. The mass spectrometry facility of JNCASR is acknowledged. The work was supported by grants from the Department of Biotechnology, India (Project no. BT/INF/22/SP27679/2018). H.B. thanks JC Bose fellowship (Grant number: JCB/2019/000006), Science and Engineering Research Board (SERB) (Grant Number. CRG/2019/004150), Department of Science and Technology, Government of India, and intramural funding from JNCASR.

## Conflict of Interest

The authors declare no competing financial interests.

## Authorship contribution statement

**Anusha Chandrashekarmath:** Data curation, Formal analysis, Investigation, Methodology, Validation, Visualization, Writing – original draft, Writing – review and editing. **Arpitha Suryavanshi:** Data curation, Formal analysis, Investigation, Methodology, Validation, Visualization. **Chitralekha Sen Roy:** Investigation. **Hemalatha Balaram:** Conceptualization, Data curation, Formal analysis, Funding acquisition, Methodology, Project administration, Resources, Supervision, Validation, Visualization, Writing – original draft, Writing – review and editing.

## Notes

### Competing Interest Statement

The authors have declared no competing interest.

