## Supplementary material for "Perturbations in fumarate levels in *Plasmodium berghei* leads to cysteine succination and impairs ookinete formation": SI Appendix

**This PDF file includes:**

Supporting text
Figures S1 to S25
Tables S1 to S2
SI References

**Other supporting materials for this manuscript include the following:**

Dataset 1

**Supporting text**

**Methods**

**Generation of plasmid constructs for gene knockout and tagging in *P. berghei***
The pJAZZ-OK-*Pbdtc*, carrying full length *P. berghei dtc* (PBANKA_0706700) and flanking regions generated as a genomic DNA library construct was procured from PlasmoGEM repository, Wellcome Trust Sanger Institute, UK (1, 2). *Pbdtc* knockout construct was generated from the library construct using earlier methods (3, 4). Briefly, this involved replacement of the gene of interest with the bacterial positive/negative marker, Zeo-PheS followed by an exchange for *Plasmodium* positive/negative selection marker, hDHFR-yFCU. Knockout constructs for *mqo* (PBANKA_1116300), *mdh* (PBANKA_1117700), and *ogc* (PBANKA_1438700) genes were procured from the PlasmoGEM repository. All constructs procured from the repository or generated in the study were validated by restriction digestion or PCR.

For the generation of PbCEN-PbOGC-GFP construct, the *Pbogc* gene was amplified by PCR using *P. berghei* genomic DNA as template, and primers P33 and P34 (Table S2). The BamHI linearised plasmid, pBCEN5-GFP and the *Pbogc* insert were setup for In-Fusion cloning (Takara Bio). *E. coli* XL1-Blue competent cells were transformed with the DNA mix, and the obtained colonies were screened by PCR for the presence of the *ogc* gene and the plasmids isolated from the positive clones were confirmed by DNA sequencing.

**Ethics statement**
The handling of BALB/c or C57BL/6 mice were carried out by adhering to the standard procedures prescribed by the Committee for the Purpose of Control and Supervision of Experiments on Animals (CPCSEA), a statutory body under the Prevention of Cruelty to Animals Act, 1960 and Breeding and Experimentation Rules 1998, Constitution of India. The mice were maintained at the Animal Facility, JNCASR and the study was approved by Institutional animal ethics committee (IAEC) of JNCASR that comes under the purview of CPCSEA.

**Culturing of erythrocytic stages of *P. berghei* and cryopreservation**
*Plasmodium berghei* ANKA (MRA-671) strain used for the study was procured from Malaria Research and Reference Reagent Resource Center (MR4, USA) and was maintained in BALB/c or C57BL/6 mice (6-8 weeks old). Around 100 µl of parasite cell suspension from a cryopreserved stock was injected intraperitoneally into a mouse. Monitoring of parasitemia in mice was performed by drawing infected blood from a tail snip, followed by preparation of smear on a glass slide, that was fixed with methanol and stained with Giemsa stain. The smear was examined under the microscope using a 100 x oil immersion objective lens. For cryopreservation, blood with a parasitemia of approximately 5 %, was harvested from the infected mouse into an incomplete medium (RPMI 1640 supplemented with L-glutamine, 25 mM HEPES, pH 7.4) containing 200 U $ml^{-1}$ of heparin. Thereafter, 0.2 ml of this suspension was mixed with 30 % glycerol in 1x PBS (0.3 ml) in a cryotube and flash-frozen in liquid nitrogen. Revival of cryopreserved stock was done by thawing at 37 °C for 1-2 min and 100 µl of the cell suspension was injected into a mouse intraperitoneally.

**Transfection of asexual erythrocytic stages of *P. berghei***
For generation of single knockouts and PbCEN-PbOGC-GFP transgenic line, 100 µl of cryopreserved stock of wild-type *P. berghei ANKA* parasites was injected intra-peritoneally into a mouse. For the generation of *P. berghei ΔmqoΔmdh* and *ΔdtcΔogc*, clonal line of marker-free *Δmqo* and *Δdtc* parasites was injected, respectively. The parasitemia was monitored by

microscopic examination of Giemsa-stained smears of blood drawn from the tail vein of the infected mouse. The protocol followed for transfection was as described earlier (5) with minor modifications. For the generation of *Pbmqo* knockout, the schizonts were obtained by culturing infected blood *in vitro*. Briefly, the infected blood was centrifuged at 450 x g for 10 min followed by suspension of the cells in 50 ml of complete medium (RPMI-1640 with glutamine, 25 mM HEPES, 10 mM $NaHCO_3$, and 20% fetal bovine serum) and transfer to a 250 ml reagent bottle. The bottle was gassed for 3 min with a gas mixture of 5% $O_2$, 5% $CO_2$, and 90% $N_2$, and tightly sealed. The bottle was incubated at 36.5 °C, overnight at 90 rpm, that was just enough to prevent the cells from settling at the bottom of the bottle. For generation of the remaining transgenic lines, first the presence of schizonts was confirmed through microscopic examination of Giemsa-stained smears followed by drawing the blood into 0.4 ml of 1 mg $ml^{-1}$ heparin in incomplete RPMI-1640 medium. The parasite containing blood was layered on 50-60% Nycodenz solution in 1x PBS and centrifuged at 200 x g for 7-10 minutes with no brake applied. A layer containing schizonts that separated at the interface was collected, washed thrice with incomplete RPMI 1640 medium and a parasite pellet was obtained. P5 transfection kit from Lonza and NotI-digested linearised knockout constructs or PbCEN5_PbOGC_GFP plasmid were used for transfection. The parasite pellet was resuspended in 110 µl of nucleofector solution (82 µl of P5 + 18 µl of supplement + 10 µl of plasmid) and was pulsed using the pulse code FP 167 or FP148 of 4D Amaxa Nucleofector. To the transfected mix, 50 µl of incomplete medium was added to aid in the recovery of the cells and administered intravenously to a healthy mouse. The presence of infected erythrocytes was confirmed after 3 days of injection, after which, pyrimethamine (7 mg $ml^{-1}$) drug pressure (against the *Plasmodium* selectable marker hDHFR) was applied by providing it *ad libitum*. Smears of the blood from the infected mouse were made frequently, Giemsa stained and microscopically examined for the presence of drug-resistant parasites. On obtaining pyrimethamine resistant parasites, the infected blood at 10 % parasitemia was harvested from the mouse, stocks were cryopreserved and genotyping was done.

**Genotyping of drug-resistant *P. berghei***

On obtaining pyrimethamine-resistant parasites, genomic DNA was isolated and genotyping was performed by PCR. For the genomic DNA isolation, the harvested blood containing *P. berghei* parasites was mixed with 1 x erythrocyte lysis buffer (150 mM $NH_4Cl$, 10 mM $KHCO_3$, 1 mM $Na_2EDTA$, pH 7.4), incubated at 4 °C for 10 minutes, and centrifuged at 500 x g for 10 minutes to obtain the parasite pellet. The pellet was resuspended in 350 µl of TNE buffer (10 mM Tris pH 8.0, 100 mM NaCl, 5 mM EDTA pH 8.0). To this parasite suspension, RNAseA (0.2 mg $ml^{-1}$) and SDS (1%) were added, and TNE was added to make up the volume to 500 µl followed by incubation at 37 °C for 10 minutes. Pronase (0.2 mg $ml^{-1}$) was added and incubated for an additional 1 hour. 1 mL of Tris-saturated phenol was added, gently mixed by inverting, and centrifuged at 10000 x g for 5 minutes. The aqueous layer was transferred to a fresh tube, 1 ml of 1:1 phenol: chloroform was added and centrifuged at 10000 x g for 5 minutes. The top aqueous layer was transferred to a fresh tube, 1 ml of chloroform-isoamyl alcohol was added, and centrifuged at 10000 x g for 5 minutes. The supernatant was transferred to a fresh tube to which $1/10^{th}$ the volume of 3 M sodium acetate, pH 4.5 and 1 ml of absolute ethanol were added sequentially. Upon gentle inversion and mixing of the contents in the tube and centrifuging at 10000 x g for 1 min, a DNA pellet was obtained. The supernatant was discarded, and 70% ethanol was added to the pellet, followed by centrifugation at 10000 x g for 1 min. The DNA pellet was left to air dry and resuspended in water, and stored at -20 °C. The presence of the selectable marker, the absence of a gene, and the locus of integration for each gene knockout were validated by PCR genotyping. Primers used for genotyping are listed in Table S2.

**Localisation of OGC in *P. berghei***

The glycerol stock of *P. berghei*, WT and pBCEN5-PbOGC-GFP transfectants were injected into BALB/c mice. At a parasitemia of 10-20%, blood was collected in heparin containing incomplete RPMI 1640 medium. To the parasite pellet obtained after centrifugation, one wash with warm incomplete RPMI 1640 medium was performed. The parasites were resuspended in incomplete RPMI 1640 containing 120 µg $ml^{-1}$ of MitoTracker Orange (Thermo Scientific) and incubated in candle jar at 37 °C for 25 minutes with slight rotation. This was followed by Hoescht staining at 10 µg $ml^{-1}$ for 10 minutes at 37 °C in a candle jar with slight rotation. Later, the parasite pellet was washed thrice with 1 x PBS. The pellet was resuspended in Vectashield (Vector laboratories) and mounted on a glass slide. Images were acquired using DeltaVision microscope connected to CoolSNAP_HQ2 camera using 100 x oil immersion objective. The microscopy images were processed using Fiji software (6).

**Recycling of *Plasmodium* drug selectable marker by negative selection**

To obtain marker-free parasites, the genetically modified and pyrimethamine-resistant parasites were subjected to negative selection for excision of the drug-selectable marker. A mouse (BALB/c) was injected intra-peritoneally with 0.1 ml of a thawed suspension of the cryopreserved stock of a mutant *P. berghei* line. The parasitemia of the mouse was monitored frequently until it reached between 0.1 % and 0.5 %. Thereafter, the infected mouse was fed *ad libitum* with 5-fluorocytosine (5-FC) at 1mg $ml^{-1}$, provided in drinking water. Microscopic examination of Giemsa-stained blood smear prepared 48 hours after drug treatment indicated visual absence of parasites. The drug-containing water was replenished every 4 days of treatment till the reappearance of parasites was seen in the blood smear. Blood harbouring 5-FC drug-resistant parasites was harvested from the mouse for cryopreservation and genomic DNA isolation. Genotyping by PCR was carried out using appropriate primers to check for the absence of the hDHFR-yFCU marker. The 5FC-resistant parasites were used for cloning by limiting dilution to obtain marker-free clones. The clones were validated by PCR genotyping for the absence of marker and sensitivity of the clonal line to pyrimethamine before using for the next genetic manipulation experiment.

**Generation of clonal lines of *P. berghei* by limiting dilution**

A mouse was injected intraperitoneally with the cryopreserved stock of the mutant *P. berghei* and parasitemia was monitored regularly. When parasitemia reached between 0.5 and 1 %, infected blood was harvested from the mouse. The blood was diluted 100-fold in an incomplete medium and the number of erythrocytes per µl of the diluted blood was estimated using a Neubauer chamber. The number of infected erythrocytes per µl of the diluted blood was calculated by multiplying the total number of erythrocytes per µl with the percent parasitemia. The infected blood was diluted in incomplete medium to obtain one infected erythrocyte per 100 µl suspension that was injected intravenously into each mouse of a batch comprising 10 mice. The appearance of parasites in the mice was checked by making smears regularly. Blood was drawn from mice that showed infection and used for cryopreservation and genomic DNA isolation.

**Determination of the growth rate of intra-erythrocytic stages of *P. berghei***

Cryopreserved stocks of WT and clones of null mutants of *P. berghei* were injected into independent mice and parasitemia was monitored by microscopic examination of Giemsa-stained smears. At a parasitemia of < 2%, around 10 µl of the infected blood was diluted 100-fold in incomplete medium and mounted on a Neubauer chamber for estimation of number of erythrocytes per µl of blood. The study was carried out in four groups each with its wild-type control. Groups 1, 2 and 3 involving *fh*, *mqo*, *mdh*, *dtc*, and *mqo/mdh* consisted of 5 mice for each parasite line. Group 4 involving *ogc* and *dtc/ogc* had 3-5 mice for each parasite line. For groups 1-3, $1 \times 10^6$ and for group 4, $1 \times 10^4$ parasites were used for establishing infection. The parasitemia in mice infected with WT and null mutants was monitored daily by microscopic examination of Giemsa-stained blood smears. The growth rate of the parasites was evaluated by plotting the daily monitored parasitemia against time (number of days). The mortality rate of the mice infected with either WT or null mutants was estimated by plotting the percentage survival of mice versus time (number of days).

**Estimation of percent male and female gametocytes of *P. berghei***

The ability of the null mutants to form gametocytes was assessed along with enumeration of gametocytaemia. The mice were first administered phenylhydrazine (0.05 mg $g^{-1}$ body weight of mouse) to induce reticulocytosis two days before infection with parasites. The study was carried out in three groups each with its wild-type control. Groups 1 and 2 involving *fh*, *mqo*, *mdh*, *dtc*, and *mqo/mdh* consisted of 4 mice for each parasite line. Group 3 involving *ogc* and *dtc/ogc* had 5 mice for each parasite line. For groups 1 and 2, $1 \times 10^7$ parasites and for group 3, $5 \times 10^5$ parasites were intraperitoneally injected into each mouse. After 2-4 days, the establishment of infection in all mice was checked by microscopic assessment of Giemsa-stained smears. The mice were treated with sulfadiazine (30 mg $L^{-1}$) *ad libitum* to kill the asexual stages of the parasites. Giemsa-stained smears made 48 hours after sulfadiazine treatment were microscopically examined for the enumeration of male and female gametocytes.

***In vitro* ookinete maturation of *P. berghei* gametocytes**

For the conversion of gametocytes to ookinetes, the steps involved harvest of gametocyte-enriched blood, *in vitro* activation of gametocytes to form gametes, and fertilization to form zygotes followed by maturation to ookinetes. The protocol followed for gametocyte enrichment of mouse blood was as mentioned in the above section. Around 1 ml of gametocyte containing blood collected in heparin from the infected mouse, was diluted to a total volume of 30 ml in the ookinete culture medium (RPMI 1640, 25 mM HEPES, 20 % foetal bovine serum, 24 mM $NaHCO_3$, 367.3 mM hypoxanthine, 100 μM xanthurenic acid, pH 8) and incubated at 19 ºC for 24 hours. The ookinete production (ookinete conversion) rates, defined as the percentage of female gametes that develop into mature ookinetes were determined by examination of Giemsa-stained smears. The % ookinete conversion was estimated using the formula reported (7).

% ookinete conversion = (Number of mature ookinetes * 100) / (Number of female gametes + Number of zygotes + Number of ookinetes).

**Enrichment, isolation, and equilibration of *P. berghei* gametocytes**

Mice were injected intraperitoneally with phenylhydrazine (100 μl of 12.5 mg $ml^{-1}$ prepared in 0.9% NaCl) and after two days, were infected with WT *P. berghei* ANKA or the knockout lines. The parasitemia was monitored by drawing blood from a tail snip and observing the Giemsa-stained smears under the microscope (100 x oil immersion objective). Once the parasitemia reached 10 - 20%, sulfadiazine (30 mg $L^{-1}$, dissolved in drinking water) drug pressure was applied for two days. On the third day, Giemsa-stained smears were checked under the microscope to ensure the death of all asexual stages and for the presence of gametocytes. Thereafter, the mice were bled and the blood collected in warm heparin in RPMI 1640 medium (RPMI 1640 10.4 g $L^{-1}$, HEPES 5.96 g $L^{-1}$, $NaHCO_3$ 2 g $L^{-1}$, hypoxanthine 100 μM, and thymidine 16 μM) was immediately transferred to a 37 °C incubator. All the subsequent steps were carried out at 37 °C. Leucocytes were removed by passing the blood through cellulose packed in Pierce centrifuge columns (Thermo Scientific). The pellet was resuspended with RPMI 1640 medium such that it was 0.46 times the initial volume of blood. This was layered on 57% Nycodenz made in 1 x PBS and centrifuged at 1000 x g for 20 minutes without brake. The gametocyte layer was carefully collected and washed with 5 ml of minimal medium (composition adapted from (8), provided in Dataset 1) twice, and the final pellet was resuspended in minimal medium to a final volume of 1 ml. 50-100 µl of this suspension was used to make smears and enumerate number of erythrocytes per µl of suspension using hemocytometer. The remaining 900-950 µl was added to 5 ml of minimal medium in a T25 flask and equilibrated at 37 °C for 90 minutes with mild rotation. Using the count estimated with the hemocytometer, aliquots of 2-3 x $10^7$ parasites were added to 5 ml minimal medium in a T25 flask and incubated with or without isotope tracers (mentioned in the sections below) for two hours at 37 °C.

**Metabolite extraction**

The parasite suspension was centrifuged at 1000 x g for 2 minutes, the supernatant removed, and the cell pellet resuspended in ice-cold 1x PBS. This suspension was centrifuged at 16000 x g for 1 minute at 4 °C, followed by another PBS wash. The pellet was solubilised using 1 ml of ice-cold 90% methanol prepared using mass spectrometry (MS) grade water and methanol. Metabolites were extracted by brief vortexing for 10 seconds. This step was repeated twice. During extraction, the sample was spiked with the internal standards, 3-(cyclohexylamino)-1-propanesulfonic acid (CAPS) (15µM), piperazine-1,4-bis (2-ethanesulfonic acid) (PIPES) (30µM) (9), and N-methyl-D-glucamine (NMG) (10µM) (10). The extract was centrifuged at 16000 x g for 15 minutes at 4 °C. 950 µl of the supernatant was transferred to a fresh vial and stored at -80 °C. The extract was later dried under nitrogen flow and stored at -80 °C until analysis.

In all the experiments, blood from two or three mice was pooled, and leucocytes were removed and layered on Nycodenz. The gametocytes obtained were split into two or three flasks equally to yield two or three technical replicates.

As a control, metabolites were extracted from uninfected erythrocytes (uRBC). Prior to harvest of blood, mice were treated with phenylhydrazine and sulfadiazine to mimic the test condition and the protocols starting from bleeding, removal of leucocytes, equilibration, and extraction were similar to those followed for parasite-infected blood. All cell preparations were ensured to be free of asexual blood stages with uninfected erythrocyte contamination below 5%.

**Stable isotope tracing with U-$^{13}C_6$-glucose and U-$^{13}C_5{}^{15}N_2$-glutamine**

The isotope tracers $^{13}C_6$-glucose and $^{13}C_5{}^{15}N_2$-glutamine were procured from Cambridge Isotopes Laboratories, USA. The procedure for gametocyte enrichment, harvesting, leucocyte removal, and equilibrating the gametocytes in minimal medium was the same as mentioned above up to the hemocytometer counting step. After 90 minutes of equilibration, the parasite suspension was centrifuged at 1000 x g for 2 minutes, the medium was removed, and the pellet was washed in minimal medium without glucose or glutamine. The pellet was resuspended in minimal medium without glucose or glutamine and split in such a way that each flask had 2-3 x $10^7$ parasites. Each T25 flask had 5 ml of minimal medium with $^{13}C_6$-glucose at a concentration of 8 mM or $^{13}C_5{}^{15}N_2$-glutamine at a concentration of 2 mM and was incubated at 37 °C for another 2 hours at 160 rpm. Metabolite extraction was the same as mentioned above.

**Liquid chromatography-mass spectrometry (LC-MS) of extracted metabolites**

The dried extract was solubilised in 30 µl of freshly prepared 40% MS grade acetonitrile/water, vortexed for 20 minutes at 4 °C and centrifuged at 16000 x g for 15 minutes at 4 °C. 20 µl of the supernatant was transferred to the autosampler vial, which was placed in the sampler compartment of the UHPLC. 4 µl of the extract was used for each run. The remaining sample was stored at -80 °C.

Mass spectral analysis was carried out on a Q Exactive HF mass spectrometer (Thermo Scientific, USA) equipped with a Dionex Ultimate 3000 UHPLC system. The solubilised metabolite mixture was injected into a ZIC-pHILIC column (Sequant, 150 x 2.1 mm, 5 µm) fitted with a guard column (20 x 2.1 mm). The column temperature was maintained at 45 °C, and the autosampler temperature was set at 10 °C. The solvent systems used were: Solvent A, 20 mM ammonium carbonate in water, pH 9.4, and Solvent B, 100% acetonitrile. Metabolites were eluted over a 45-minute run consisting of an initial equilibration with 80% B for 1 min, followed by gradients of 80% B to 40% B in 20 min and 40% B to 5% B in 10 min. Thereafter, isocratic flows at 5% B for 3 min, 5%-80% B in 1 min, and finally equilibration at 80% B for another 10 mins was followed. MS data were acquired in dual polarity mode using Xcalibur software v4.1 (Thermo Scientific) with Full MS resolution of 60,000 at m/z 200 Th, 3 microscans, AGC target of 1e6, and maximum IT of 200 ms. The scan range used was 70 to 1050 m/z. HESI source was used with sheath gas and auxiliary gas flow rates of 30 and 15, respectively, spray voltage of 4 kV (positive) and 3.5 kV (negative), capillary temperature of 320 °C, S-lens RF level at 50, and auxiliary gas heater temperature at 100 °C. Mass spectrometric data were acquired at the Central Instrumentation Facility, Molecular Biology and Genetics Unit, JNCASR.

A few samples of extracts from *P. berghei* gametocytes were initially examined in both MS and MS/MS mode and later, for the purpose of relative quantification, only in the MS mode. This generated MS/MS fragmentation data for a metabolite of a specific mass served as a basis for validation, as and when required. These MS/MS spectra could be compared with available MS/MS patterns in the database, aiding, particularly, in the identification of metabolites that were not included as standards. MS2 was acquired at 30,000 resolution, 2 microscans, AGC target of 1e5, maximum IT of 100ms, loop count of 10, isolation window of 4 m/z, normalised collision energy of 30, and dynamic exclusion of 10s. Metabolites such as oxidised glutathione (GSSG), 2-succino-cysteine (2SC), and succinic-GSH were identified by their masses and established by MS/MS patterns. Only the MS alone mode of the sample runs was considered for the final analysis.

**Analysis of mass spectrometric data**

Semi-targeted metabolomics was carried out for 23 metabolites of interest for which retention time (RT) on the ZIC-pHILIC column was confirmed either by injecting standards or where available, by comparing with MS/MS spectra in the database (Fig. S8). The extracted ion chromatogram (EIC) for each of these metabolites obtained using the El-MAVEN software (11) was individually examined and only those that yielded reliable and reproducible peaks were considered for further analyses.

The LC-MS data were analysed using El-MAVEN software (v0.12.1-beta) (11). The data obtained in .RAW format was converted to mzXML using RawConverter software (12). A list of compounds for which the retention time was known was input into El-MAVEN as a targeted compound list along with the mzXML data files. Processing was carried out individually with positive polarity and negative polarity using the following settings: maximum retention time difference between peaks in a group was set to 1 minute; EIC extraction window was set to 5 ppm, and match retention time was set to 1 minute. The area of the extracted peak was used for the quantification of a metabolite. The area generated from the software was exported in .csv format and further analysis across the samples was done in Excel. For samples labelled with

$^{13}C$-glucose and $^{13}C^{15}N$-glutamine, similar steps were carried out, along with choosing the right isotope tracer/s. After extracting the area for each of the isotopologues of a metabolite in stable isotope labelled samples, natural abundance correction was carried out using the graphical user interface R package of IsoCorrectoR (13). The tracer impurity correction was also applied. The values of the EIC areas were normalised to $3x10^7$ cells before comparing across WT and knockouts. GraphPad Prism version 9 was used to perform statistical analysis and to generate the graphs. Statistical analyses were done using unpaired two-tailed t-tests. If the F-test showed significant variances, Welch's correction was performed where necessary. Statistical significance is shown for those isotopologues that showed substantial differences in levels from WT.

Among the internal standards spiked during the extraction procedure, NMG was not included in the analysis due to the broad nature of its EIC. The detection and variability of CAPS and PIPES across 100-104 samples in negative and positive polarity is summarised in Dataset S1. As the EIC area of these metabolites did not vary significantly across the samples, normalisation with regard to these values was not performed. However, RT correction was applied in a few samples due to changes in RT of internal standard.

A total of 17 quality control (QC) runs were carried out during the course of the LC-MS study. These runs were carried out to establish the level of variation over the time course of the study. The QC sample consisted of 10 metabolites as shown in the Dataset S1. The areas of EIC across all the runs are listed in the Dataset S1 along with the average and standard deviation. A shift in RT for aspartate, α-KG, glutamate, and PIPES was observed in QC_16 and the EIC values were obtained for the corrected RT values. The *P. berghei* samples run during this period had also shown shifts in RT, which were corrected.

**Correction for in-source fragmentation of malate to fumarate**
With the gradient and flow rate used for the ZIC-pHILIC column, we could not resolve the elution of malate and fumarate. Malate undergoes in-source dehydration during the ionisation process leading to the formation of fumarate (14). To estimate the levels of fumarate produced from malate due to in-source water loss, the malate standard curve was generated using 7 different concentrations of malate standard. The concentration ranged from 0.5 µM to 50 µM, and 2 µl of each concentration was injected (1 pmol to 100 pmol) onto the ZIC-pHILIC column. Full MS/MS scan was carried out with the same gradient and MS settings as detailed above. The scan range of 55-825 m/z was used. Data were processed using the Processing setup of Xcalibur software v4.1 (Thermo Scientific). The area of the extracted ion chromatogram (EIC) for m/z values for malate and fumarate was subtracted from their respective blanks and plotted. For each injection, the EIC area corresponding to m/z of malate and the area for MS-generated fumarate were plotted, giving rise to malate standard curve and the derived standard curve for MS-generated fumarate (Dataset S1). From this study, it was estimated that the amount of in-source produced fumarate was 13% of the malate area (Dataset S1). Fumarate levels were corrected in the unlabelled data. Corrections that led to negative numbers were eliminated from analysis. It should be noted that despite this elimination of negative numbers, a minimum of at least 3 replicate data were available. The steps used for correction are as listed below.

1. Malate and fumarate areas were computed from El-MAVEN for each sample of metabolites extracted from *P. berghei* gametocytes.
2. The concentration of malate was computed from the malate standard curve.
3. This concentration of malate was used to back calculate the fumarate area. This was done using the derived standard curve for MS-generated fumarate.

4. This computed fumarate area was subtracted from the area of m/z corresponding to fumarate in the sample. This generated the corrected area of fumarate, and this was compared across WT and knockout samples.

**Detection of succinated proteins in *PbΔfh* gametocytes by Western blot**

The *P. berghei* WT and *Δfh* gametocytes were isolated, equilibrated, and incubated in minimal medium as described above. The parasites were washed with 1x PBS, twice and the cell pellet was resuspended in 1ml of 1x PBS containing protease inhibitor cocktail and 1 mM PMSF. Lysis was carried out with 10 rounds of freeze (liquid nitrogen)-thaw (37 °C) cycles and centrifuged at 16000 x g for 15 minutes to obtain a clear lysate. Protein concentration in the lysate was estimated with the Bradford reagent (Sigma) using BSA as the standard. The 4x SDS-dye to a final concentration of 1x was mixed with 50 µg equivalent of lysate of both WT and *Δfh* gametocytes, boiled and loaded on to 12% SDS-PAGE. The blotting of the gel was done on PVDF membrane using the semi-dry method. Following transfer, the membrane was blocked using 5% skimmed milk in 1x PBS. Thereafter, the blot was incubated with rabbit anti-2SC (1:1500) (Biosynth, UK) primary antibody for over 12 hours in the cold, washed with 1x PBS containing 0.1% Tween-20 and finally incubated with goat anti-Rabbit HRP (1:3000) as the secondary antibody. The blot was developed by using the chemiluminescence reagent from Advansta, USA.

**Proteomic analysis of *Pbwt* and *PbΔfh* gametocytes**

The procedures followed for the enrichment, isolation, equilibration, and incubation in minimal medium of *Pbwt* and *PbΔfh* gametocytes were the same as described above. After equilibration, the parasites were pelleted by centrifugation at 1000 x g for 3 minutes, followed by two washes with 1x PBS. The parasite pellet was stored at -80 °C until further processing. The gametocyte pellet was resuspended in 1 ml of 1x PBS along with protease inhibitor cocktail and PMSF and lysed by freeze (liquid $N_2$)-thaw (37 °C) cycles. The debris were pelleted down by centrifugation at 16000 x g for 15 minutes at 4 °C. The clear protein lysate was transferred to a fresh tube and protein estimation was carried out using the Bradford reagent (Sigma) with BSA as the standard. A volume equivalent to 100 µg of total protein from each of *Pbwt* and *PbΔfh* lysate was mixed with 4x SDS-dye, boiled and loaded on a 10% SDS-PAGE, and stained using Coomassie-Brilliant Blue dye (Sigma). Equivalent mass protein bands ranging from 140 KDa to 35 KDa were cut out to obtain 5 fractions each of *Pbwt* and *PbΔfh* and subjected to in-gel trypsin digestion. The bands lower than 35 KDa and higher than 140 KDa were not further processed as their intensity in the Coomassie-stained gel was very low. The entire proteomic analysis including gametocyte protein extraction, separation on SDS-PAGE, in-gel tryptic digestion, and MS/MS analysis was done in duplicate.

In-gel trypsin digestion was carried out using the protocol of Shevchenko and coworkers (15) with some modifications. Briefly, the gel was cut into small pieces of 1mm cube each and washed twice with mass spectrometry grade water and destained twice using 1:1 of mass spectrometry grade methanol:50 mM ammonium bicarbonate, pH 7.8, for 15 minutes. The gel pieces were dried for 5 minutes using 1:1 of mass spectrometry grade acetonitrile:50 mM ammonium bicarbonate, pH 7.8, followed by 30 seconds of 100% acetonitrile. The trace amount of remaining liquid was removed by drying in a Speed-Vac for 10 minutes. The gel pieces were hydrated using 25 mM DTT in 50 mM ammonium bicarbonate and incubated at 56 °C for 25 minutes. Thereafter, the gel pieces were washed with water, dried in acetonitrile as described above. The gel pieces were rehydrated using 30 mM iodoacetamide in 50 mM ammonium bicarbonate and incubated for 30 minutes at room temperature in the dark. The gel pieces were washed and dried as detailed above and rehydrated in 50 mM ammonium bicarbonate containing trypsin (Promega) and incubated for 12 hours at 37 °C. Incubation of the gel pieces with newly added trypsin was continued for an additional 4 hours. The supernatant containing the tryptic peptides was collected, and the gel pieces were further extracted with 50% acetonitrile containing 5% formic acid by vortexing for 20 minutes and sonication in a water bath sonicator for 15 minutes. The extraction process was carried out twice and all the extracted fractions were clubbed and dried using a Speed-Vac. The dried tryptic peptides were resuspended in 2% acetonitrile containing 0.1% formic acid and desalted using C18 spin columns (Thermo Scientific). Peptides were eluted from the C18 spin columns using 60% acetonitrile containing 0.1% formic acid and dried using a Speed-Vac.

The desalted tryptic peptides were resuspended in 25 µl of 2% acetonitrile containing 0.1% formic acid, vortexed for 20 minutes and centrifuged at 16000 x g for 15 minutes. 2 µl of the supernatant was injected into the column. Easy-Spray column (Thermo Fisher Scientific, PepMap RSLC C18 of 2 µm particle size, 100 Å pore size, 75 µm inner diameter, either 25 cm or 50 cm in length) along with a guard column (Acclaim PepMap100, C18, 3 µm particle size, 100 Å pore size, 75 µm inner diameter, 2 cm length) were used. Three corresponding fractions of WT and *fh* knockout were injected into a 25 cm, and two corresponding fractions were injected into a 50 cm Easy-Spray column. The nano LC system was connected to a Q-exactive HF mass spectrometer (Thermo Scientific). The column was equilibrated with solvent A for 10 column volumes before sample injection. The peptides were eluted using 2% acetonitrile containing 0.1% formic acid as solvent A and 80% acetonitrile with 0.1% formic acid as solvent B with a flow rate of 300 nl min$^{-1}$ using the following run conditions over 120 minutes; equilibration at 5% B for 1 min, gradient from 5% B to 25% B in 76 min, isocratic at 25% B for 5 min, gradient from 25% B to 35% B in 10 min, isocratic at 35% B for 5 min, gradient from 35% to 95% B in 8 min, and finally isocratic at 95% B for 15 minutes. The 25 cm length column was maintained at 40 °C, and the 50 cm length column at 45 °C. Full MS spectra were acquired at a resolution of 120,000 at m/z 200 Th, microscans set to 1, AGC target value of 1e6, maximum ion injection time of 25 ms, and scan range of 300-2000 m/z. ddMS2 spectra were acquired at a resolution of 15,000 at m/z 200 Th, microscans set to 1, AGC target value of 5e4, maximum ion injection time of 50 ms, loop count of 15, and collision energy of 30. Lock mass tolerance was set to 10 ppm. A nanospray ion source was used with a spray voltage of 1.7 kV, capillary temperature of 275 °C, and S-lens RF level of 55.

The data were analysed using Proteome Discoverer version 2.4 (Thermo Scientific, USA). A total of 20 raw files were processed as a sample group of 4 (each sample group has 5 fractions). The protein databases were downloaded from Uniprot (UP000000589 for *Mus Musculus* and UP000074855 for *Plasmodium berghei* ANKA, dated April 24, 2024). Data were processed using Sequest HT as a search engine against the Uniprot FASTA databases of *Mus Musculus, Plasmodium berghei*, and a contaminant database. The following parameters were used; maximum trypsin missed cleavage sites was set to 2, precursor mass tolerance of 10 ppm, and fragment mass tolerance of 0.05 Da. The following dynamic modifications were included: oxidation (+15.995 Da at Met), 2-succination (+116.011 Da at Cys), carbamidomethylation (+57.021 Da at Cys), deamidation (+0.984 Da at Asn or Gln), acetylation (+42.011 Da (at N-terminal amino group)), Met-loss (-131.040 Da at N-terminus), and Met-loss+Acetyl (-89.030 Da at N-terminus). Among the detected proteins, a search was made for peptides with at least 2 peptide spectrum matches (PSM) and 2SC modification. Included in the search were also the intensity of precursor ions (available "quan info") and inclusion of only those peptides assigned a confidence score of "high" during their identification (Sequest HT). Among the seven succinated proteins reported in the study, at least 2 PSMs for the modified peptide were present in one of the replicates and a minimum of 1 PSM in the other replicate.

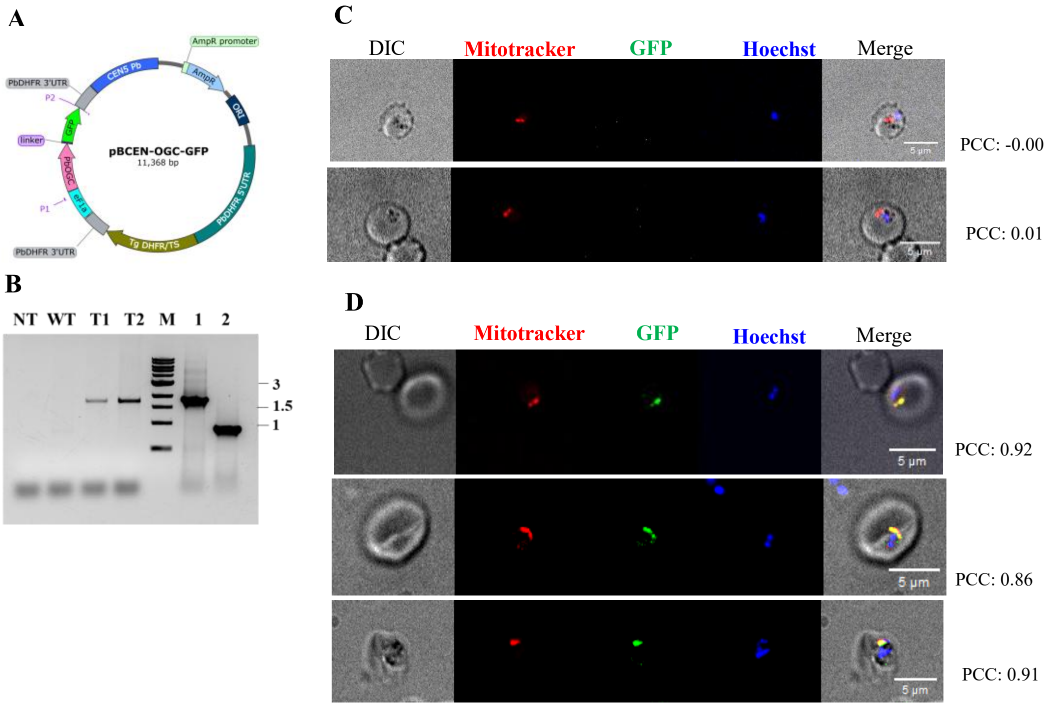


**Figure S1. PbOGC-GFP localizes to the mitochondrion.** (A) A schematic of the PbCEN5-PbOGC-GFP plasmid (generated using SnapGene viewer). P1 and P2 primers used for genotyping are shown in the schematic. (B) Genotyping PbOGC-GFP expressing parasites. NT, no template control; WT, *P. berghei* wild-type genomic DNA; T1 and T2, gDNA from the two clones of PbOGC-GFP expressing parasites; M, molecular weight marker; 1, positive control, PbCEN5-OGC-GFP plasmid; 2, PbCEN5-GFP (empty vector) plasmid as control. If the *ogc-gfp* gene is present, a band at 1.8 kbp is expected with P1 and P2 primers. If the empty vector is present, a band at 863 bp is expected. Both the clones showed a band at 1.8 kbp, thus, confirming the presence of *ogc-gfp*. The band sizes in (B) indicate the size of the markers in kbp. (C) Two representative microscopy images of *Pb* wild-type are shown after Hoescht and Mitotracker staining. (D) Three representative microscopy images of PbOGC-GFP expressing parasites are shown after staining with Hoescht and Mitotracker. The image analyses in (C) and (D) were carried out using ImageJ, and PCC values were obtained using the coloc2 plugin of ImageJ.

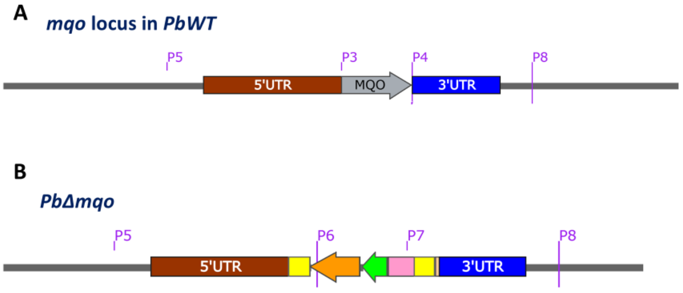


C

| Primers | Verification of | Expected band (bp) | |
|---|---|---|---|
| | | *PbWT* | *PbΔmqo* |
| P3 & P4 | *Pbmqo* ORF | 1563 | --- |
| P5 & P6 | 5′ integration | --- | 4492 |
| P7 & P8 | 3′ integration | --- | 3343 |

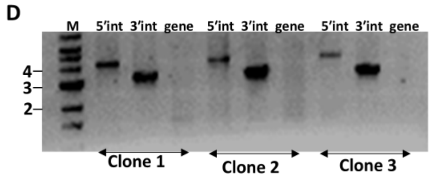


**Figure S2. Genotyping of *Δmqo P. berghei*.** Schematic of (A) intact *Pbmqo* gene locus and (B) *Pbmqo* gene replaced by the selectable marker cassette (hDHFR-yFCU) in *Δmqo*; yellow, PbDHFR 3′UTR; orange, yFCU; green, hDHFR; pink, PbeF1α promoter; tan, 3x HA. The primers used for genotyping by PCR are indicated on the schematics. (C) List of primer pairs used for PCR to confirm integration of the marker cassette at 5′ and 3′ ends, and the absence of *Pbmqo* gene. (D) PCR genotyping of clonal lines of *Δmqo* parasites. Numbers next to the marker lane correspond to the size in kilo base pair (kbp) of the adjacent band. 5′int, 5′ integration; 3′int, 3′ integration; gene, *Pbmqo*. Schematic maps generated using SnapGene.

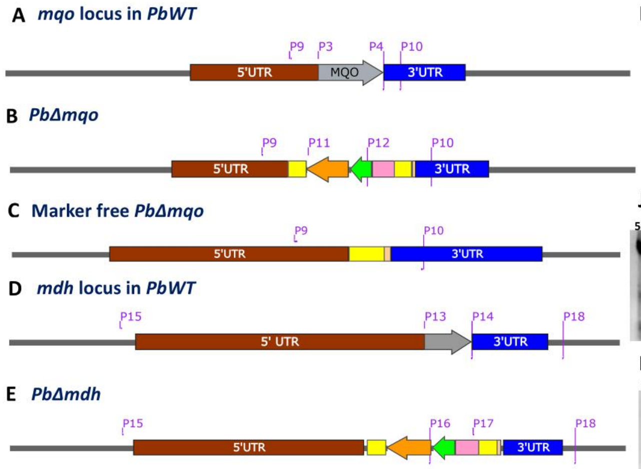


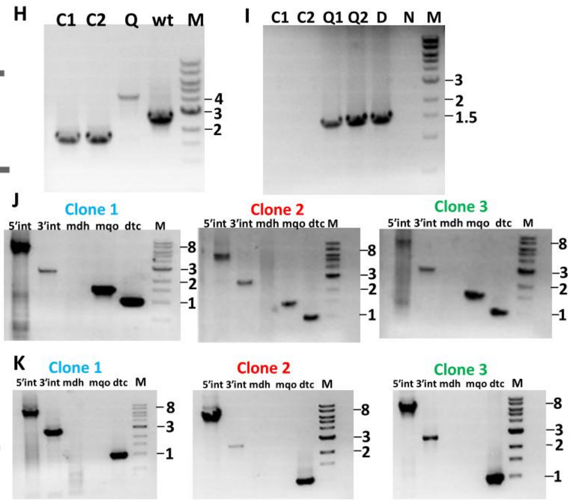


F

| Primers | Verification of | Expected band (bp) | | |
|---|---|---|---|---|
| | | *PbWT* | *PbΔmqo* | Marker free *PbΔmqo* |
| P9 & P10 | Marker excision | 2659 | 4436 | 1648 |
| P11 & P12 | Presence of markers | ---- | 1548 | ---- |

G

| Primers | Verification of | Expected band (bp) | | |
|---|---|---|---|---|
| | | *PbWT* | *PbΔmdh* | *PbΔmqoΔmdh* |
| P3 & P4 | *Pbmqo* ORF | 1563 | 1563 | --- |
| P13 & P14 | *Pbmdh* ORF | 942 | --- | --- |
| P15 & P16 | 5′ integration | --- | 7682 | 7682 |
| P17 & P18 | 3′ integration | --- | 2557 | 2557 |

**Figure S3. Genotyping of *Δmdh* and *ΔmqoΔmdh (Δmm) P. berghei*.** Schematic of the relevant loci of (A) intact *Pbmqo* gene in the WT parasites, (B) *Pbmqo* gene knocked out and replaced by selectable marker cassette, hDHFR-yFCU in *Δmqo* parasites; yellow, PbDHFR 3′UTR; orange, yFCU; green, hDHFR; pink, PbeF1α promoter; tan, 3x HA, (C) selectable marker excised in *Δmqo* parasites, (D) intact *Pbmdh* gene in the WT parasites; grey, *Pbmdh* gene and (E) *Pbmdh* gene knocked out and replaced by selectable marker cassette, hDHFR-yFCU in *Δmdh* or *Δmm* parasites. The primers used for genotyping by PCR are indicated on the schematics and the expected bands are shown in (F) and (G). (H) PCR genotyping using oligonucleotides P9 and P10, and genomic DNA isolated from the clones C1 and C2 obtained by limiting dilution cloning of 5-FC treated *Δmqo* parasites, Lanes correspond to Q, uncloned *Δmqo* parasites before treatment with 5-FC and retaining the drug selectable marker; C1, clone 1; C2, clone 2; wt, *P. berghei* WT; M, 1kbp NEB ladder. PCR analysis show excision of the marker cassette. (I) PCR genotyping using genomic DNA isolated from C1 and C2 clonal lines of *Δmqo* using oligonucleotides P11 and P12 for confirmation of the absence of the marker cassette; C1, clone 1; C2, clone 2; M, 1kbp NEB ladder; Q1 and Q2 are uncloned 5-FC treated *Δmqo* parasites that retain the drug selectable marker, D, *Δdtc* parasites harbouring the selectable marker that was used as control; N, no template. The PCR results in this gel confirm the absence of the selectable marker cassette. (J) PCR genotyping using the genomic DNA isolated from clone 1, clone 2, and clone 3 of *Δmdh* parasites. (K) PCR genotyping using the genomic DNA isolated from clone 1, clone 2, and clone 3 of *Δmm* parasites. The lanes in (J) and (K) are as follows; 5′int, PCR with primers P15 and P16 confirms 5′ integration of the marker cassette; 3′int, PCR with primers P18 and P17 confirms 3′ integration of the marker cassette; mdh, PCR with P13 and P14 for the presence/absence of *Pbmdh* gene; mqo, PCR using primers P3 and P4 for the presence/absence of *Pbmqo* gene; dtc, PCR using primers P19 and P20 (primers are indicated in Fig. S4A for *dtc* gene); M, 1kbp NEB ladder. Numbers next to the marker lane correspond to the size in kbp of the adjacent band. Schematic maps generated using SnapGene.

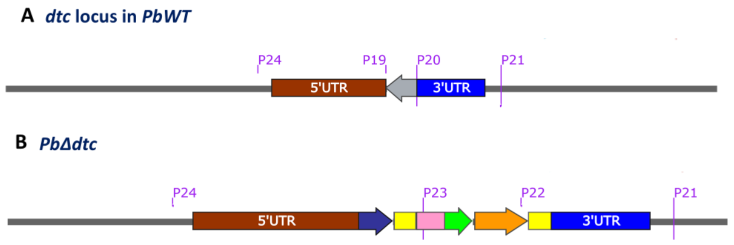


C

| Primers | Verification of | Expected band (bp) | |
|---|---|---|---|
| | | *PbWT* | *PbΔdtc* |
| P19 & P20 | *Pbdtc* ORF | 960 | --- |
| P21 & P22 | 3′ integration | --- | 3149 |
| P23 & P24 | 5′ integration | --- | 5165 |

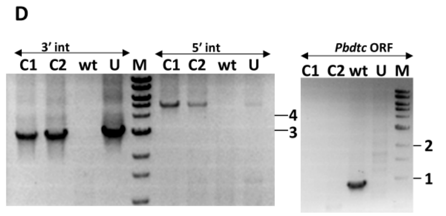


**Figure S4. Genotyping of *Δdtc P. berghei.*** Schematic of (A) intact *Pbdtc* gene locus; grey, *dtc* gene, (B) *Pbdtc* gene replaced by selectable marker cassette, hDHFR-yFCU; yellow, PbDHFR 3′UTR; orange, yFCU; green, hDHFR; pink, PbeF1α promoter; indigo, mcherry. Location of primers used for PCR genotyping are indicated on the schematic map. (C) Oligonucleotides used for genotyping. (D) PCR genotyping of clones of *Δdtc* obtained by limiting dilution. Lane, C1, and C2 are the clones of *Δdtc*; WT, wild-type *P. berghei*, U, uncloned *Δdtc*, M, 1kbp NEB ladder. 3′int, 3′ integration; 5′int, 5′ integration. The band sizes in kbp in panel (D) indicate the size of the marker. Schematic maps were generated using SnapGene.

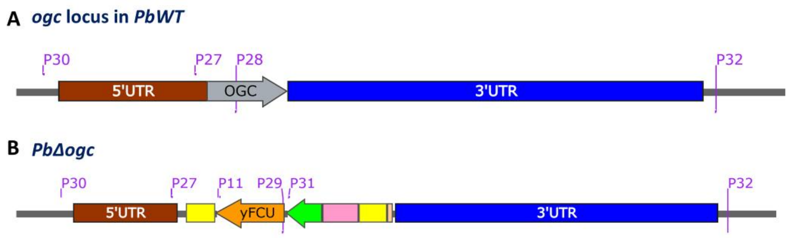


C

| Primers | Verification of | Expected band (bp) | |
|---|---|---|---|
| | | *PbWT* | *PbΔogc* |
| P27 & P28 | *Pbogc* gene | 490 | --- |
| P29 & P30 | 5′ integration | --- | 3575 |
| P31 & P32 | 3′ integration | --- | 7081 |
| P11 & P29 | Presence of markers | --- | 1046 |

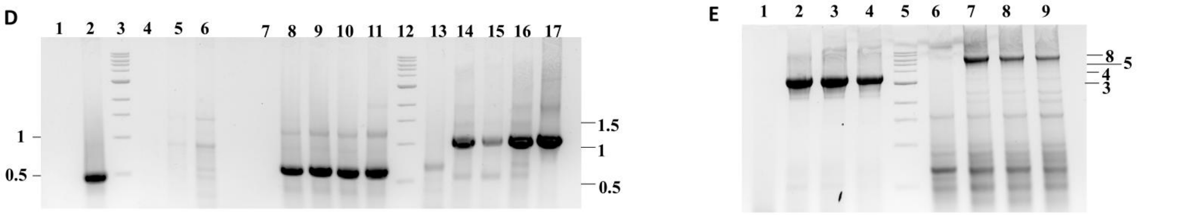


**Figure S5. Genotyping of *Δogc P. berghei.*** Schematic of (A) *P. berghei* wild-type *ogc* gene locus and (B) *P. berghei ogc* gene locus upon knockout of the *ogc* gene and integration of the selection markers (hDHFR and yFCU); yellow, PbDHFR 3′UTR; orange, yFCU; green, hDHFR; pink, PbeF1α promoter; tan, 3x HA. The primers used for genotyping are shown in the schematics and the expected bands are indicated in (C). (D) and (E) Genotyping of the three clonal lines of *Δogc* parasites. (D) Primers P27 and P28 were used to check the presence of *ogc* gene (lanes 1-6), PCR validation for PBANKA_100510 gene as a control (lanes 7-11), primers P11 and P29 were used to check the presence of selection marker (lane 13-17). Lanes 1 and 7, no template control; lanes 2, 8, and 13, gDNA of PbWT; lanes 4-6, 9-11, 14-16 are C1, C2, and C3 clonal lines of *Δogc*, obtained after limiting dilution cloning; lane 17, *ogc* knockout construct as control; lanes 3 and 12, molecular weight marker. *Δogc* C1 and C2 clonal lines did not show any band at 490 bp, confirming the absence of *ogc* gene. All three clonal lines showed the presence of the marker. (E) Primers P29 and P30 were used to check the 5′ integration (lanes 1-4) and primers P31 and P32 were used to check the 3′ integration (lanes 6-9). Lane 1 and 6, gDNA of PbWT; lanes 2-4 and 7-9 are C1, C2, and C3 clonal lines of *Δogc*, obtained after limiting dilution cloning; lane 5, molecular weight marker. All three clones showed the expected bands of 5′ and 3′ integration. The band sizes in kbp in panels (D) and (E) indicate the size of the marker. All the schematic maps were generated using SnapGene viewer.

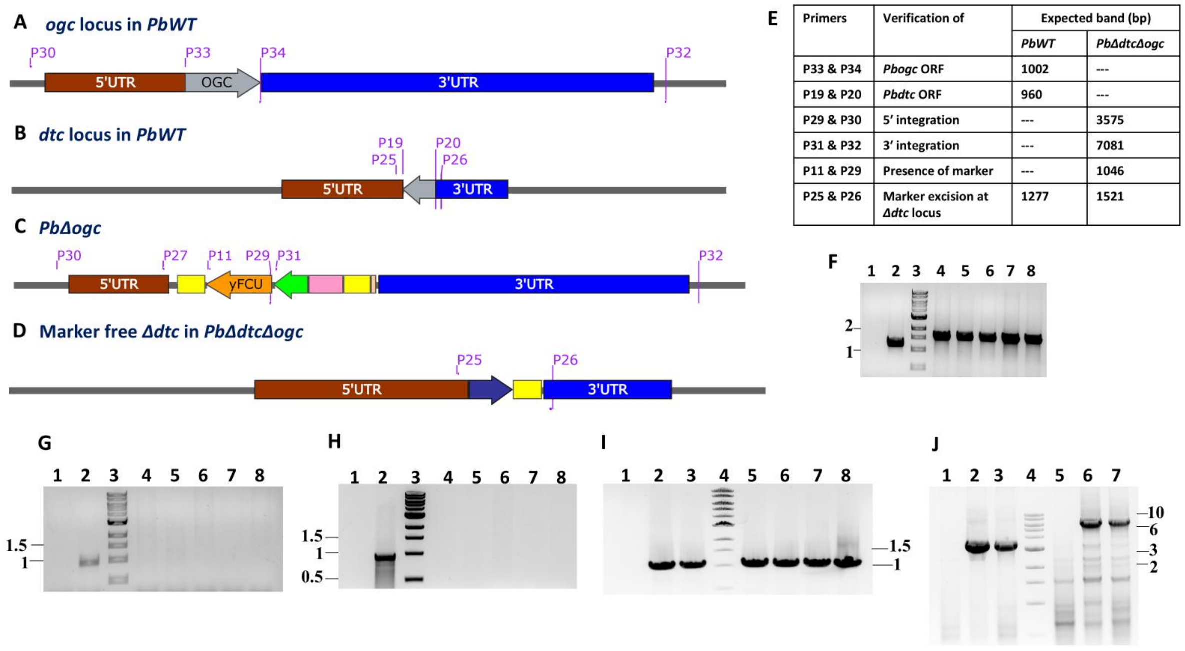


| Primers | Verification of | Expected band (bp) | |
|---|---|---|---|
| | | *PbWT* | *PbΔdtcΔogc* |
| P33 & P34 | *Pbogc* ORF | 1002 | --- |
| P19 & P20 | *Pbdtc* ORF | 960 | --- |
| P29 & P30 | 5′ integration | --- | 3575 |
| P31 & P32 | 3′ integration | --- | 7081 |
| P11 & P29 | Presence of marker | --- | 1046 |
| P25 & P26 | Marker excision at *Δdtc* locus | 1277 | 1521 |

**Figure S6. Genotyping *ΔdtcΔogc* (*Δod*) *P. berghei*.** Schematic of (A) *P. berghei* wild-type *ogc* gene locus, (B) *P. berghei* wild-type *dtc* gene locus; grey, *dtc* gene, (C) *P. berghei ogc* gene locus upon knockout and integration of the selection markers in *Δod* and (D) *P. berghei* marker free *Δdtc* locus in *Δod*; yellow, PbDHFR 3′UTR; orange, yFCU; green, hDHFR; pink, PbeF1α promoter; tan, 3x HA; indigo, mcherry. The primers used for genotyping are shown in the schematics and the expected bands are indicated in (E). (F) PCR verification of marker free *Δdtc* locus in *Δod* using primers P25 and P26. The PCR confirms the marker free *Δdtc* locus in *Δod*. (G) PCR verification of *Δod* for the absence of *ogc* gene using primers P33 and P34. The PCR confirms the absence of the *ogc* gene in *Δod*. (H) PCR verification of *Δod* for the absence of *dtc* gene using primers P19 and P20. The PCR confirms the absence of the *dtc* gene. Lanes in (F) – (H): Lane 1, no template control; Lane 2, *P. berghei* wild-type genomic DNA; lane 3, molecular weight marker, lanes 4 and 5, genomic DNA of drug-resistant *Δod* parasites (unclonal); lanes 6-8, genomic DNA of *Δod* parasites of C1, C2, and C3 clones, respectively, obtained after limiting dilution cloning. (I) PCR verification of *Δod* parasites for the presence of selection marker using primers P11 and P29. Lane 1, *P. berghei* wild-type genomic DNA; lane 2 and 3, genomic DNA of drug-resistant *Δod* parasites (unclonal), lane 4, molecular weight marker; lanes 5-7, genomic DNA of *Δod* parasites of C1, C2, and C3 clones, respectively, obtained after limiting dilution cloning; lane 8, *dtc* knockout construct as positive control. The *Δod* parasites showed the expected band confirming the presence of markers. (J) PCR verification of *Δod* parasites for 5′ and 3′ integration at the *ogc* locus. Lanes 1-3 for 5′ integration using primers P29 and P30; lanes 5-7 for 3′ integration using primers P31 and P32; lane 1 and 5, *P. berghei* wild-type genomic DNA; lane 2, 3, 6, and 7, genomic DNA of *Δod* parasites; lane 4, molecular weight marker. *Δod* parasites showed the expected bands for 5′ integration and 3′ integration. The band sizes in (F) - (J) figures indicate the size of the marker. All the schematic maps were generated using SnapGene viewer.

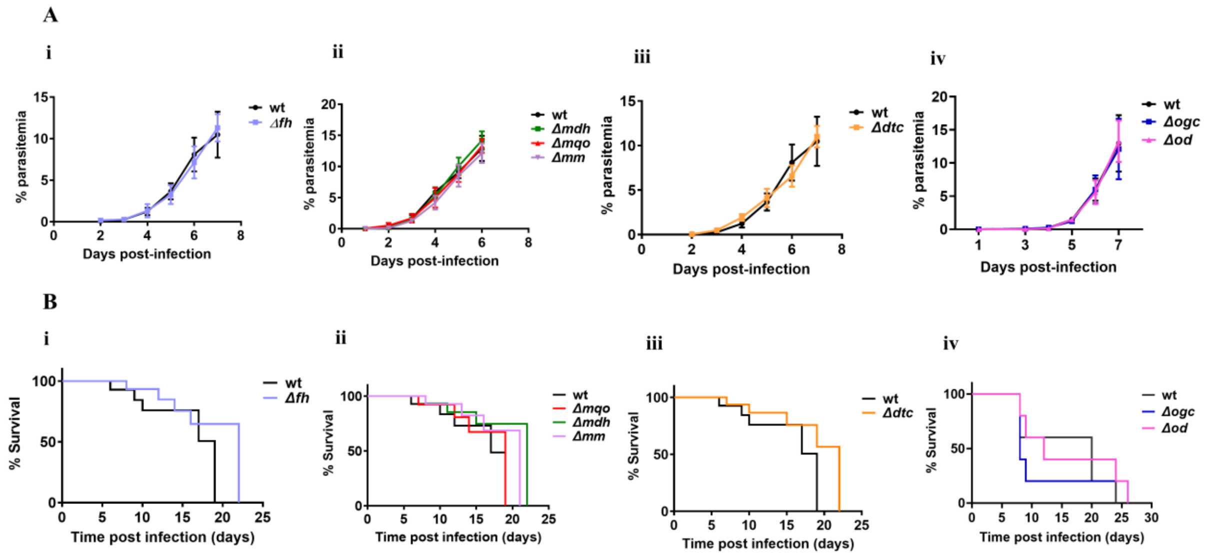


**Figure S7. Comparison of the growth curves and percent survival of mice infected with *P. berghei* WT and knockouts.** (A) Comparison of the growth curves of the asexual intraerythrocytic stages between WT and the seven knockouts. Growth rate of (i) *Δfh*, (ii) *Δmdh*, *Δmqo*, and *Δmm*, (iii) *Δdtc*, and (iv) *Δogc* and *Δod* is plotted with their respective WT. The parasitemia is represented as mean and error bars indicate standard deviation (SD); N = 3-5 for wild-type, N = 4-5 for knockouts. Statistical analysis using Student's unpaired t-test did not show any significant difference in the growth rate between wild-type and the knockouts. (B) The percentage survival of C57BL/6 mice infected with *P. berghei* WT and the knockouts. The curve for the survivability of mice infected with knockouts was found to be not significantly different from that of the curve for the mice infected with wild-type parasites. Statistical analysis using the Log-rank (Mantel-Cox) test showed a p-value of 0.2952, 0.3757, 0.1732, 0.1925, 0.2261, 0.7676, and 0.4639 and the Gehan-Breslow-Wilcoxon test showed a p-value of 0.6443, 0.5339, 0.2851, 0.2676, 0.4078, 0.866, and 0.3496 between the *PbWT* and *Δfh*, *Δmqo*, *Δmdh*, *Δmqo/Δmdh*, *Δdtc*, *Δogc*, *Δod* line. N= 3-5 for WT and N=4-5 for knockouts.

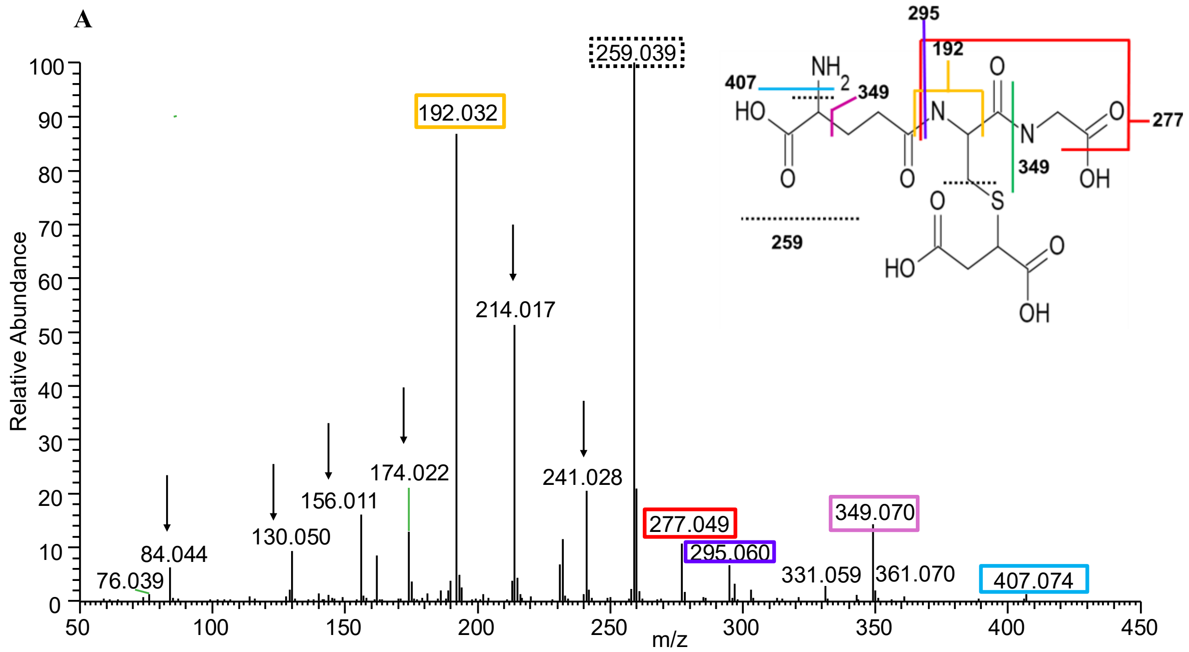
A
Relative Abundance
m/z
259.039
192.032
214.017
241.028
174.022
156.011
130.050
84.044
76.039
277.049
295.060
349.070
331.059
361.070
407.074
295
192
407
349
277
259
HO
OH


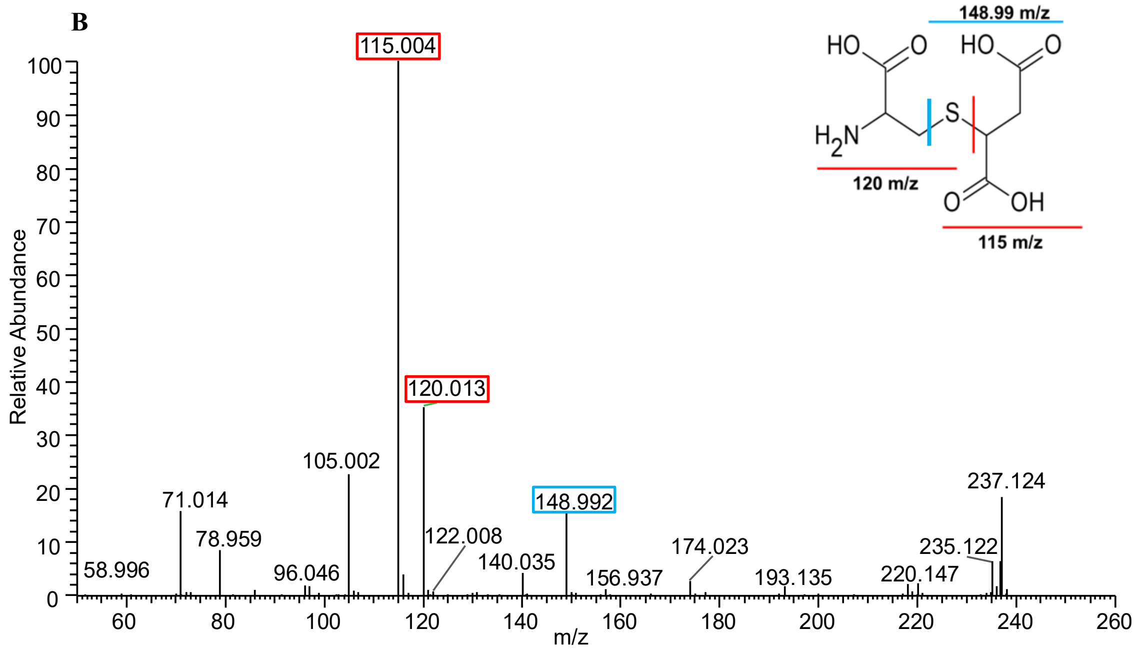
B
Relative Abundance
m/z
115.004
120.013
105.002
71.014
78.959
58.996
96.046
122.008
140.035
148.992
156.937
174.023
193.135
220.147
235.122
237.124
148.99 m/z
120 m/z
115 m/z

**C**

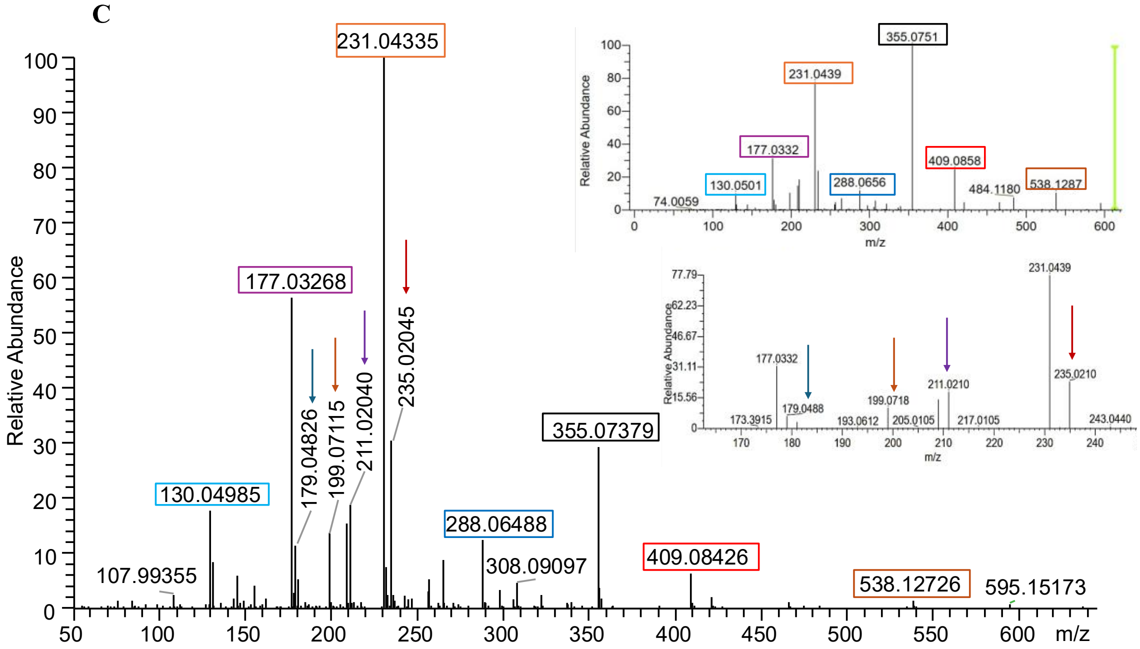


**Figure S8. MS/MS spectra of succ-GSH, 2SC, and GSSG metabolites**. (A) The MS/MS generated fragmentation pattern obtained for the precursor ion of m/z 424.1017 in positive polarity corresponding to succ-GSH. The inset shows the structure of succ-GSH. The fragmentation occurring at different bonds are indicated by different colours and the corresponding ions boxed in the same colour in the MS/MS spectrum are shown. All the expected ions (16) were observed for the metabolite. The other ions corresponding to 241, 214, 174, 156, 130, and 84 m/z arising from succ-GSH fragmentation were also observed earlier (16). These fragments are not shown in the structure (inset) for clarity and are indicated by arrows. (B) The MS/MS generated fragmentation pattern obtained for the precursor ion of m/z 236.0233 in negative polarity corresponding to 2SC. The inset shows the structure of 2SC. Fragmentation at the bond indicated in red generates the fragment ions of m/z 120 and 115 m/z. Fragmentation at the bond indicated in blue generates the fragment ion of m/z 148.99 in negative polarity. The MS/MS fragment ion masses confirm the parent ion to be 2SC. The structures in inset of (A) and (B) were generated using ChemSketch (17). (C) The MS/MS generated fragmentation pattern obtained for the precursor ion of m/z 613.1584 in positive polarity corresponding to GSSG. The two insets show the expected fragmentation patterns for GSSG from HCD fragmentation in positive polarity obtained from the mzCloud database (https://www.mzcloud.org/ HighChem LLC). The boxed ions and ions highlighted with an arrow in the spectrum obtained from this study are those that correspond to that deposited in the mzcloud database. These are indicated in the same colour across the main panel and the insets.

**A**

**B**

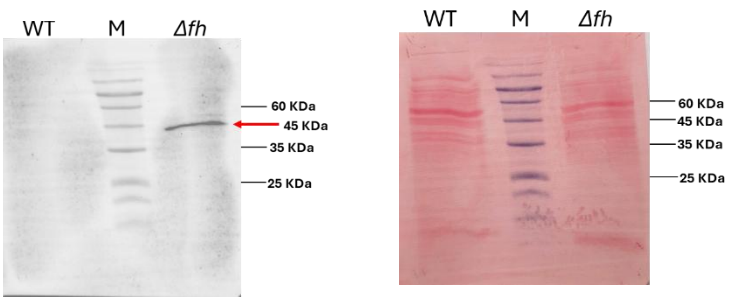


**Figure S9. Presence of a succinated protein in *fh* knockout gametocytes.** (A) Western blot with Anti-2SC antibody detected succinated protein/s at 45 KDa (indicated by a red arrow) only in the lysate of the *fh* knockout gametocytes. Lane M, pre-stained protein marker. The sizes of the marker bands are indicated on the right. Equal total protein concentrations were loaded for the WT and *Δfh*. (B) Ponceau stain of the blot in (A) did not show discrepancies between WT and *fh* knockout in loading or transfer of the protein bands.

**A**

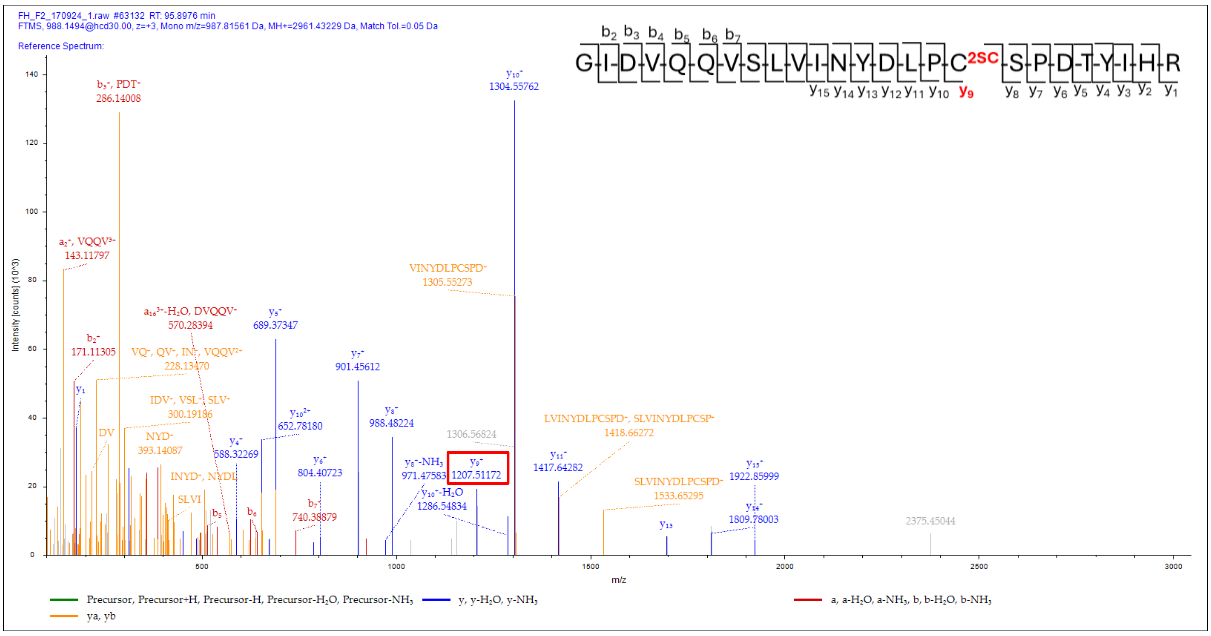

**B**

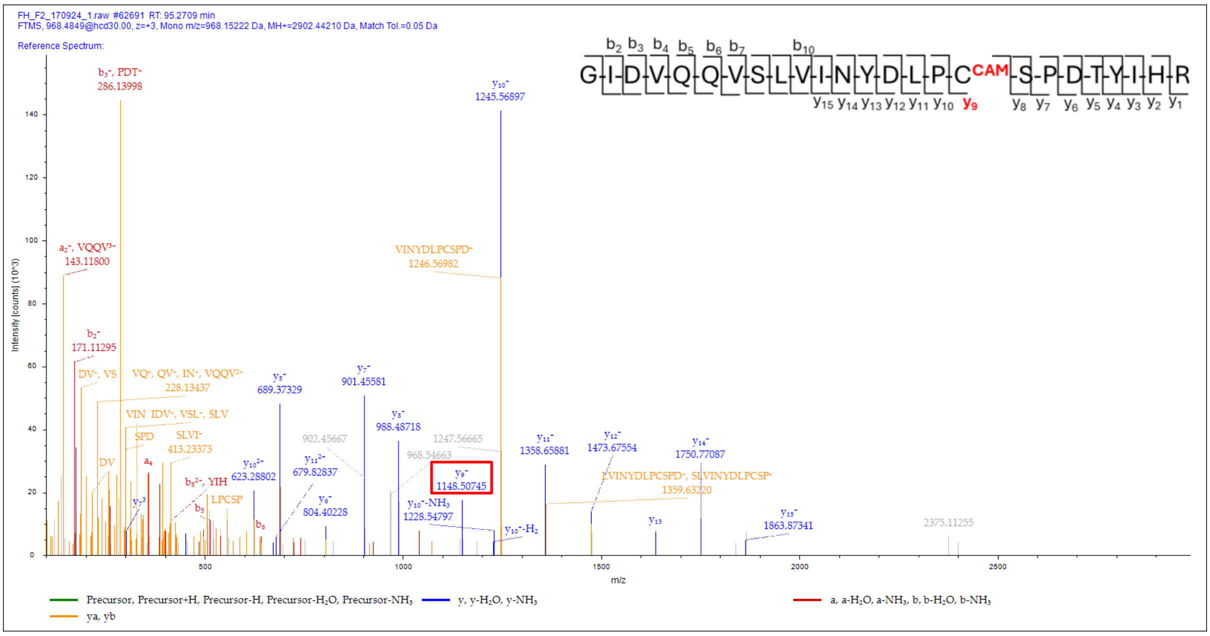


**C**

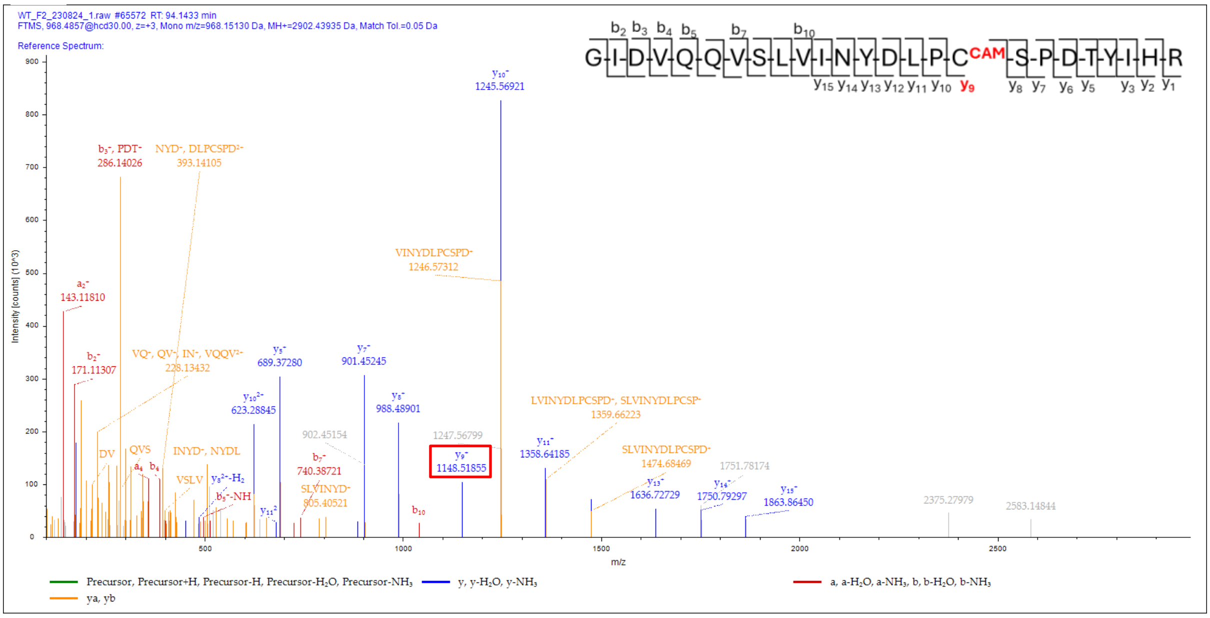


**Figure S10. MS/MS spectra of peptide with sequence GIDVQQVSLVINYDLPCSPDTYIHR belonging to RNA helicase.** (A) MS/MS of peptide of m/z 988.1494 (z=3) from *Δfh*. The y9 ion of m/z 1207.51172 corresponding to 2SC is highlighted in a red box. (B) MS/MS of peptide of m/z 968.4849 (z=3) from *Δfh*. The y9 ion of m/z 1148.50745 corresponding to cysteine-carbamidomethylation (Cys-CAM) is highlighted in a red box. (C) MS/MS of peptide of m/z 968.4857 (z=3) from WT. The y9 ion of m/z 1148.51855 corresponding to Cys-CAM is highlighted in a red box. The WT did not have the corresponding peptide with the cysteinyl residue succinated.

A

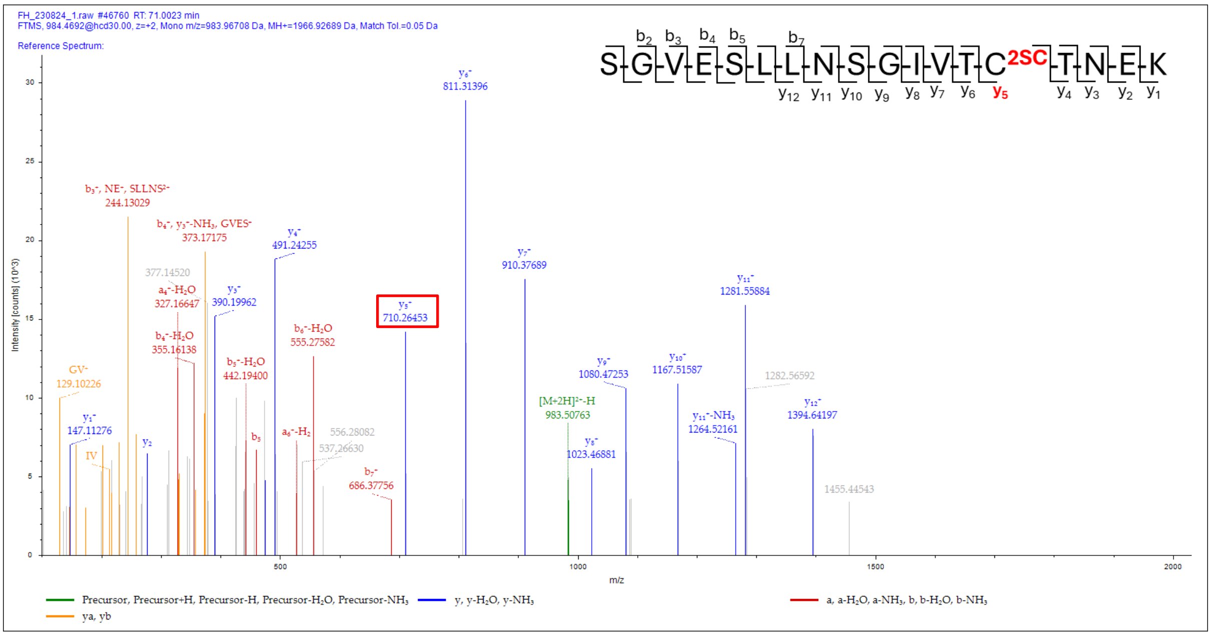

FH_230824_1.raw #46760 RT: 71.0023 min
FTMS, 984.4692@hcd30.00, z=+2, Mono m/z=983.96708 Da, MH+=1966.92689 Da, Match Tol.=0.05 Da
Reference Spectrum:
Intensity [counts] (10^3)
m/z
y5+ 710.26453
y6+ 811.31396
y7+ 910.37689
y11+ 1281.55884
y12+ 1394.64197
[M+2H]2+-H 983.50763
Precursor, Precursor+H, Precursor-H, Precursor-H2O, Precursor-NH3
y, y-H2O, y-NH3
a, a-H2O, a-NH3, b, b-H2O, b-NH3
ya, yb


B

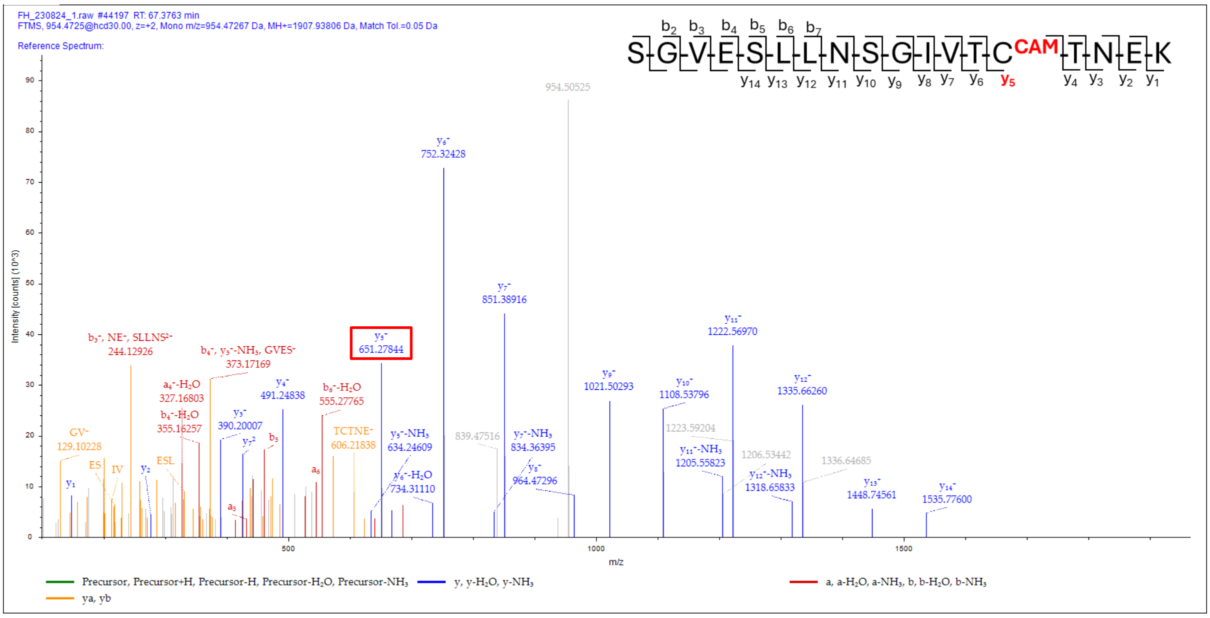

FH_230824_1.raw #44197 RT: 67.3763 min
FTMS, 954.4725@hcd30.00, z=+2, Mono m/z=954.47267 Da, MH+=1907.93806 Da, Match Tol.=0.05 Da
Reference Spectrum:
Intensity [counts] (10^3)
m/z
y5+ 651.27844
y6+ 752.32428
y7+ 851.38916
y11+ 1222.56970
954.50525
Precursor, Precursor+H, Precursor-H, Precursor-H2O, Precursor-NH3
y, y-H2O, y-NH3
a, a-H2O, a-NH3, b, b-H2O, b-NH3
ya, yb

**C**

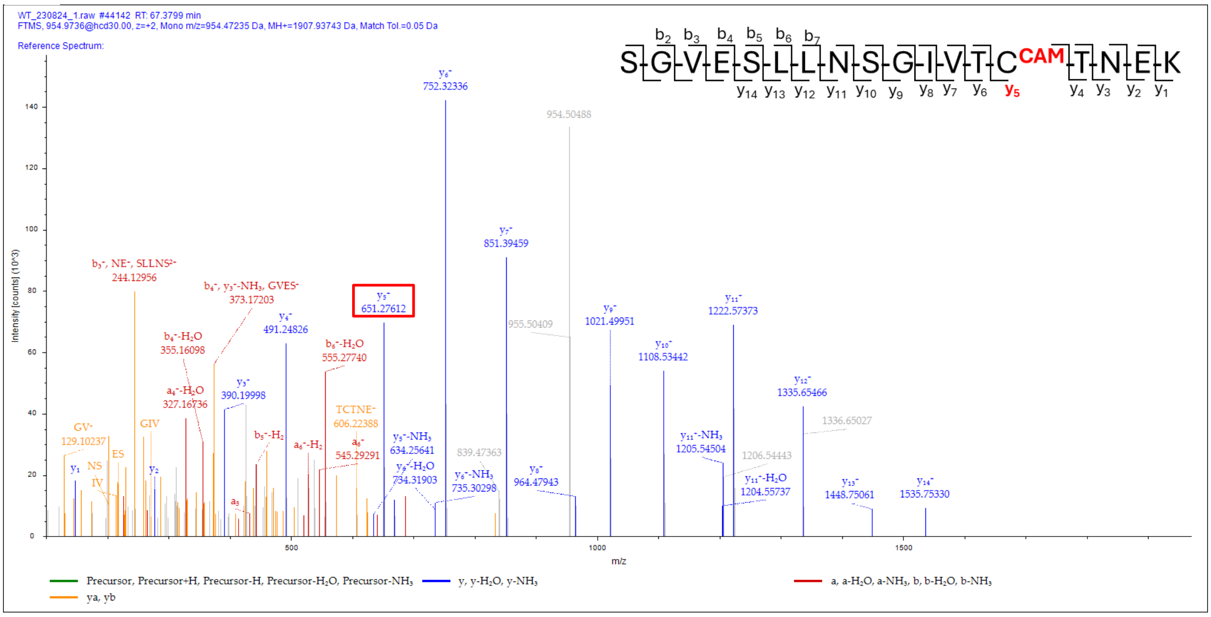


**Figure S11. MS/MS spectra of peptide with sequence SGVESLLNSGIVTCTNEK belonging to tRNA import protein.** (A) MS/MS of peptide of m/z 984.4692 (z=2) from *Δfh*. The y5 ion of m/z 710.26453 corresponding to 2SC is highlighted in a red box. (B) MS/MS of peptide of m/z 954.4725 (z=2) from *Δfh*. The y5 ion of m/z 651.27844 corresponding to Cys-CAM is highlighted in a red box. (C) MS/MS of peptide of m/z 954.9736 (z=2) from WT. The y5 ion of m/z 651.27612 corresponding to Cys-CAM is highlighted in a red box. The WT did not have the corresponding peptide with the cysteinyl residue succinated.

**A**

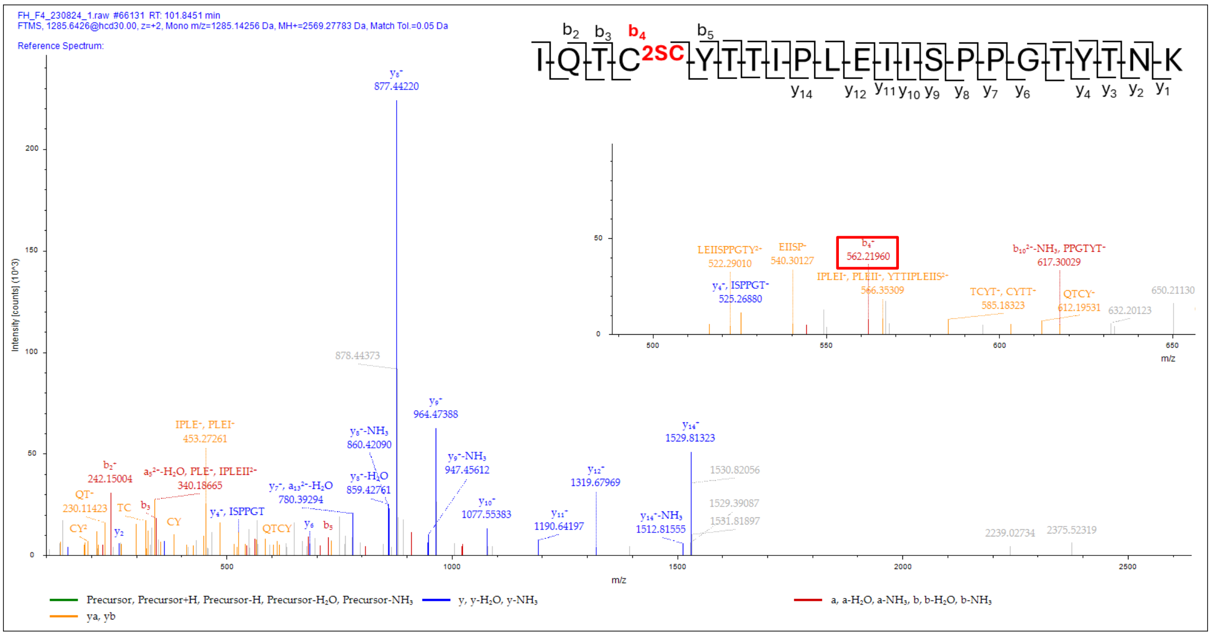

B

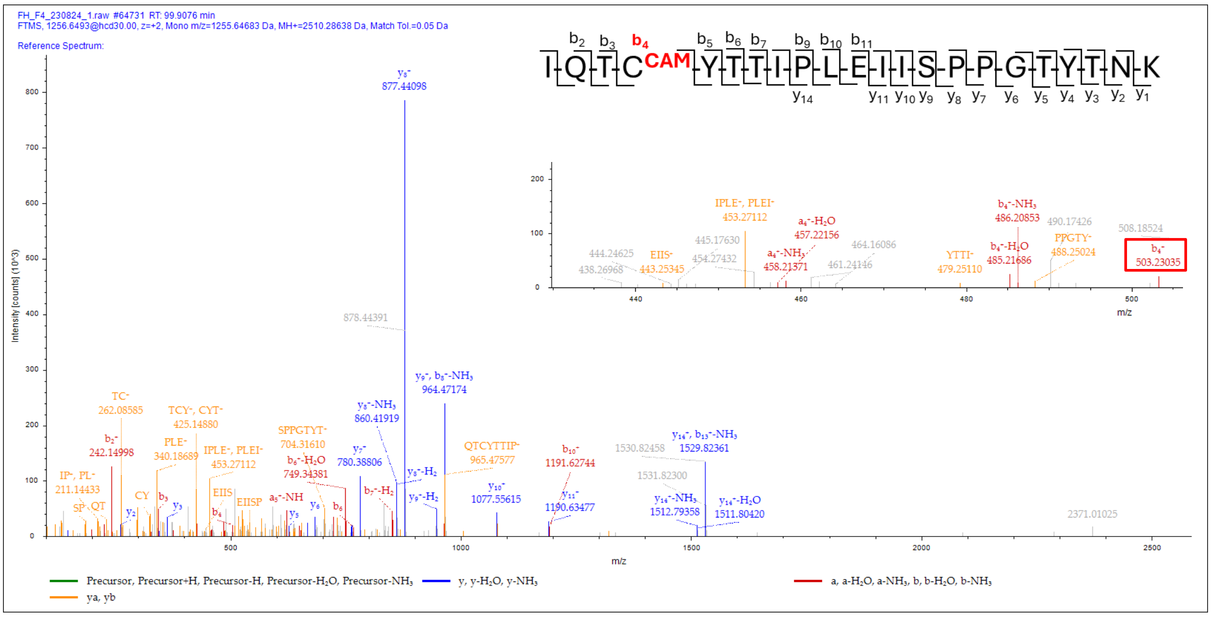


C

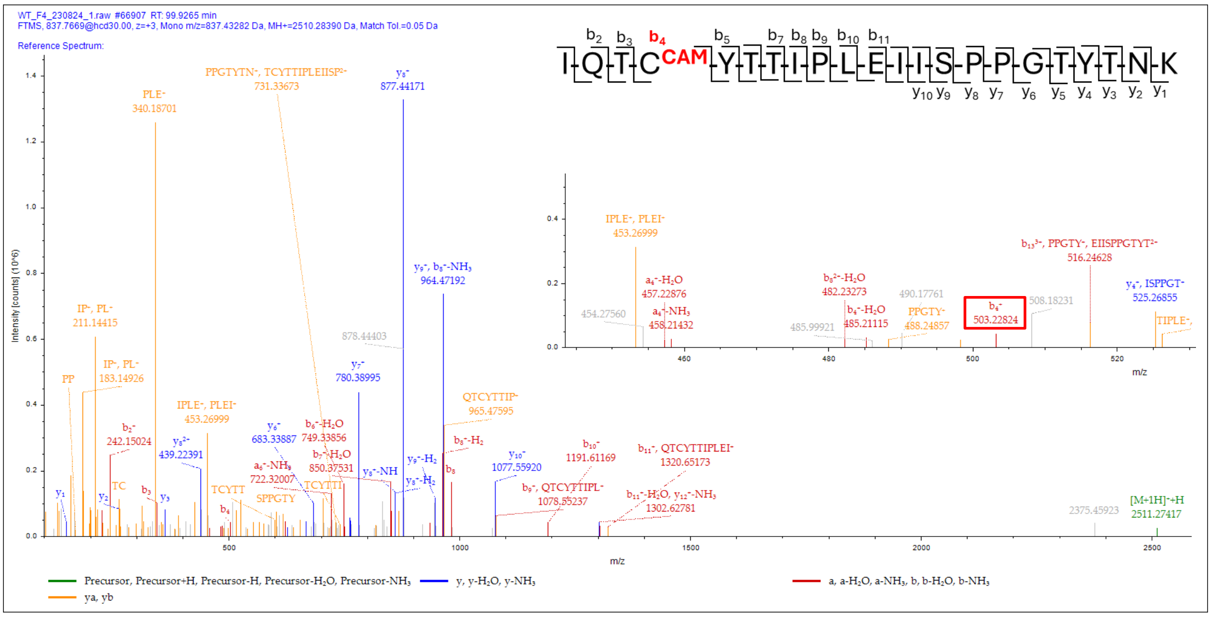


**Figure S12. MS/MS spectra of peptide with sequence IQTCYTTIPLEIISPPGTYTNK belonging to Heat shock protein 110.** (A) MS/MS of peptide of m/z 1285.6426 (z=2) from *Δfh*. The b4 ion of m/z 562.2196 corresponding to 2SC is highlighted in a red box in the inset. (B) MS/MS of peptide of m/z 1256.6493 (z=2) from *Δfh*. The b4 ion of m/z 503.23035 corresponding to Cys-CAM is highlighted in a red box in the inset. (C) MS/MS of peptide of m/z 837.7669 (z=3) from WT. The y5 ion of m/z 503.22824 corresponding to Cys-CAM is highlighted in a red box in the inset. The inset in all three panels shows a portion of axis zoomed in. The WT did not have the corresponding peptide with the cysteinyl residue succinated.

**A**

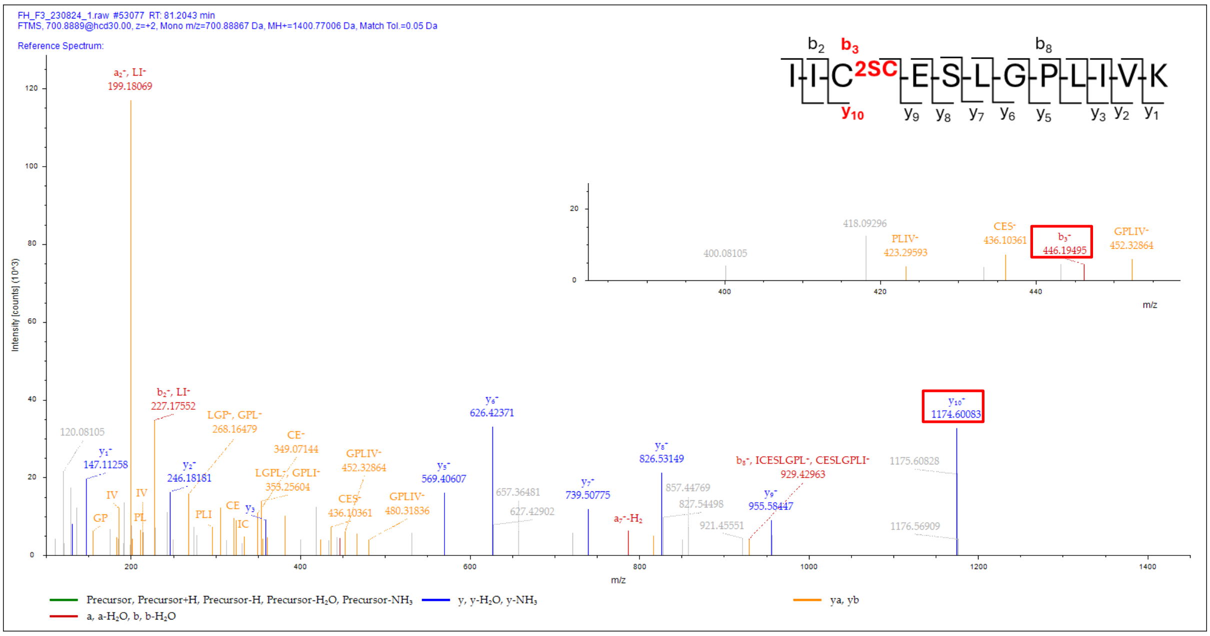

FH_F3_230824_1.raw #53077 RT: 81.2043 min
FTMS, 700.8889@hcd30.00, z=+2, Mono m/z=700.88867 Da, MH+=1400.77006 Da, Match Tol.=0.05 Da
Reference Spectrum:
I-I-C 2SC -E-S-L-G-P-L-I-V-K
b2 b3 b8
y10 y9 y8 y7 y6 y5 y3 y2 y1
Intensity [counts] (10^3)
m/z
a2+, LI+ 199.18069
b2+, LI+ 227.17552
LGP+, GPL+ 268.16479
CE+ 349.07144
GPLIV+ 452.32864
y5+ 569.40607
y6+ 626.42371
y7+ 739.50775
y8+ 826.53149
b8+, ICESLGPL+, CESLGPLI+ 929.42963
y9+ 955.58447
y10+ 1174.60083
1175.60828
1176.56909
120.08105
y1+ 147.11258
y2+ 246.18181
LGPL+, GPLI+ 353.25604
CES+ 436.10361
GPLIV+ 480.31836
657.36481
627.42902
857.44769
827.54498
921.45551
a8+-H2
400.08105
418.09296
PLIV+ 423.29593
CES+ 436.10361
b3+ 446.19495
GPLIV+ 452.32864
Precursor, Precursor+H, Precursor-H, Precursor-H2O, Precursor-NH3
y, y-H2O, y-NH3
ya, yb
a, a-H2O, b, b-H2O


**B**

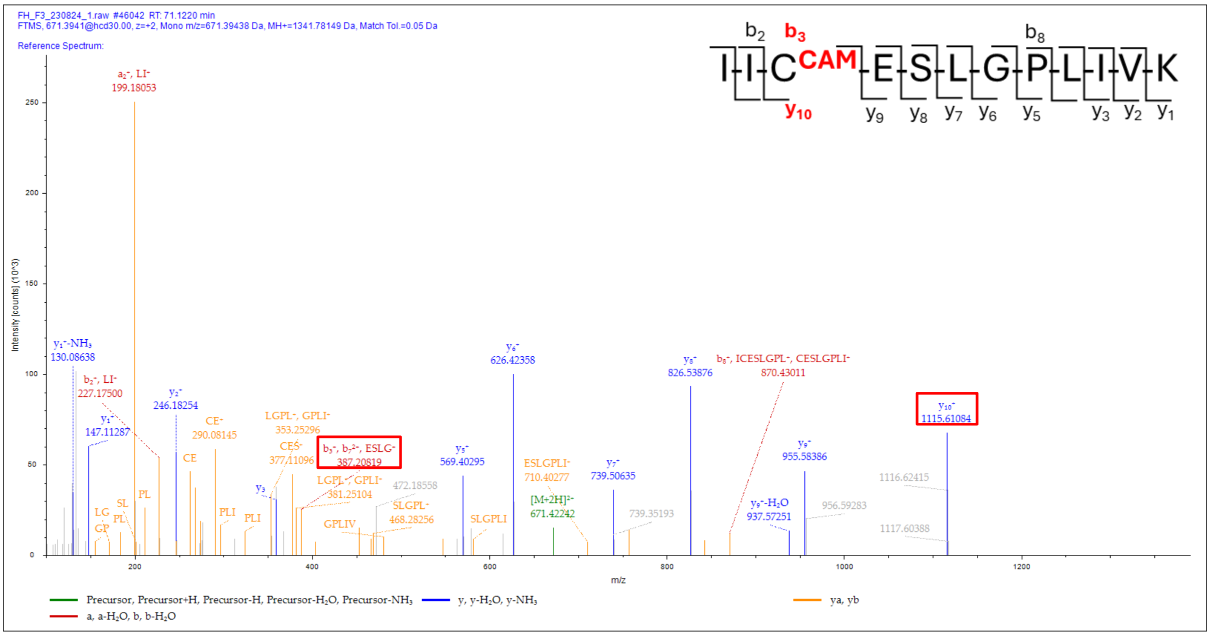

FH_F3_230824_1.raw #46042 RT: 71.1220 min
FTMS, 671.3941@hcd30.00, z=+2, Mono m/z=671.39438 Da, MH+=1341.78149 Da, Match Tol.=0.05 Da
Reference Spectrum:
I-I-C CAM -E-S-L-G-P-L-I-V-K
b2 b3 b8
y10 y9 y8 y7 y6 y5 y3 y2 y1
Intensity [counts] (10^3)
m/z
a2+, LI+ 199.18053
y1+-NH3 130.08638
b2+, LI+ 227.17500
y1+ 147.11287
y2+ 246.18254
CE+ 290.08145
LGPL+, GPLI+ 353.25296
CES+ 377.11096
b3+, b7+2+, ESLG+ 387.20819
LGPL+, GPLI+ 381.25104
472.18558
SLGPL+ 468.28256
y5+ 569.40295
y6+ 626.42358
ESLGPLI+ 710.40277
[M+2H]2+ 671.42242
y7+ 739.50635
739.35193
y8+ 826.53876
b8+, ICESLGPL+, CESLGPLI+ 870.43011
y9+ 955.58386
y9+-H2O 937.57251
956.59283
y10+ 1115.61084
1116.62415
1117.60388
Precursor, Precursor+H, Precursor-H, Precursor-H2O, Precursor-NH3
y, y-H2O, y-NH3
ya, yb
a, a-H2O, b, b-H2O

**C**

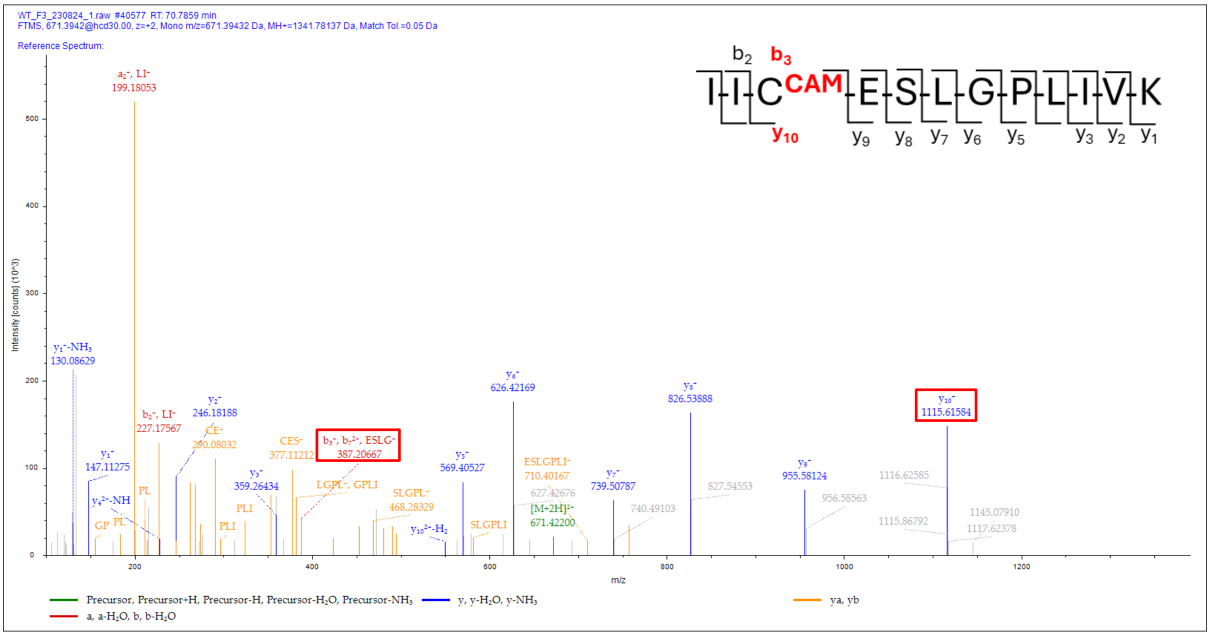


**Figure S13. MS/MS spectra of peptide with sequence IICESLGPLIVK belonging to CUGBP Elav-like family member 2.** (A) MS/MS of peptide of m/z 700.8889 (z=2) from *Δfh*. The b3 ion of m/z 446.19495 and y10 ion of m/z 1174.60083 corresponding to 2SC are highlighted in red boxes. The inset shows a portion of axis zoomed in. (B) MS/MS of peptide of m/z 671.3941 (z=2) from *Δfh*. The b3 ion of m/z 387.20819 and y10 ion of m/z 1115.61084 corresponding to Cys-CAM are highlighted in red boxes. (C) MS/MS of peptide of m/z 671.3942 (z=2) from WT. The b3 ion of m/z 387.20667 and y10 ion of m/z 1115.61584 corresponding to Cys-CAM are highlighted in red boxes. The WT did not have the corresponding peptide with the cysteinyl residue succinated.

**A**

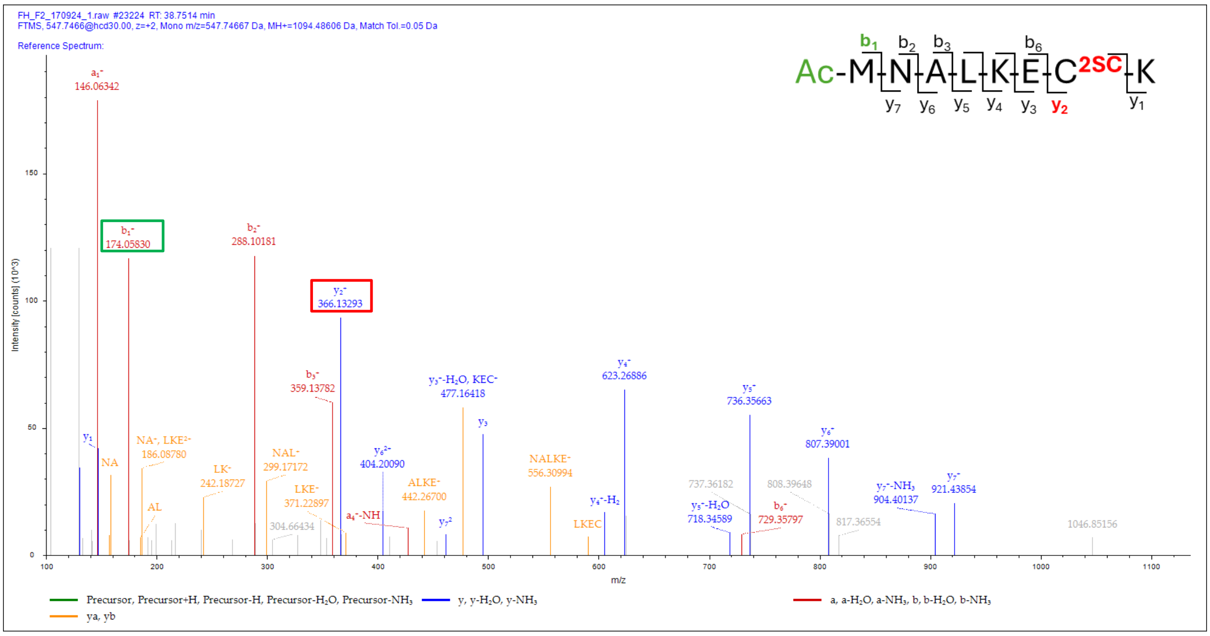

**B**

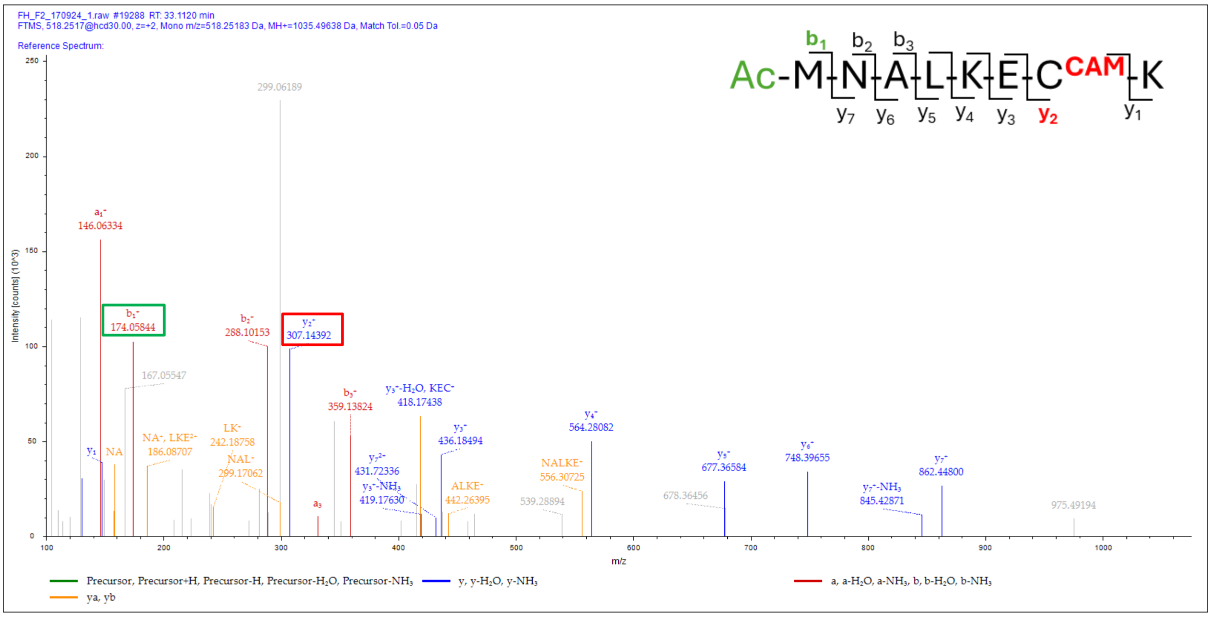


**C**

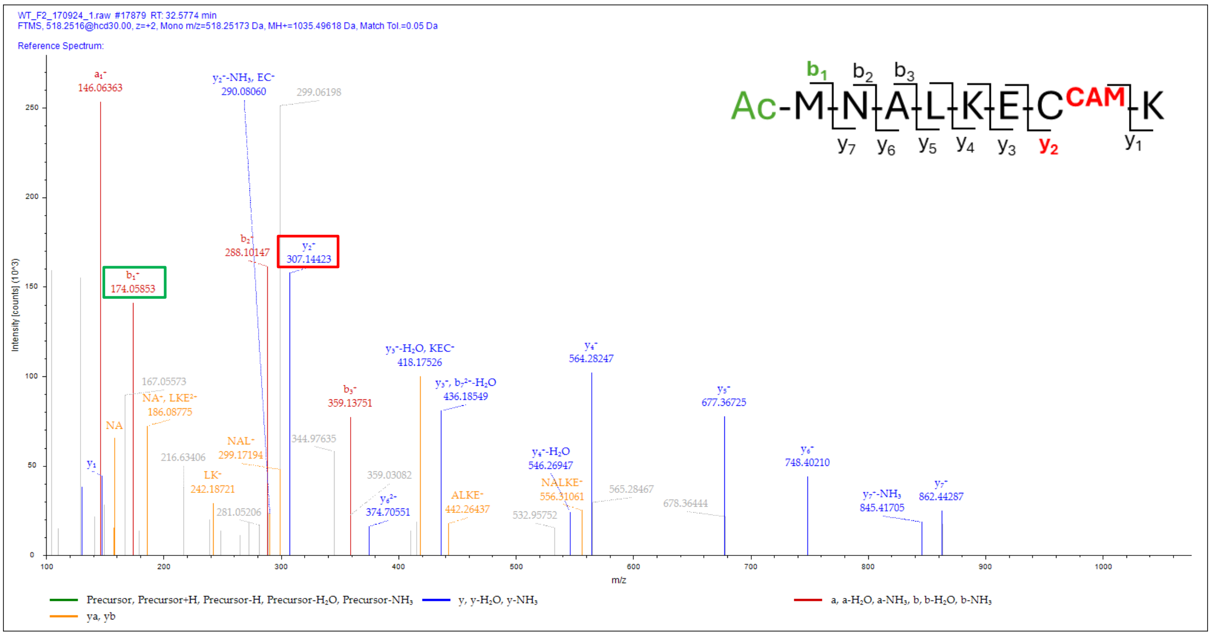


**Figure S14. MS/MS spectra of peptide with sequence MNALKECK belonging to Chromatin assembly factor 1 protein WD40 domain.** (A) MS/MS of peptide of m/z 547.7466 (z=2) from *Δfh*. The y2 ion of m/z 366.13293 corresponding to 2SC and b1 ion of m/z 174.0583 corresponding to N-terminal methionine acetylation are highlighted in red and green boxes, respectively. (B) MS/MS of peptide of m/z 518.2517 (z=2) from *Δfh*. The y2 ion of m/z 307.14392 corresponding to Cys-CAM and b1 ion of m/z 174.05844 corresponding to N-terminal methionine acetylation are highlighted in red and green boxes, respectively. (C) MS/MS of peptide of m/z 518.2516 (z=2) from WT. The y2 ion of m/z 307.14423 corresponding to Cys-CAM and b1 ion of m/z 174.05853 corresponding to N-terminal methionine acetylation are highlighted in red and green boxes, respectively. The WT did not have the corresponding peptide with the cysteinyl residue succinated.

**A**

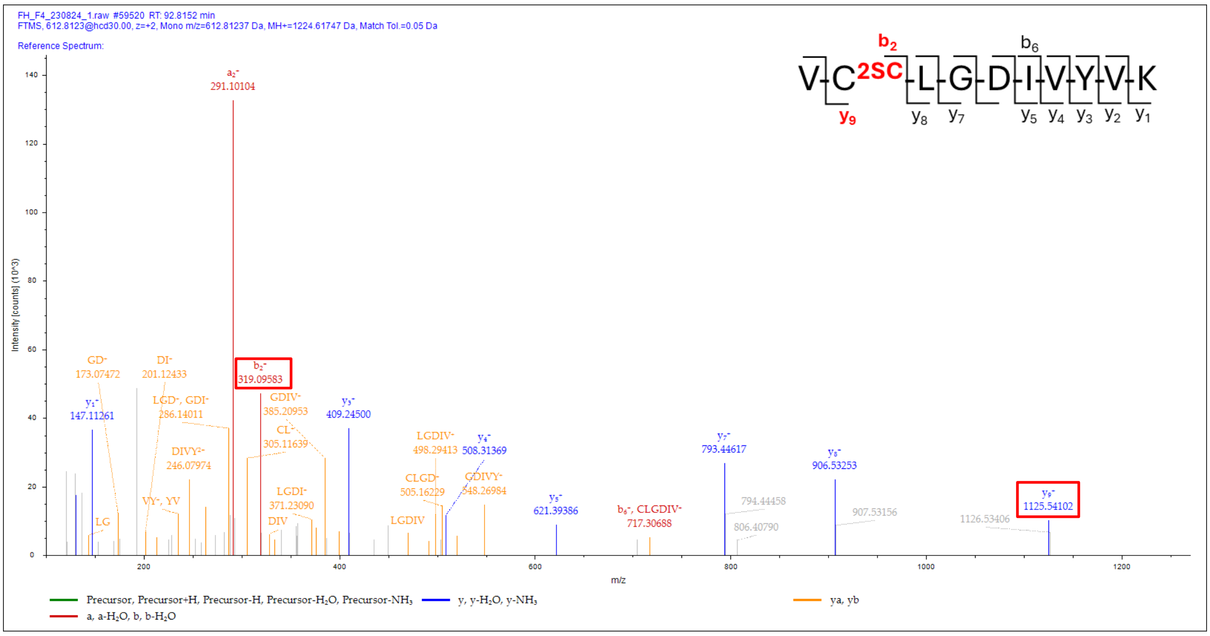


**B**

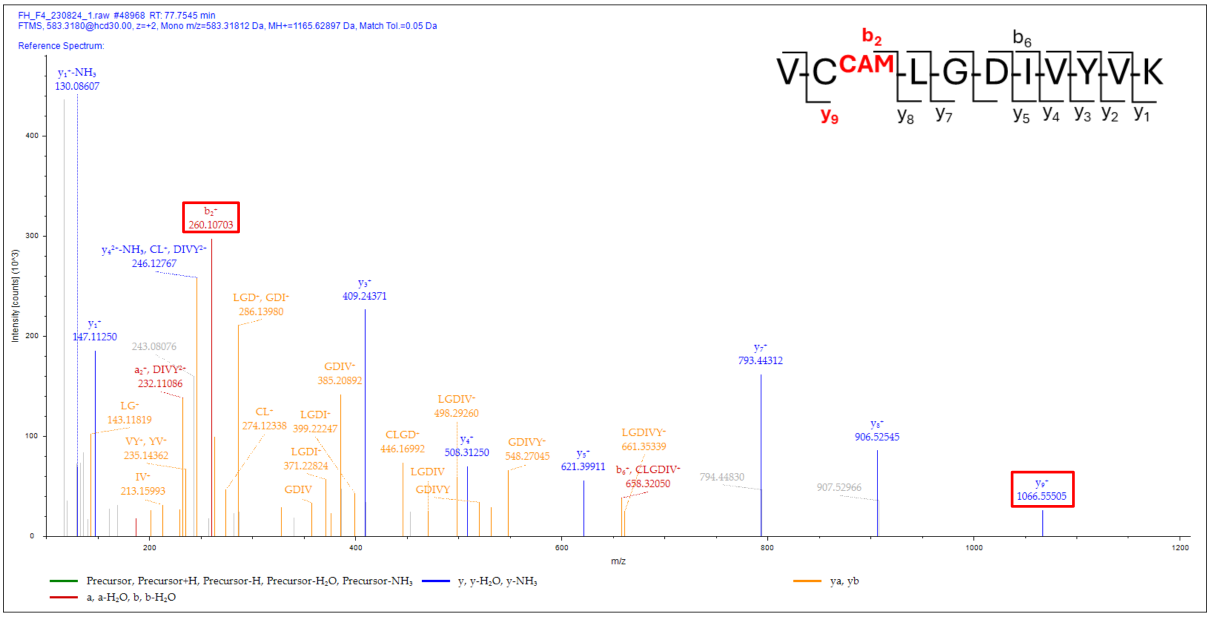

C

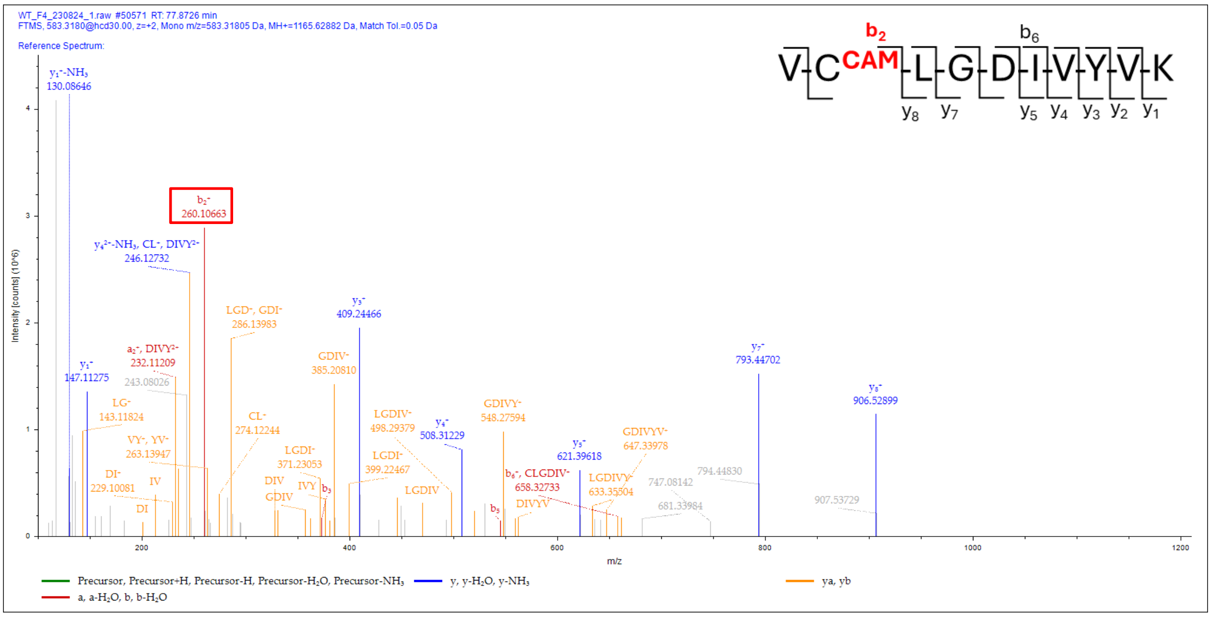


**Figure S15. MS/MS spectra of peptide with sequence VCLGDIVYVK belonging to cell division cycle protein 48 homologue.** (A) MS/MS of peptide of m/z 612.8123 (z=2) from *Δfh*. The b2 and y9 ions of m/z 319.09583 and 1125.54102 corresponding to 2SC are highlighted in red boxes. (B) MS/MS of peptide of m/z 583.318 (z=2) from *Δfh*. The b2 and y9 ions of m/z 260.10703 and 1066.55505 corresponding to Cys-CAM are highlighted in red boxes. (C) MS/MS of peptide of m/z 583.318 (z=2) from WT. The b2 ion of m/z 260.10663 corresponding to Cys-CAM is highlighted in a red box. The WT did not have the corresponding peptide with the cysteinyl residue succinated.

A

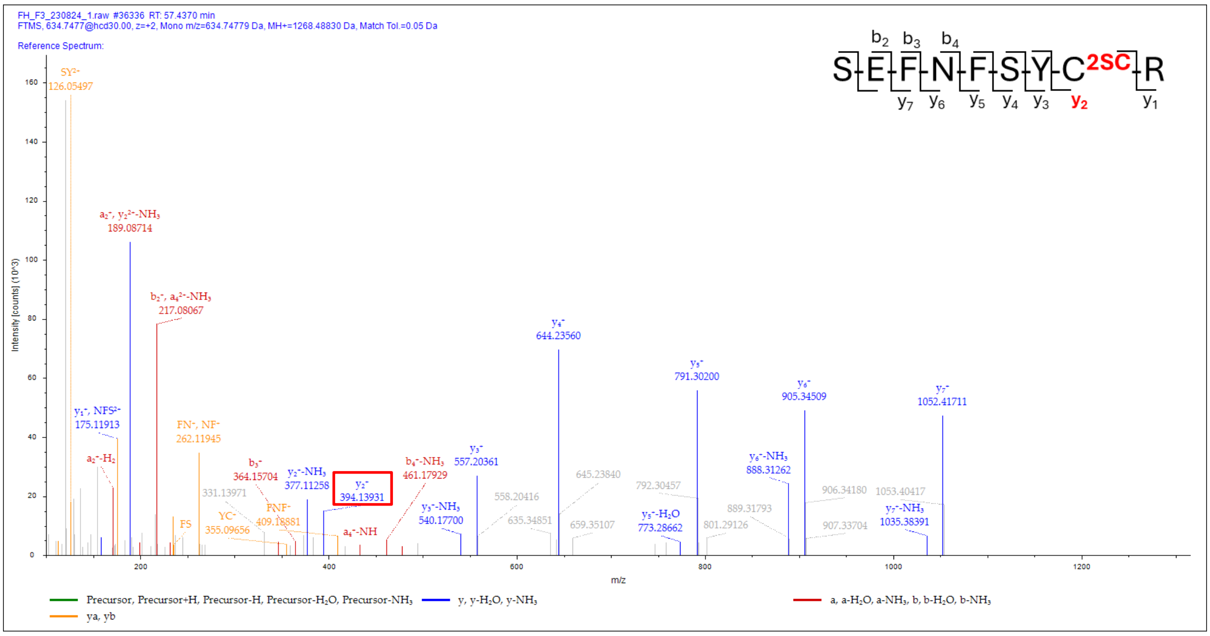

B

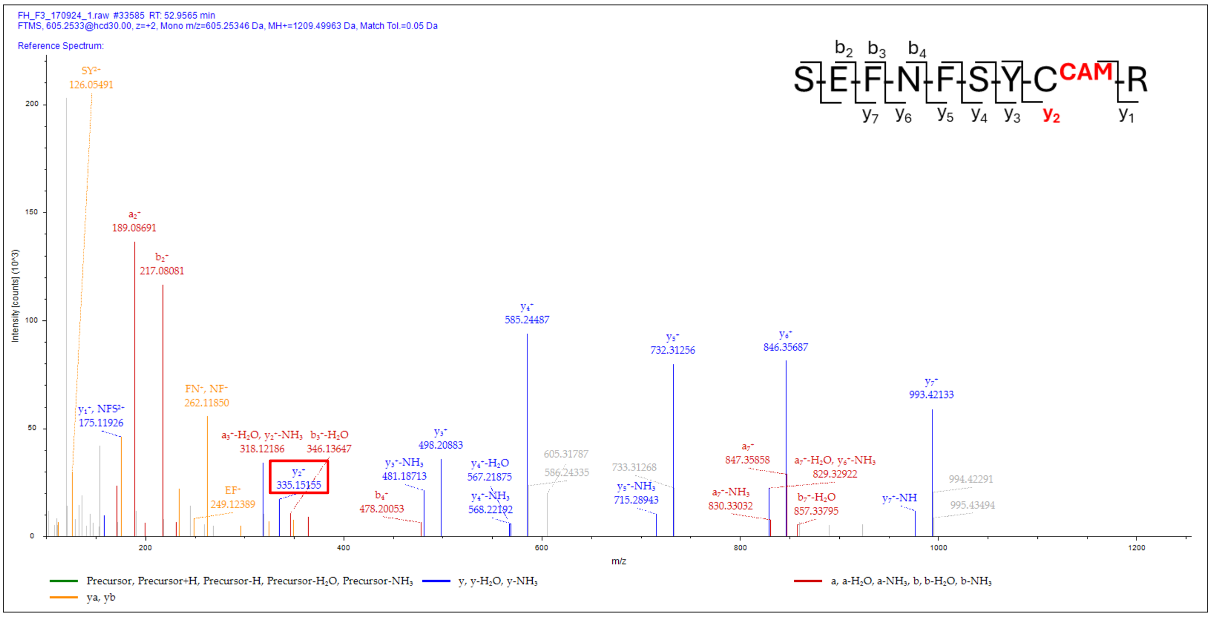


C

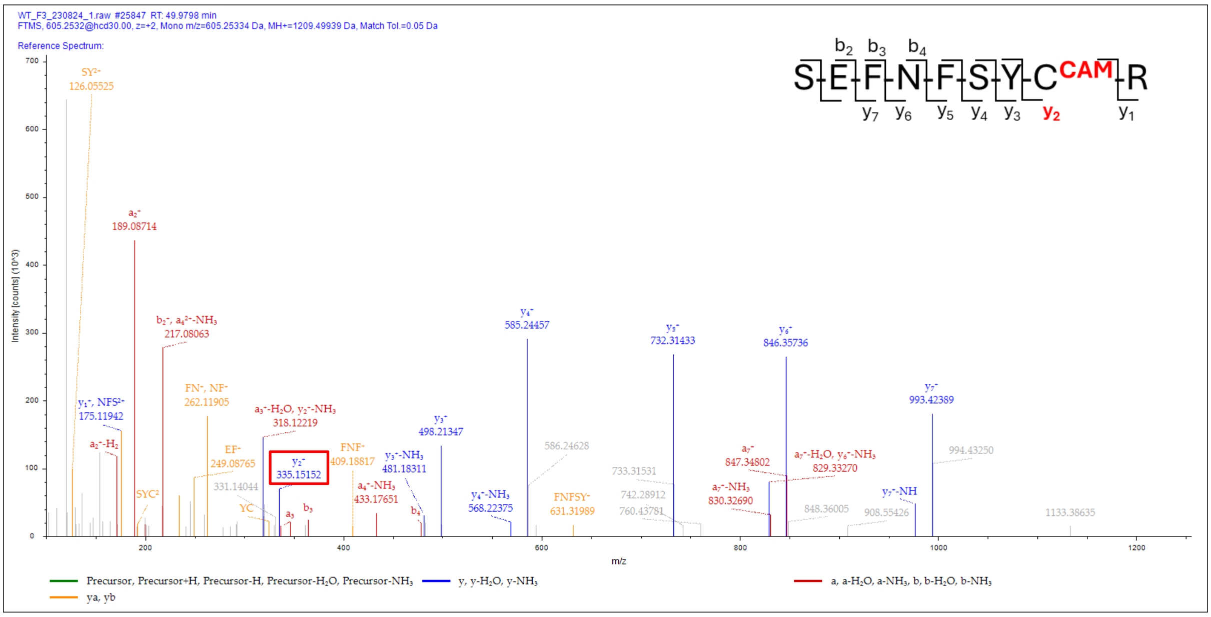


**Figure S16. MS/MS spectra of peptide with sequence SEFNFSYCR belonging to phosphoglucomutase.** (A) MS/MS of peptide of m/z 634.7477 (z=2) from *Δfh*. The y2 ion of m/z 394.13931 corresponding to 2SC is highlighted in a red box. (B) MS/MS of peptide of m/z 605.2533 (z=2) from *Δfh*. The y2 ion of m/z 335.15155 corresponding to Cys-CAM is highlighted in a red box. (C) MS/MS of peptide of m/z 605.2532 (z=2) from WT. The y2 ion of m/z 335.15152 corresponding to Cys-CAM is highlighted in a red box. The WT did not have the corresponding peptide with the cysteinyl residue succinated.

A

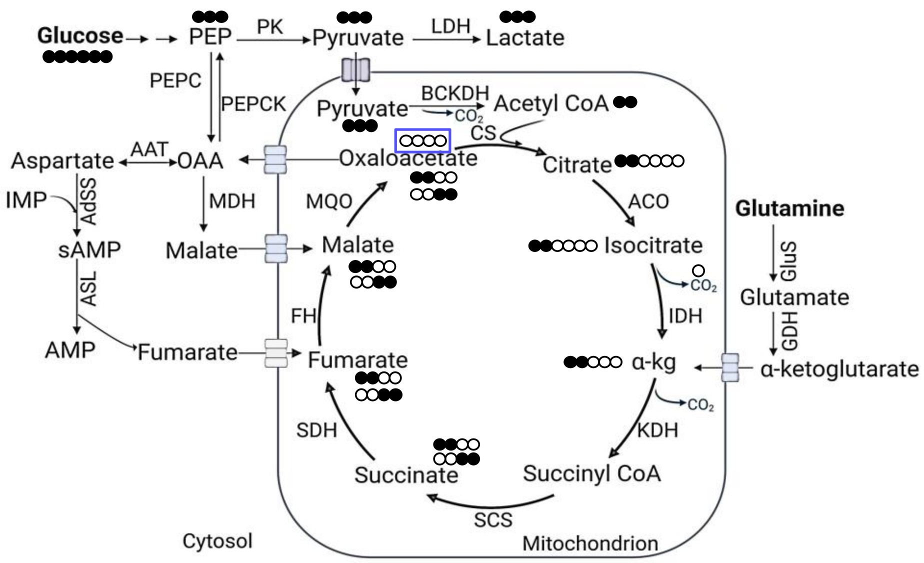
Glucose
PEP
PK
Pyruvate
LDH
Lactate
PEPC
PEPCK
Pyruvate
BCKDH
$CO_2$
Acetyl CoA
CS
Aspartate
AAT
OAA
Oxaloacetate
Citrate
IMP
AdSS
MDH
MQO
ACO
Glutamine
sAMP
Malate
Malate
Isocitrate
$CO_2$
GluS
Glutamate
ASL
FH
IDH
GDH
AMP
Fumarate
Fumarate
α-kg
α-ketoglutarate
$CO_2$
SDH
KDH
Succinate
Succinyl CoA
SCS
Cytosol
Mitochondrion

B

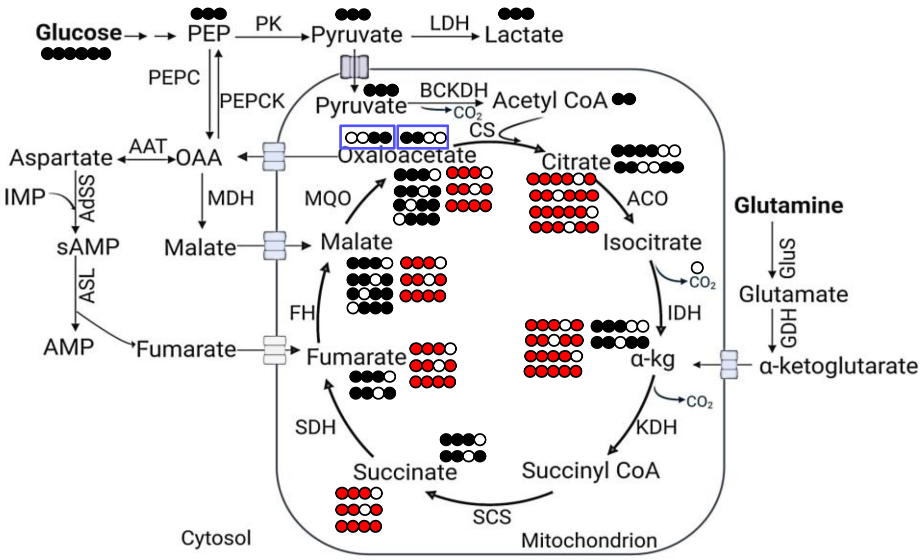
Glucose
PEP
PK
Pyruvate
LDH
Lactate
PEPC
PEPCK
Pyruvate
BCKDH
$CO_2$
Acetyl CoA
CS
Aspartate
AAT
OAA
Oxaloacetate
Citrate
IMP
AdSS
MDH
MQO
ACO
Glutamine
sAMP
Malate
Malate
Isocitrate
$CO_2$
GluS
Glutamate
ASL
FH
IDH
GDH
AMP
Fumarate
Fumarate
α-kg
α-ketoglutarate
$CO_2$
SDH
KDH
Succinate
Succinyl CoA
SCS
Cytosol
Mitochondrion

C

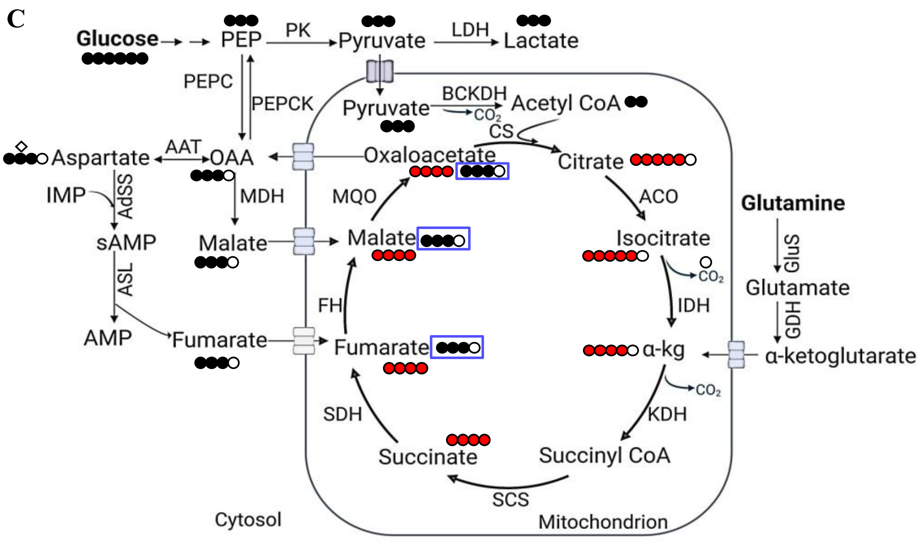


D

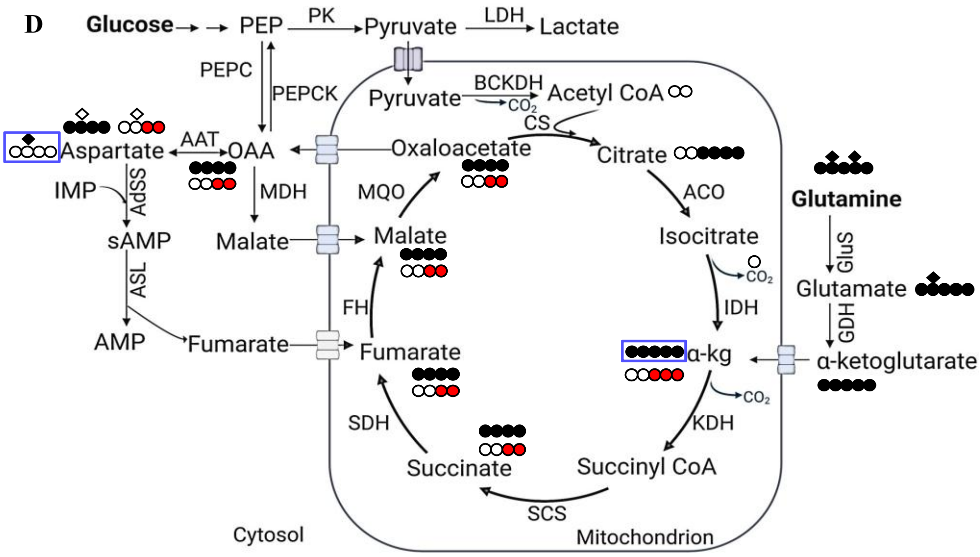


**Figure S17. Expected isotopologues of TCA cycle intermediates upon using (A-C) $^{13}C_6$-glucose and (D) $^{13}C_5{}^{15}N_2$-glutamine as tracer.** The schematics represent the expected labelling patterns in the metabolites of interest in the TCA cycle and associated pathways in the cytosol. The black and red circles represent $^{13}C$ carbon and white circles $^{12}C$ carbon. (A) The expected isotopologues from the first round of the TCA cycle. The canonical entry of pyruvate into the TCA cycle gives rise to M+2 acetyl-CoA in the mitochondrion, which upon condensing with unlabelled OAA (in blue box) gives rise to M+2 isotopologues of α-KG, succinate, fumarate, malate, and OAA. (B) The expected isotopologues from the second (black circles) and third round (red circles) of the TCA cycle. The M+2 labelled OAA from the first cycle (in blue box) condenses with M+2 acetyl-CoA and produces +3/+4 α-KG and +3 of succinate, fumarate, malate, and OAA in the 2$^{nd}$ round. The third round (shown as red circles) generates +4/+5 α-KG and +3/+4 of succinate, fumarate, malate, and OAA. After multiple rounds of the TCA cycle, the majority of α-KG will be +5 labelled and the downstream metabolites will be +4 labelled (18, 19). The labelling at different isotopomer positions arising from carbon scrambling is not shown for clarity in (B). It should be noted that OAA +2 or +3 condensing with acetyl-CoA M+0 can give rise to +1 or +2 isotopologues in metabolites of interest. However, as pyruvate achieves almost complete labelling within the time frame of incubation with $^{13}C_6$ glucose, the isotopic patterns arising from M+0 acetyl-CoA are not discussed. (C) The expected isotopologues from the anaplerosis of fumarate and malate. In the cytosol, M+3 malate and fumarate are downstream products arising from M+3 PEP. The M+3 malate and fumarate upon anaplerotic entry (shown as blue box) into the TCA cycle, give rise to M+3 OAA which upon condensing with M+2 acetyl-CoA gives rise to +4 of α-KG, succinate, fumarate, malate, and OAA (shown as red circles). (D) The expected isotopologues from the anaplerosis of α-KG. The diamond shape symbol represents nitrogen atoms of Gln, Glu, and Asp. M+5, N+2 glutamine gives rise to M+5, N+1 glutamate by loss of a molecule of ammonia. Labelled glutamate is converted to M+5 α-KG that anaplerotically feeds into the TCA cycle (shown as a blue box). In the first round, succinate, fumarate, and malate are M+4 labelled (black circles) and at the beginning of the second round, M+4 OAA condenses with acetyl-CoA, which subsequently generates, M+3 α-KG (red circles) (18). M+3 α-KG proceeds to yield M+2 succinate, fumarate, malate, and OAA (red circles). The labelled OAA (M+2 and M+4) from the TCA cycle exits the mitochondria and is converted to aspartate by the AAT reaction giving rise to M+2 or M+4 aspartate. At the same time, aspartate aminotransferase (AAT) catalyses the transfer of an amino group from glutamate to OAA, thus giving rise to different isotopologues of aspartate all having the N+1 nitrogen. The figure shows only M+0, N+1 aspartate for clarity (blue box). The schematic of the TCA cycle was generated using Biorender.

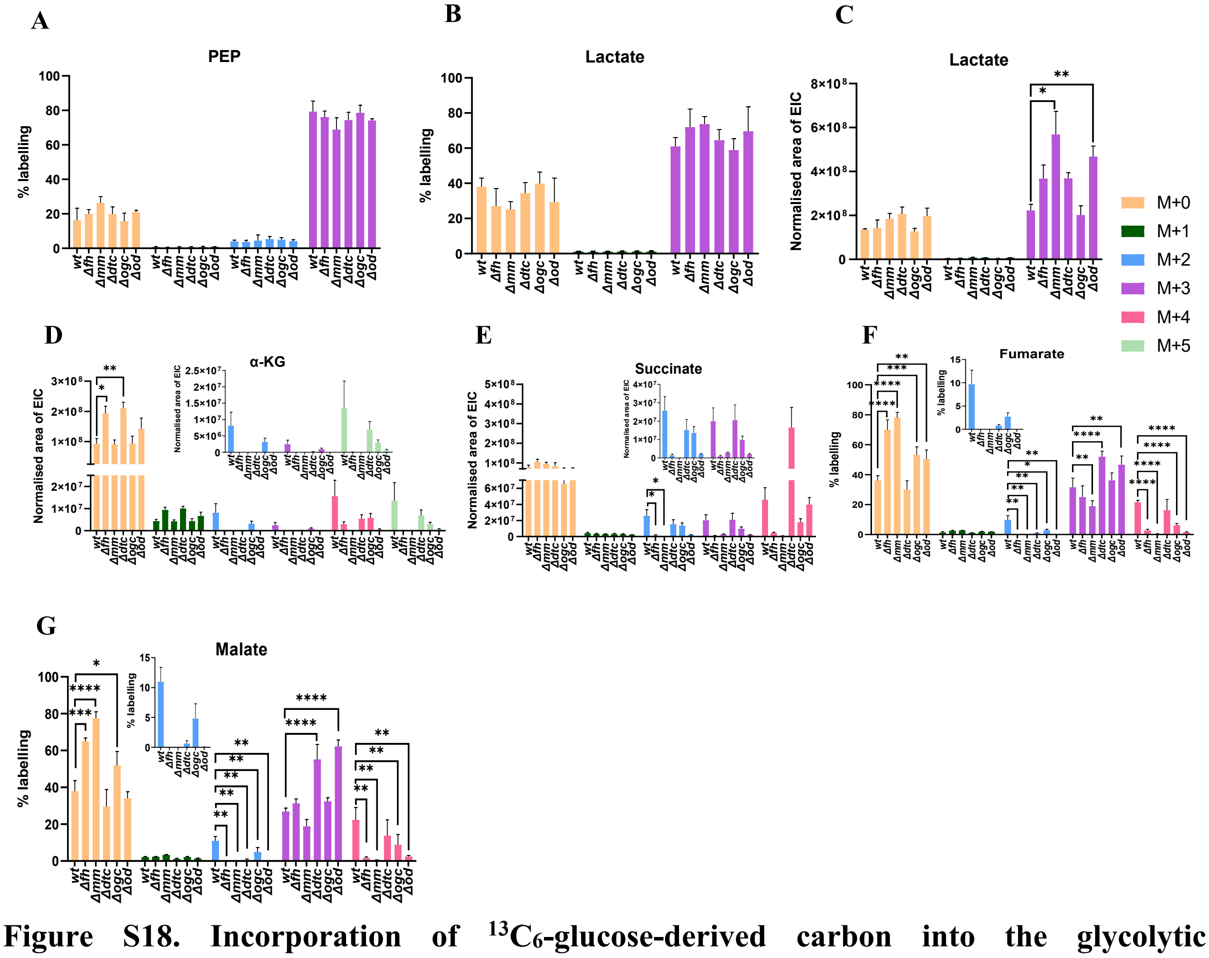


**Figure S18. Incorporation of $^{13}C_6$-glucose-derived carbon into the glycolytic intermediates, PEP and lactate and TCA cycle intermediates.** The bar graphs represent the percent labelling in the isotopologues of (A) PEP, (B) lactate, (F) fumarate, and (G) malate from the WT and knockout gametocytes. The mean values with error bars representing SD were plotted. The EIC area of the various isotopologues of (C) lactate, (D) α-KG, and (E) succinate in the WT and knockout gametocytes. Mean values with error bars representing SEM were plotted. The insets in D-G show the zoomed-in plots for some of the isotopologues. N=2 for wt, *Δfh*, *Δogc*, and *Δod* and N=3 for *Δmm* and *Δdtc* each with two or three technical replicates. Unpaired two-tailed t-tests were carried out using GraphPad Prism v9 ($p<0.05$ (*), $p<0.005$ (**), $p<0.001$ (***), $p<0.0001$ (****)).

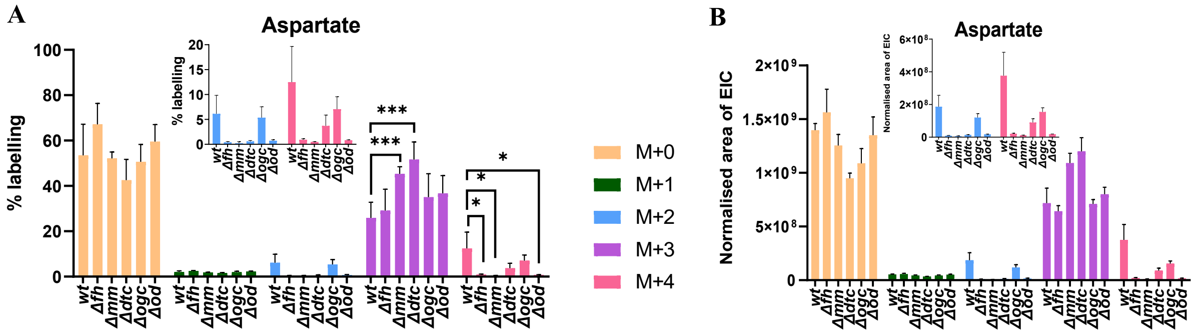


**Figure S19. Incorporation of $^{13}C_6$-glucose-derived carbon in aspartate.** (A) The bar graph represents the percent labelling in the isotopologues of aspartate from WT and knockout gametocytes. The mean values with error bars representing SD were plotted. (B) The EIC area of the various isotopologues of aspartate in the WT and knockout gametocytes represented as mean with SEM as error bars. The inset in (A) and (B) shows the zoomed-in plots for some of the isotopologues. N=2 for WT, *Δfh*, *Δogc*, and *Δod* and N=3 for *Δmm* and *Δdtc* each with two or three technical replicates. Unpaired two-tailed t-tests were carried out using GraphPad Prism v9 ($p<0.05$ (*), $p<0.005$ (**), $p<0.001$ (***), $p<0.0001$ (****)).

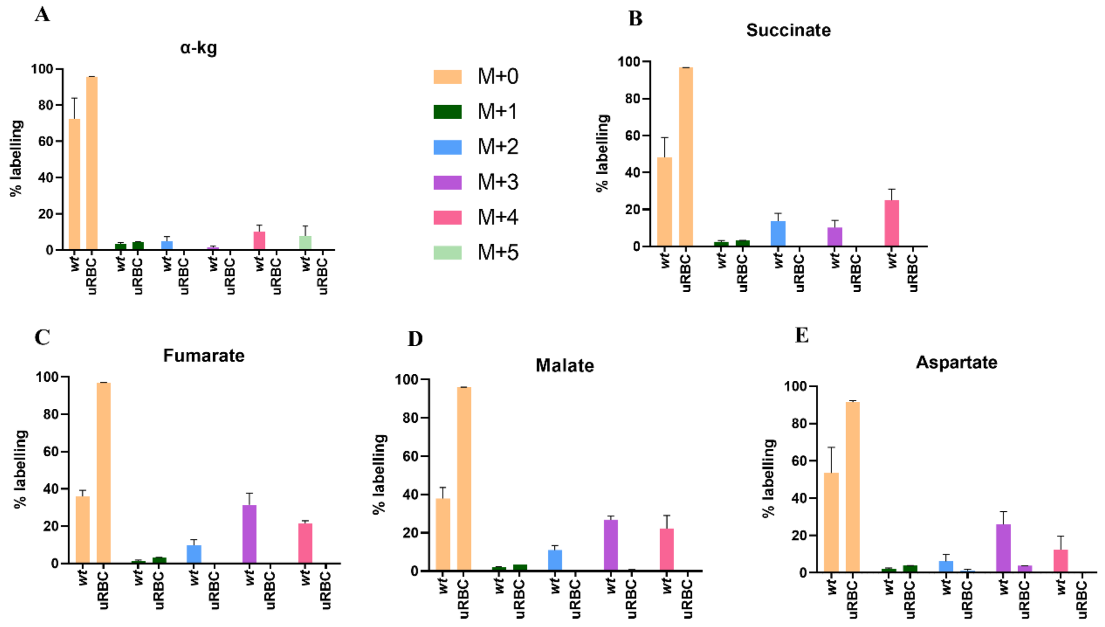


**Figure S20. Comparison of the level of incorporation of $^{13}C_6$-glucose-derived carbon into select metabolites across uninfected (uRBC) and *Pb* WT gametocyte-infected erythrocytes.** Label incorporation in (A) α-KG, (B) succinate, (C) fumarate, (D) malate, and

(E) aspartate. Mean values from three and four independent measurements for uRBC and *Pb* WT, respectively, with error bars representing SD are plotted.

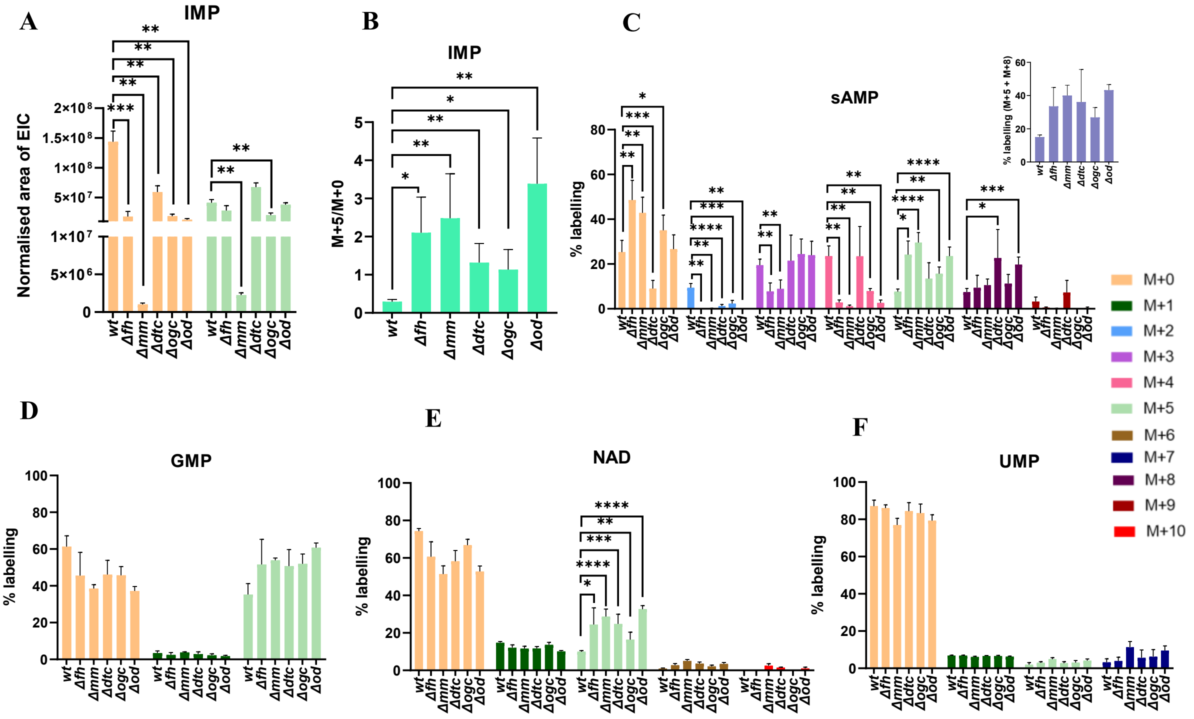


**Figure S21. Incorporation of $^{13}C_6$-glucose-derived carbon into nucleotides.** (A) The EIC area of various isotopologues of IMP in the WT and knockout gametocytes. Mean values with error bars representing SEM were plotted. (B) Ratio of EIC area of M+5/M+0 isotopologues of IMP in the WT and knockouts. The percent labelling in the isotopologues of (C) sAMP, inset shows sum of labelling in M+5 and M+8 isotopologues, (D) GMP, (E) NAD, and (F) UMP from WT and knockout gametocytes. Mean values with error bars representing SD were plotted. N=2 for WT, *Δfh*, *Δogc*, and *Δod* and N=3 for *Δmm* and *Δdtc* each with two or three technical replicates. Unpaired two-tailed t-tests were carried out using GraphPad Prism v9 ($p<0.05$ (*), $p<0.005$ (**), $p<0.001$ (***), $p<0.0001$ (****)).

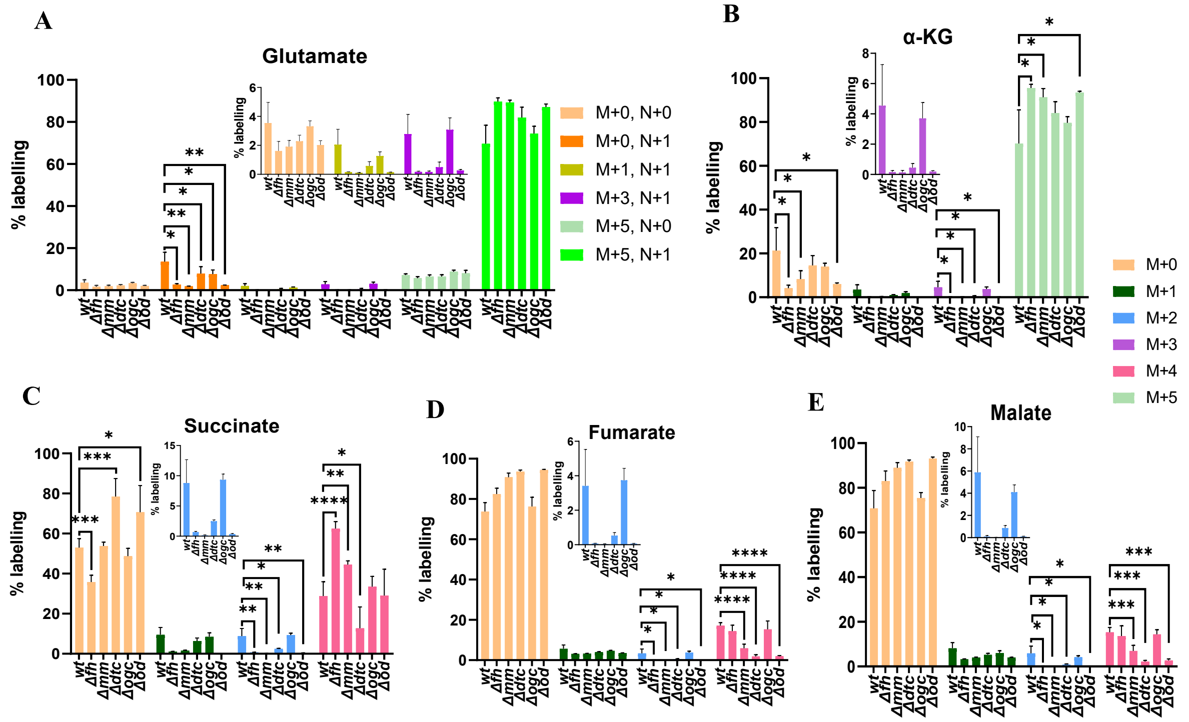


**Figure S22. Incorporation of $^{13}C_5{}^{15}N_2$-glutamine-derived carbon into glutamate and the TCA cycle intermediates**. The bar graphs represent the percent enrichment in the isotopologues of (A) glutamate, (B) α-KG, (C) succinate, (D) fumarate, and (E) malate from WT and knockout gametocytes. Mean values with error bars representing SD are plotted. The inset in all panels are zoomed-in plots for some of the isotopologues. N=2 for WT and knockouts with two to three technical replicates each. Unpaired two-tailed t-tests were carried out using GraphPad Prism v9; p<0.05 (*), p<0.005 (**), p<0.001 (***), p<0.0001 (****).

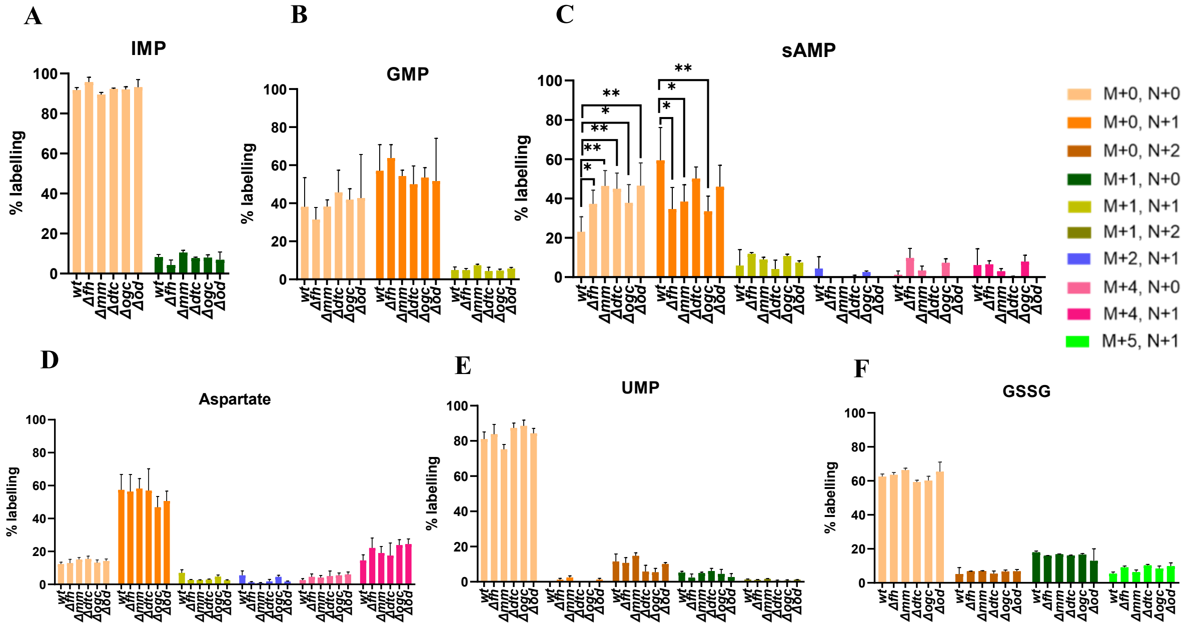


**Figure S23. Incorporation of $^{13}C_5{}^{15}N_2$-glutamine-derived carbon or nitrogen into aspartate and nucleotides.** The bar graphs represent the percent labelling in the isotopologues of (A) IMP, (B) GMP, (C) sAMP, (D) aspartate, (E) UMP, and (F) GSSG from WT and knockout gametocytes. Mean values with error bars representing SD were plotted. N=2 for WT and knockouts with two to three technical replicates each. Unpaired two-tailed t-tests were carried out using GraphPad Prism v9 ($p<0.05$ (*), $p<0.005$ (**), $p<0.001$ (***), $p<0.0001$(****)).

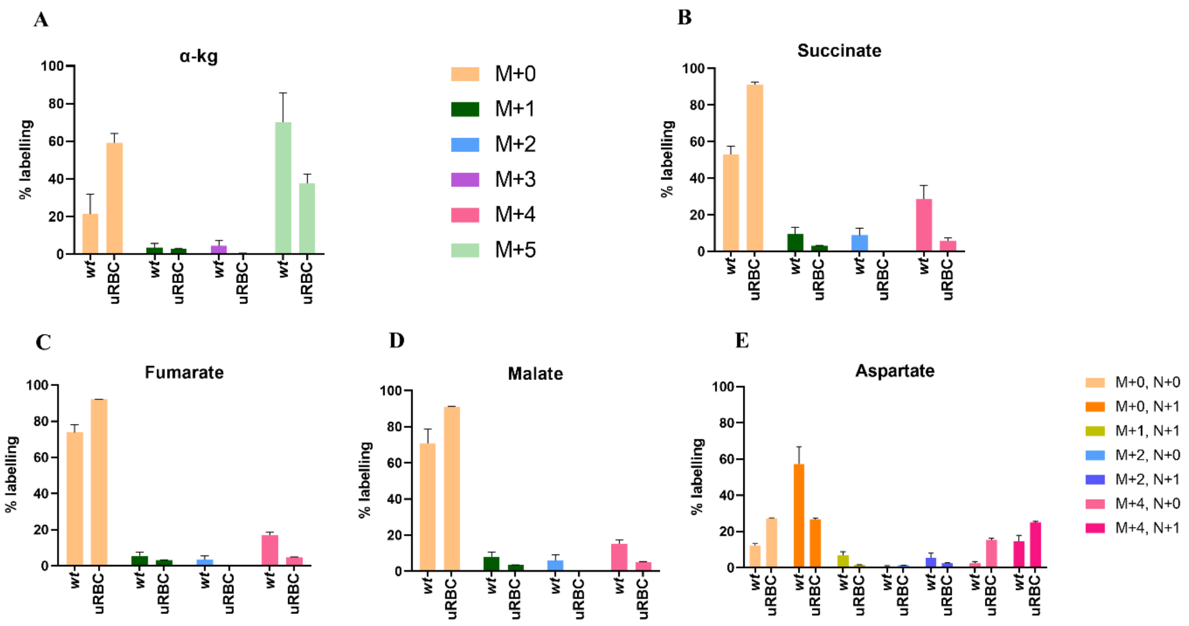


**Figure S24. Comparison of the level of incorporation of $^{13}C_5{}^{15}N_2$-glutamine-derived carbon or nitrogen into select metabolites across uninfected (uRBC) and *Pb* WT gametocyte-infected erythrocytes.** Label incorporation into (A) α-KG, (B) succinate, (C) fumarate, (D) malate, and (E) aspartate. Mean values from three and five independent measurements for uRBC and *Pb* WT, respectively, with error bars representing SD are plotted. The color key for the bars used in panels (A) to (D) is provided on top while the color key for panel (E) is placed adjacent to it.

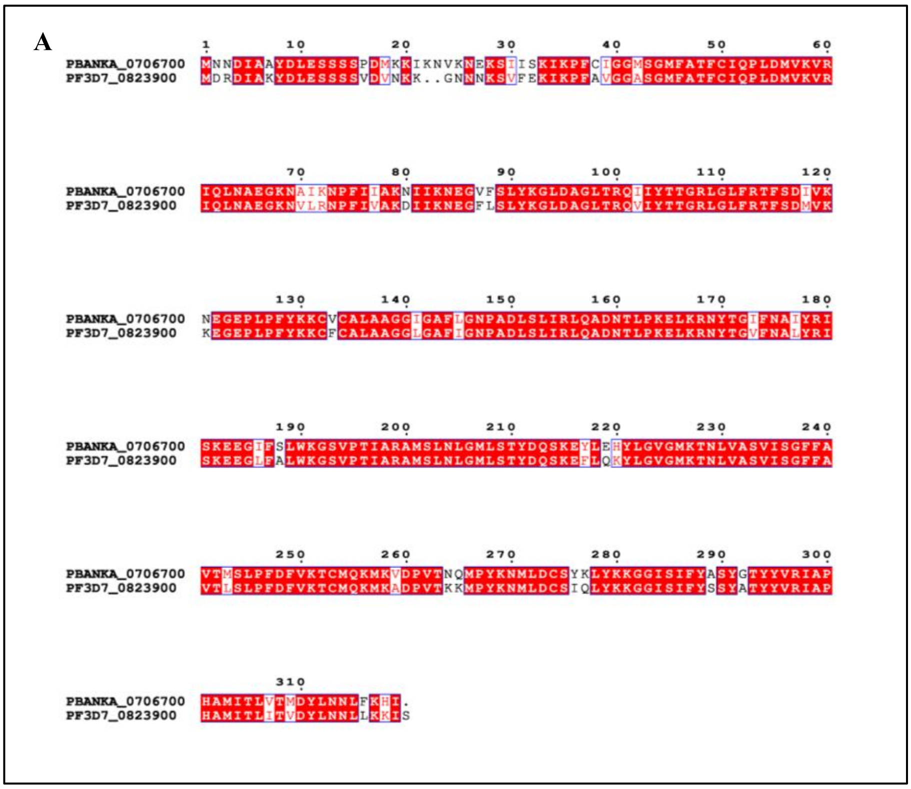

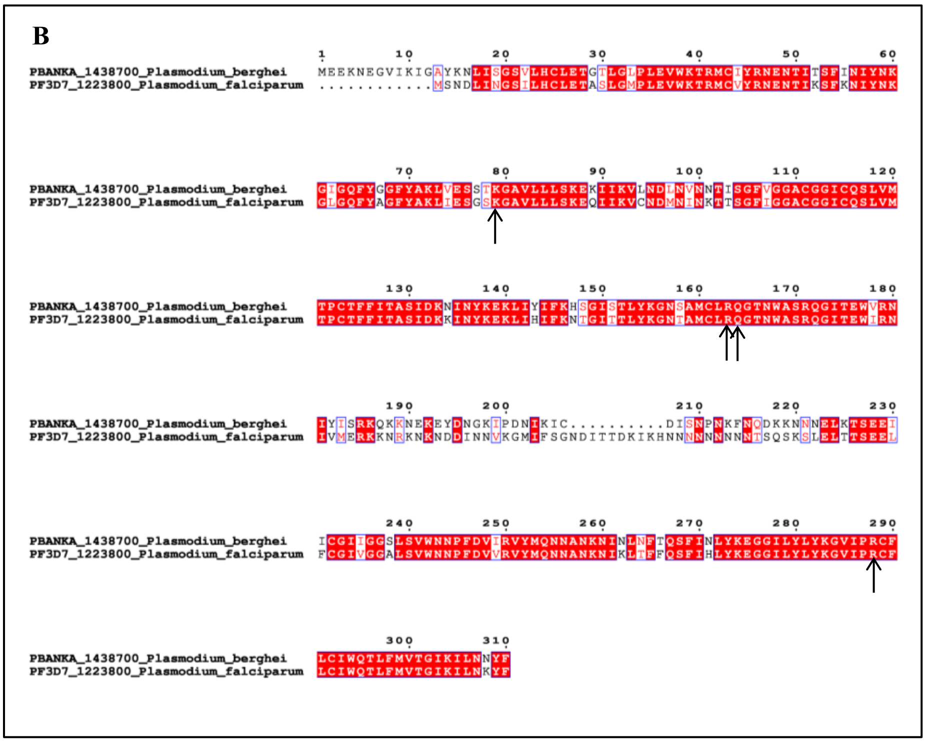

**Figure S25. Sequence alignment** of (A) DTC and (B) OGC sequences. PBANKA_0706700, PbDTC; Pf3D7_0823900, PfDTC; PBANKA_1438700, PbOGC; Pf3D7_1223800, PfOGC. The residues in PfOGC involved in substrate binding as proposed earlier (20) are indicated by black arrows.

**Table S1. Comparison of metabolite levels across PbWT gametocytes and uninfected RBC (uRBC).**

| Metabolites | **PbWT[a]** | | **uRBC[b]** | | **Fold change** |
|---|---|---|---|---|---|
| | Average | Std dev | Average | Std dev | PbWT/uRBC |
| Pyruvate | 3.68E+07 | 9.06E+06 | 2.20E+06 | 6.58E+05 | 16.67 |
| PEP | 1.76E+07 | 5.30E+06 | 1.26E+06 | 9.17E+04 | 13.98 |
| Lactate | 4.12E+08 | 1.62E+08 | 9.29E+06 | 1.63E+06 | 44.38 |
| α-KG | 1.08E+08 | 1.80E+07 | 6.59E+05 | 2.84E+05 | 164.31 |
| Succinate | 1.34E+08 | 8.27E+07 | 1.80E+06 | 3.41E+04 | 74.65 |
| Fumarate | 1.09E+08 | 7.37E+07 | 1.56E+06 | 2.11E+05 | 69.94 |
| Malate | 3.40E+08 | 1.40E+08 | 2.05E+07 | 3.37E+06 | 16.56 |
| Aspartate | 2.45E+09 | 4.13E+08 | 1.23E+08 | 1.37E+07 | 19.90 |
| Glutamine | 1.41E+09 | 3.38E+08 | 4.32E+08 | 6.57E+06 | 3.25 |
| Glutamate | 9.96E+09 | 2.48E+09 | 8.60E+07 | 2.35E+07 | 115.82 |
| Alanine | 2.40E+07 | 8.51E+06 | 5.04E+06 | 9.01E+05 | 4.76 |
| GSSG | 8.61E+08 | 2.30E+08 | 3.04E+07 | 1.17E+06 | 28.29 |
| Succinyl-GSH | 3.05E+07 | 1.61E+07 | 1.33E+05 | 8.15E+03 | 229.10 |
| Lactoyl-GSH | 2.69E+07 | 9.27E+06 | 3.70E+05 | 4.42E+04 | 72.50 |
| 2SC | 1.60E+06 | 9.38E+05 | 1.63E+05 | 8.80E+03 | 9.83 |
| UMP | 2.68E+08 | 8.73E+07 | 6.79E+04 | 6.30E+04 | 3943.43 |
| NAD | 2.46E+08 | 9.78E+07 | 2.14E+06 | 2.24E+05 | 114.97 |
| GMP | 9.07E+07 | 2.86E+07 | 3.60E+05 | 9.23E+04 | 251.91 |
| IMP | 1.62E+08 | 9.32E+07 | 2.36E+05 | 5.60E+04 | 686.96 |
| Hypoxanthine | 7.46E+08 | 3.87E+08 | 6.50E+06 | 6.60E+05 | 114.92 |
| Adenine | 1.96E+08 | 5.05E+07 | 2.03E+06 | 5.36E+05 | 96.45 |

[a]Erythrocytes infected with PbWT gametocytes. The EIC area corresponding to $3x10^7$ cells is listed. [b]Uninfected RBC (URBC). The EIC area corresponding to $1.5x10^6$ cells (5% of $3x10^7$) is listed.

**Table S2. List of primers along with their sequences (5′-3′)**

| | |
|---|---|
| P1 | GCCTTCAAAATTTAAATTTATTTTAATATTTCC |
| P2 | CAGCGGTTTCTTTACCAGACTCGAGTTATTTGTATAGTTCATCCATGCC |
| P3 | ATGCTGAATATTAAGAATATAGTTCGAAGGATAAATAG |
| P4 | TTACAAATAATTGACTGGATAATCCCCTTGATAGAG |
| P5 | CCTAACGTGCTTGTATTATCATCATCCTCG |
| P6 | ACTTCTTAAACCTAATCTGTAGTAAGGAAGGGATTG |
| P7 | GTGGATGAAAATATTACTGGTGCTTTGAGGGGTGAGC |
| P8 | CTTGGAAATGTCATCATTTCCTTTGCTTTCTGATTC |
| P9 | GCAATTTGGTAAACTGTTCTGTGTTATAATAACAAAGTGTAAAG |
| P10 | CACGTTTAACATTTTCATGTTATTTCAAAGTGTATCATG |
| P11 | CATCTAGACCTCTGTCGAGGGCACCAGTAACAATTCTG |
| P12 | CCACAACCTCTTCAGTAGAAGGTAAACAGAATCTGGTG |
| P13 | ATGCCAAAAATATCATTAATAGGAAGTGG |
| P14 | TTAGTCAATTACATTCAAGGCTTTTTGTGTATTTTCTC |
| P15 | GAATAATCTTAAAGTGTAGAAATTGTGAACATTGCAG |
| P16 | GGCAAGCAAGTGGGATCAGAAGGGTATGGACATTGC |
| P17 | GTGGATGAAAATATTACTGGTGCTTTGAGGGGTGAGC |
| P18 | CATAACTGCAACATTAATTCATCCTAATATATGGAGTTTG |
| P19 | CTATATGTGTTTGAAAAGATTGTTAAGATAATCCATGGTAAC |
| P20 | ATGAATAATGATATAGCGGCATATGACCTTGAATC |
| P21 | GACTGTGATCAGAAGTTCCGAGATATAACAAAAGC |
| P22 | ACTTCTTAAACCTAATCTGTAGTAAGGAAGGGATTG |
| P23 | GTGGATGAAAATATTACTGGTGCTTTGAGGGGTGAGC |
| P24 | CACCATCGTTGATATCATAATACAGTGCACAC |
| P25 | CCATACATCGATTTTTGTGATTTTGTTTTAATGAAGG |
| P26 | GGTATATTGAAAAGTAAGAAGAAAAAAAACATCATAACTTTGTAAGC ATTG |
| P27 | ACACCTTCAAAGAAACACGCCA |
| P28 | ACCTCCACATGCCCCACCTACA |
| P29 | GGCAAGCAAGTGGGATCAGAAGGGTATGGACATTGC |
| P30 | AGCATGCTGAGCATCATAGCCT |
| P31 | GTACTTAATGCCTTTCTCCTCCTGGACATCAGAG |
| P32 | GCTGCTTTGCAAAAAATGTCATCC |
| P33 | TTTATAAAACATAGGGGGATCCATGGAAGAAAAAAATGAAGGGG |
| P34 | AAAAGTTCTTCTCCTTTACTCATTCCTCCTGATCCTCCTCCTGATCCTC CAAAATAGTTATTTAATATCTTAATTCC |
